# The *Escherichia coli* Transcriptome Mostly Consists of Independently Regulated Modules

**DOI:** 10.1101/620799

**Authors:** Anand V. Sastry, Ye Gao, Richard Szubin, Ying Hefner, Sibei Xu, Donghyuk Kim, Kumari Sonal Choudhary, Laurence Yang, Zachary A. King, Bernhard O. Palsson

**Affiliations:** Department of Bioengineering, University of California San Diego, La Jolla, CA 92093, USA; Department of Biological Sciences, University of California San Diego, La Jolla, CA 92093, USA; Department of Pediatrics, University of California San Diego, La Jolla, CA 92093, USA; Novo Nordisk Foundation Center for Biosustainability, 2800 Kongens Lyngby, Denmark

## Abstract

Underlying cellular responses is a transcriptional regulatory network (TRN) that modulates gene expression. A useful description of the TRN would decompose the transcriptome into targeted effects of individual transcriptional regulators. Here, we applied unsupervised learning to a compendium of high-quality *Escherichia coli* RNA-seq datasets to identify 70 statistically independent signals that modulate the expression of specific gene sets. We show that 50 of these transcriptomic signals represent the effects of currently characterized transcriptional regulators. Condition-specific activation of signals was validated by exposure of *E. coli* to new environmental conditions. The resulting decomposition of the transcriptome provided: (1) a mechanistic, systems-level, network-based explanation of responses to environmental and genetic perturbations, (2) a guide to gene and regulator function discovery, and (3) a basis for characterizing transcriptomic differences in multiple strains. Taken together, our results show that signal summation forms an underlying principle that describes the composition of a model prokaryotic transcriptome.

## Main

The transcriptional regulatory network (TRN) senses and integrates complex environmental and intracellular information to coordinate gene expression of a cell. Reverse engineering the TRN informs how an organism responds to diverse stresses and unfamiliar environments^1–3^. A fully characterized TRN would enable the prediction and mechanistic explanation of an organism’s dynamic adaptation to environmental or genetic perturbations.

Reconstruction of a genome-scale TRN requires a substantial number of experiments to integrate the binding sites for each regulator and characterize their activities^4,5^. Unlike eukaryotic TRNs, which contain highly-connected co-associations^6^, prokaryotic TRNs exhibit a simpler structure; over 75% of genes in the model bacteria *Escherichia coli* are known targets of two or fewer TFs^7^ (Fig. S1a).

The TRN structure is encoded in the genome as regulator binding sites and is invariant to environmental dynamics. However, environmental and genetic perturbations alter the activity states of transcriptional regulators to change their DNA binding affinity, which in turn modulates the transcriptome in a condition-specific manner. Thus, a measured expression profile reflects a combination of the activity of all transcriptional regulators under the examined condition, posing a fundamental deconvolution challenge.

Compendia of expression profiles have been leveraged to infer TRNs by identifying shared patterns across gene expression profiles, rather than using direct DNA-TF binding information^8,9^. Many inference methods define groups of genes, or modules, with similar expression profiles that are often functionally related or co-expressed. A recent review showed that independent component analysis (ICA), a signal deconvolution algorithm, outperformed most other module detection algorithms in identifying groups of coregulated genes^10^.

ICA is a blind source separation algorithm used to deconvolute mixed signals into their individual sources and determine their relative strengths^11^. Prior application of ICA to microarray expression data^12^ has identified co-expressed, functionally-related gene sets^13–15^ that often map to metabolic pathways^16,17^. The overall expression levels, or activities, of the gene sets have been leveraged to classify tumor samples^18,19^ and connect transcriptional modules to disease states^20^.

A current challenge for analyzing transcriptional regulation is to separate the condition-invariant network structure from its condition-dependent expression state on a genome scale. Here, we overcome this limitation for the *E. coli* TRN by simultaneously extracting its structure and regulator activities from a transcriptomics compendium. This approach relies on: (1) the availability of high-quality, self-consistent, and condition-rich expression profiling datasets; (2) the use of ICA to concurrently identify regulator targets and activities; and (3) validation through the association of inferred regulator targets with observed molecular interactions. The elucidated TRN structure deconvolutes transcriptomic responses of *E. coli* into a summation of condition-specific effects of individual transcriptional regulators.

## Results

### Independent Component Analysis (ICA) Extracts Regulatory Signals from Expression Data

In order to extract regulatory interactions from expression data, diverse conditions must be profiled to discriminate between the effects of transcriptional regulators. Previous studies have compiled transcriptomics data from independent research groups to study the transcriptional states and regulation of *E. coli*^*21–24*^. Even after resolving the significant normalization challenge with such disparate datasets, many sources of variation remain that obscure biological signals^25–27^. These datasets mostly contain microarray data; RNA sequencing (RNA-seq) data yields higher quality data with less noise and larger dynamic range^28^.

We therefore compiled PRECISE, a Precision RNA Expression Compendium for Independent Signal Exploration. This high-fidelity expression profile compendium (median R^2^ = 0.98 between biological replicates, see Fig. S1b) comprises 190 RNA-seq datasets across 113 unique experimental conditions of *Escherichia coli* K-12 MG1655 and BW25113 generated from a single laboratory and obtained using a standardized protocol (Supplemental Dataset 1). PRECISE accounts for over 15% of publicly available RNA-seq data in NCBI GEO^29^ for *E. coli* K-12 MG1655 and BW25113 (Fig. S1c).

We applied ICA to identify independent sources of variation in gene expression in PRECISE. The traditional use of ICA as a signal decomposition algorithm is illustrated in Fig. 1a. When applied to transcriptomics data, ICA decomposes a collection of expression profiles (**X**) into (1) a set of components, which represent underlying biological signals (**S**), and (2) the components’ condition-specific activities (**A**) (Fig. 1b,c). Each component, represented by a column of **S**, contains a coefficient for each gene that represents the effect of a particular underlying signal on the gene’s expression level. Components do not contain information on the condition-specific transcriptomic state. Conversely, ICA computes activity levels for each component across every condition in the dataset, represented by a row of **A**, to account for condition-dependent expression changes (Fig. 1d). Each expression profile is represented by the summation over all components, each scaled by its condition-specific activity (Fig. 1e).

**Fig. 1:**
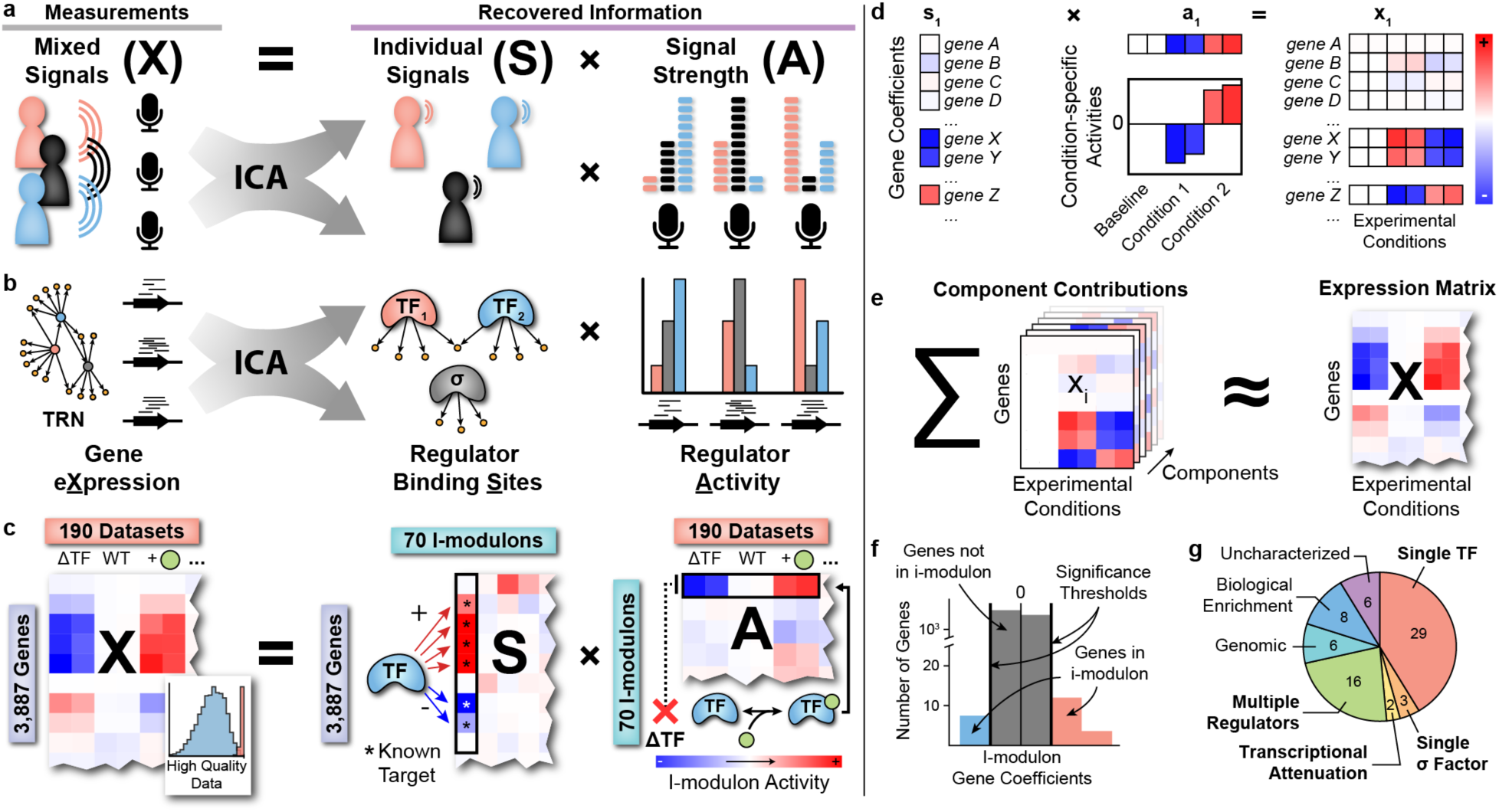
Independent Component Analysis (ICA) Extracts Regulatory Signals from Expression Data. (a) Given three microphones recording three people speaking simultaneously, each microphone records each voice (i.e. signal) at different volumes (i.e. signal strengths) based on their relative distances. Using only these measured mixed signals, ICA recovers the original signals and their relative signal strengths by maximizing the statistical independence of the recovered signals11,53. The mixed signals (**X**) are a linear combination of the matrix of recovered source signals (**S**) and the mixing matrix (**A**) that represents the relative strength of each source signal in the mixed output signals. This relationship is mathematically described as **X** = **SA**. (b) An expression profile under a specific condition can be likened to a microphone in a cell, measuring the combined effects of all transcriptional regulators. (c) Schematic illustration of ICA applied to a gene expression compendium. See Fig. S1b for additional details on data quality. The example TF is a dual regulator that primarily upregulates genes, and is activated by the green circular metabolite. Example experimental conditions shown are a TF knock-out, wild-type, and wild-type grown on medium supplemented with the activating metabolite. Each column of **X** contains an individual expression profile across 3,887 genes in *E. coli*. (d) Each component (column of **S**) contains a coefficient for each gene. These are scaled by the component’s condition-specific activities (row in **A**) to form a single layer of the transcriptomic compendium (e) The sum of all 70 layers, or components, reconstructs most of the variance in the original compendium. (f) Distribution of i-modulon categories. Categories of regulatory i-modulons are labelled in bold font. For more details, see Fig. S1 and Table 1.

ICA of PRECISE produced 70 robust components that explained 80% of the expression variation (Fig. S1d). Most gene coefficients in a component were near zero, indicating that each component affects a small number of significant genes (Fig. 1f). We removed genes with coefficients below a threshold, resulting in a set of significant genes for each component. We defined these condition-invariant sets of genes as “i-modulons”, since these genes were **i**ndependently **modul**ated at constant ratios across every condition in the database. We note that a gene may appear in multiple i-modulons if its expression is dependent on multiple underlying biological signals (Fig. S1e).

**Table 1:**
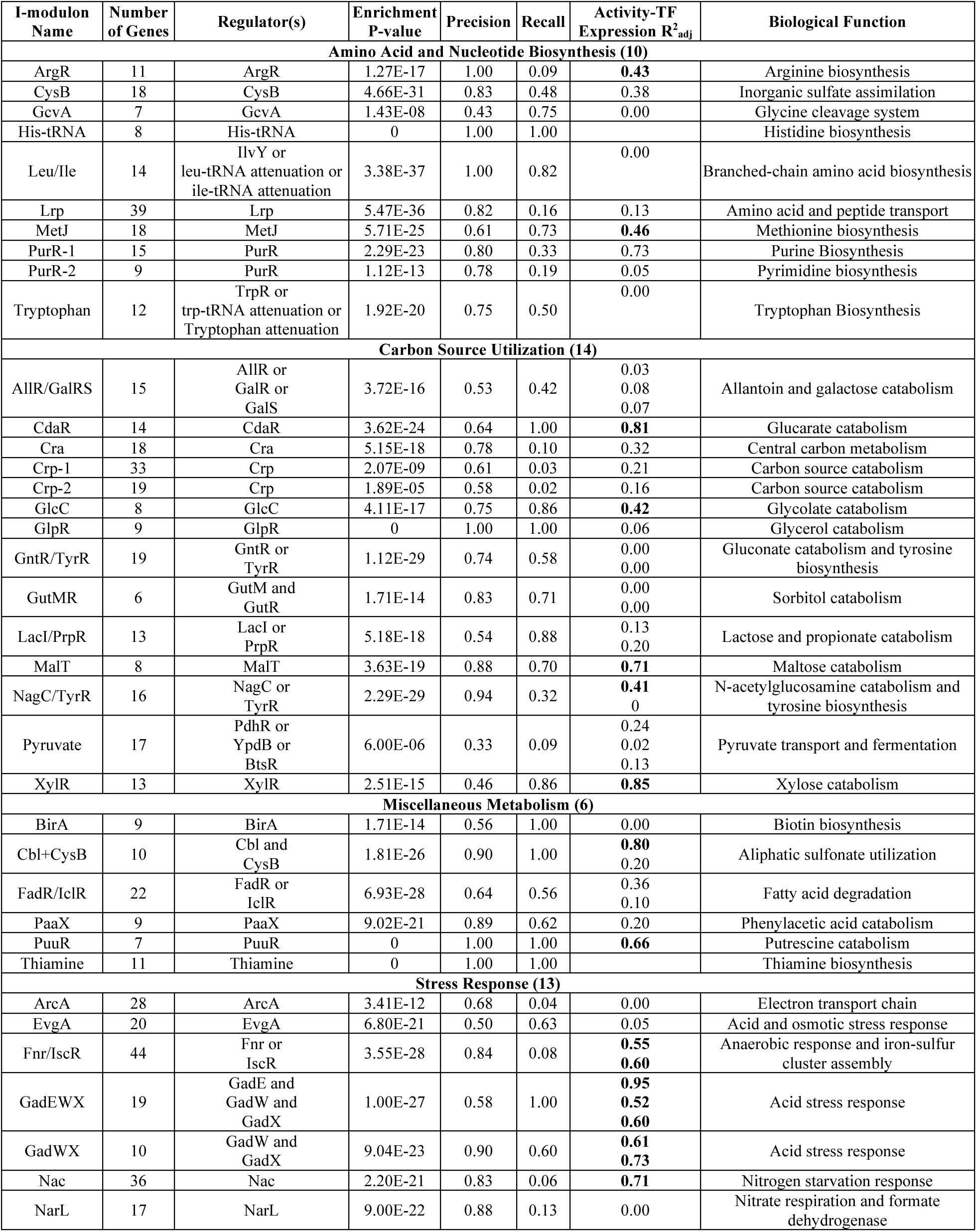
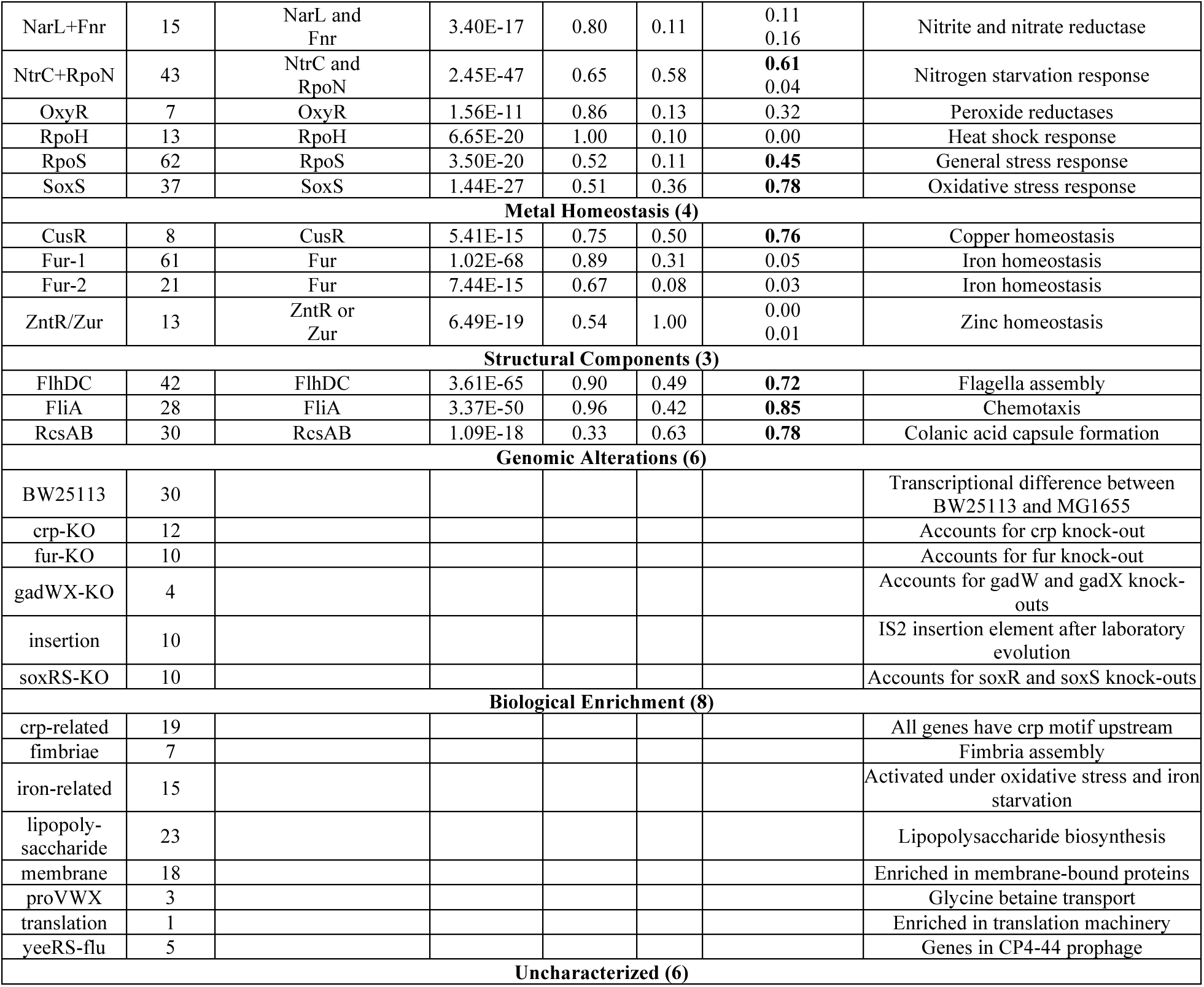
Overview of i-modulons derived from PRECISE. I-modulons are either named after their associated regulator(s) or their biological function. Strong activity-TF relationships (R^2^_adj_ > 0.4) are marked in bold.

The 70 resulting i-modulons are listed in Table 1. We hypothesized that each i-modulon was controlled by a particular transcriptional regulator, and that the i-modulon activity represented the condition-dependent activation state of the corresponding transcriptional regulator. To test this hypothesis, we examined the consistency between i-modulons and reported regulons, defined as the set of genes targeted by a common regulator, using a database of over 7,000 experimentally-derived regulatory interactions for *E. coli* ^4^ (Fig. S1f).

We identified significant overlaps between regulons and 50 of the 70 i-modulons. We defined these 50 i-modulons as “regulatory i-modulons”. Three regulatory i-modulons were linked to a single sigma factor (*rpoS, rpoH, fliA*), four were linked to transcriptional attenuation (including the thiamine riboswitch)^30–32^, and 45 were linked to TFs. Sixteen regulatory i-modulons were associated with multiple regulators, as described in Table 1. Of the 20 non-regulatory i-modulons, six i-modulons were associated with distinct genetic changes, such as gene knock-outs and strain-specific differences, and eight of the remaining i-modulons were enriched in a specific biological function or process (Fig. 1g).

I-modulons were labeled by their associated regulator (e.g., the MetJ i-modulon) or biological function (e.g., the Pyruvate i-modulon). The majority of regulatory i-modulons (30 of 50) mapped to metabolic pathways (Fig. S2a), as previously noted^16^. The remaining regulatory i-modulons represented diverse cellular responses (Fig. S2b,c). Detailed information for selected i-modulons, including gene composition, regulon enrichments, activity levels, and upstream regulator binding motifs, is available in Supplemental Dataset 2.

### Validation of I-modulon-Regulator Relationships

On average, 75% of genes in a regulatory i-modulon were reported targets of the linked transcriptional regulator(s) (Fig. 2a,b). This precision was significantly higher than the precision observed for other sparse decomposition methods and ICA applied to microarray datasets (Fig. S3a,b). We hypothesized that the remaining genes in each i-modulon were actually regulated by the associated regulator, but the binding sites were not experimentally determined. We tested this claim by performing ChIP-exo^33^ to locate binding sites for the TFs MetJ and CysB (Tables S1,2), which regulate methionine biosynthesis and sulfate assimilation, respectively.

**Fig. 2:**
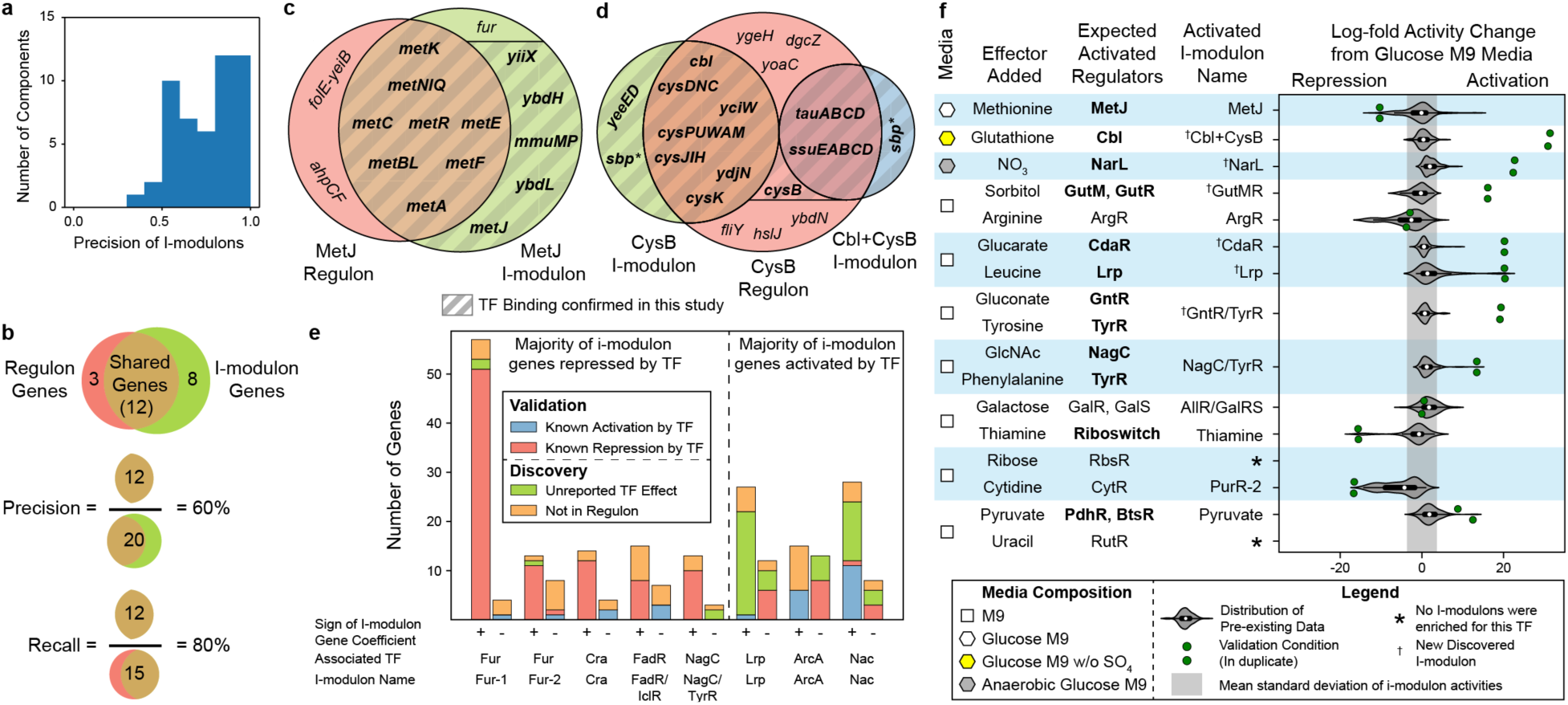
Validation of I-modulon-Regulator Relationships. (a) Precision across all 50 regulatory i-modulons. (b) Schematic illustration defining precision and recall for i-modulons. Precision is the fraction of genes in an i-modulon that are in the linked regulon, and recall is the fraction of genes in a regulon that are in the linked i-modulon (c) Comparison of genes in the MetJ regulon (red) and i-modulon (green). Genes validated by ChIP-exo are shown in bold. (d) Comparison of genes in the CysB regulon (red) and the CysB and Cbl+CysB i-modulons (green and blue, respectively). Most genes in the Cbl+CysB i-modulon were regulated by both Cbl and CysB. The starred gene, *sbp*, was a member of both i-modulons but was not in the reported regulon. Genes with TF binding as determined by ChIP-exo are shown in bold. (e) Signs of i-modulon gene coefficients for eight i-modulons linked to dual regulators, colored by reported effect of regulator. (f) Ten media for predicted i-modulon activations. Correctly activated i-modulons are shown in bold. Distribution of i-modulon activities from pre-existing data includes all data from PRECISE excluding the ten validation conditions. The gray shaded region represents the average standard deviation across pre-existing i-modulon activities. All amino acid supplements were L-form, and all sugars were D-form. Abbreviations: GlcNAc, N-acetyl-glucosamine.

We identified MetJ binding sites upstream of 17 of the 18 genes in the MetJ i-modulon using ChIP-exo, increasing the i-modulon precision from 61% to 94% (Fig. 2c). The CysB regulon was split into the CysB i-modulon and the jointly regulated Cbl+CysB i-modulon (Fig. S3c). We identified CysB binding sites upstream of all genes in both i-modulons (Fig. 2d). TF binding was not detected near 10 of the 11 genes that were in the reported MetJ or CysB regulons but not in their respective i-modulons, potentially indicating inconsistencies in previous regulon definitions.

Although MetJ is a repressor, all MetJ i-modulon gene coefficients were positive. Increased repression by MetJ was therefore represented by negative i-modulon activities. Some TFs act as dual regulators, with the ability to activate certain genes and repress others. We evaluated the sign consistency of eight i-modulons linked to dual regulators (Fig. 2e). The regulation mode was previously reported for 56% of genes in the eight i-modulons. The majority of genes in five of the eight i-modulons were reported to be repressed, indicating that the linked TFs were primarily repressors. The remaining three TFs were primarily activators. Using these designations, the i-modulon gene coefficients were sign-consistent for 98% of genes with known regulation modes.

Application of ICA also extracted the condition-specific activities of i-modulons, providing an additional source of validation. I-modulon activities were normalized such that all i-modulon activities were zero for a baseline condition (See Methods). Thus, i-modulon activities under a particular condition represented the relative up- or down-regulation of the i-modulon genes compared to the baseline condition.

In order to validate that media perturbations predictably altered specific i-modulon activities, we designed 10 expression profiling experiments to conditionally activate 20 regulators. We confirmed 75% (15/20) of predicted activations through 13 i-modulons (Fig. 2f). The sign of the i-modulon activity revealed whether the effector resulted in a net activation or repression of i-modulon genes, which was consistent with known mechanisms for the TFs. The additional data introduced six new i-modulons, while maintaining the previously computed i-modulon structure (Fig. S3d).

Seven of the ten experiments included dual perturbations to simultaneously activate two regulators. In two cases, the two regulatory effects were recognized as a single signal, resulting in the combined i-modulons NagC/TyrR and GntR/TyrR (Fig. S3e,f). Cytidine supplementation did not activate the cytidine-binding transcription factor CytR; however, the previously-identified PurR-2 i-modulon was activated. Although four media additions did not activate their related i-modulons over the reference condition, the i-modulon structure of the TRN proved robust to additional data and displayed predictive capabilities.

### ICA Reveals Independent Modulation of Genes Within the PurR Regulon

In order to gain a detailed understanding of the biological roles of individual i-modulons, we programmatically generated a summary of characteristics for each i-modulon (See Supplemental Dataset 2). Figure 3 demonstrates these characteristics for two exemplary i-modulons enriched with genes in the PurR regulon, named PurR-1 and PurR-2, respectively.

**Fig. 3:**
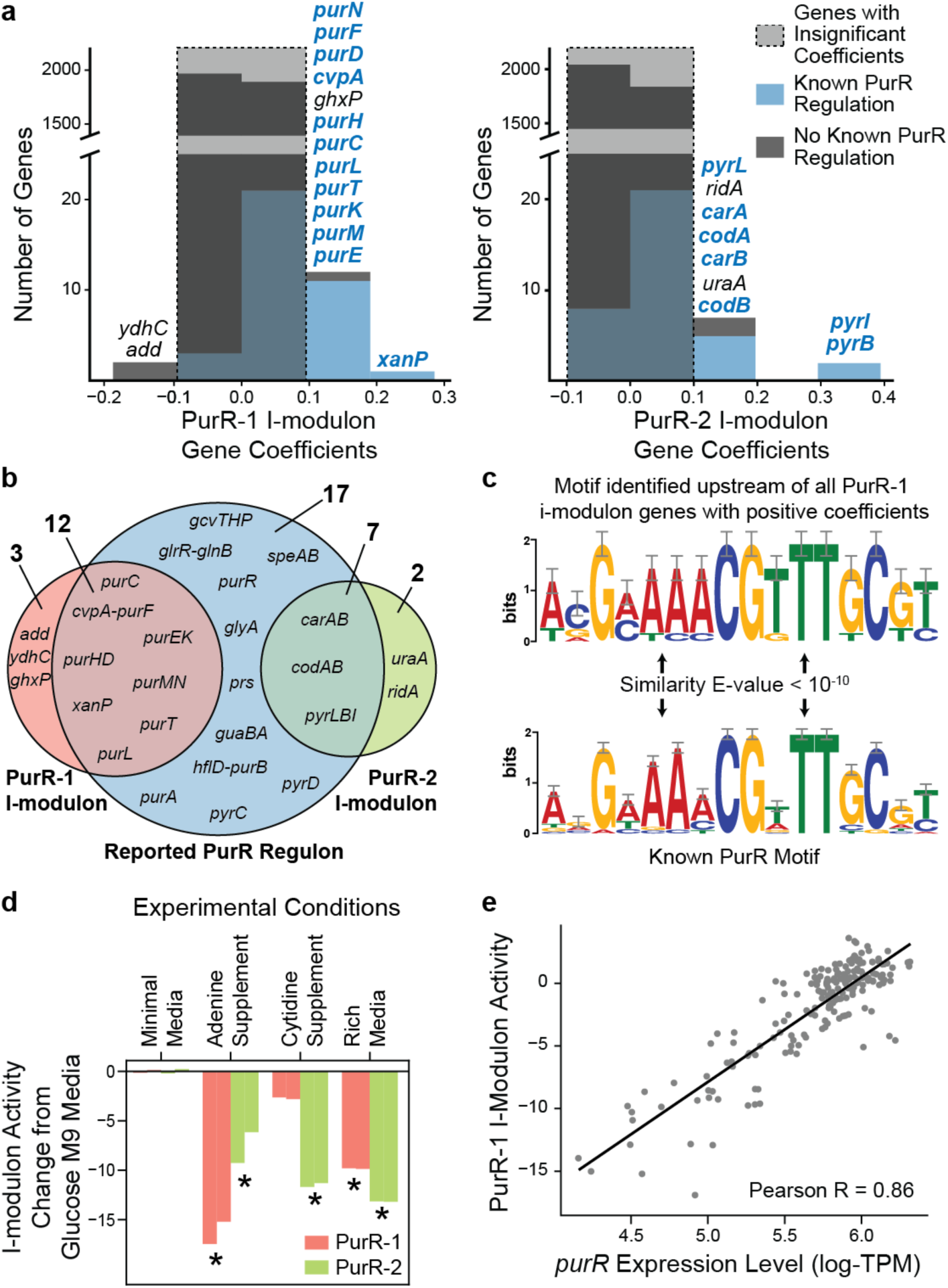
ICA Reveals Independent Modulation of Genes Within the PurR Regulon. (a) Histograms of gene coefficients in the PurR-1 and PurR-2 i-modulons. (b) Comparison of genes in the reported PurR regulon (blue), PurR-1 i-modulon (red) and PurR-2 i-modulon (green). (c) Motif identified upstream of genes in PurR-1 i-modulon compared to the reported PurR motif from RegulonDB 4. This motif was identified upstream of the guanine/hypoxanthine transporter encoding gene *ghxP*, although regulator binding was not previously reported. (d) The two PurR associated i-modulons exhibited distinct responses to environmental perturbations. (e) The PurR-1 i-modulon activity level is highly correlated with the log-transformed *purR* expression level across all conditions, whereas the PurR-2 i-modulon activity exhibits no correlation (See Fig. S4d). Of the 21 i-modulons whose activities were correlated with the expression of their associated regulons, 15 required a minimum expression level to activate the i-modulon (See Fig. S4c). Similar information on important i-modulons is available in Supplemental Dataset 2.

PurR is a repressor of nucleotide biosynthetic genes and is activated by the purines guanine and hypoxanthine^34^. The PurR-1 i-modulon contained 15 significant genes with both positive and negative coefficients, of which 13 were related to purine metabolism (See Fig. S2a). The PurR-2 i-modulon contained nine genes, of which eight were in the pyrimidine biosynthetic pathway (Fig. 3a). Together, the two PurR-related i-modulons accounted for 19 of the 36 genes in the reported PurR regulon (Fig. 3b). Segmentation of regulons into multiple constituent i-modulons was found for other global regulators such as Fur (Fig. S4a) and Crp (Fig. S4b).

The 13 genes with positive coefficients in the PurR-1 i-modulon were associated with purine biosynthesis, with 12 genes confirmed to be regulated by PurR. We detected the PurR motif upstream of the missing gene, suggesting that it is regulated by PurR (Fig. 3c). Similar analysis identified 68 previously unidentified regulator binding sites across 18 regulatory i-modulons (Table S3). The two genes with negative coefficients (*add* and *ydhC*) responded inversely to the activation of the purine biosynthetic pathway: *add*, which encodes the first enzyme in the purine degradation pathway; and *ydhC*, a putative transporter. Since *ydhC* expression was anticorrelated with purine biosynthetic gene expression, we hypothesized that *ydhC* was a purine-related efflux pump. In a similar fashion, we used the i-modulon structure to generate additional information for 87 genes with poor annotations, including 11 transporters (Table S4). One such prediction, *yjiY*, was recently independently verified as a pyruvate transporter and renamed to *btsT*^*35*^.

The condition-specific activities of the PurR-related i-modulons revealed differences in purine and pyrimidine biosynthetic gene expression (Fig. 3d). Adenine supplementation activated the repressor PurR to decrease the activity of the PurR-1 and PurR-2 i-modulons, whereas cytidine supplementation resulted in a decrease in PurR-2 activity. LB rich medium decreased both i-modulon activities.

The relationship between the quantitative activities of the PurR i-modulons and the expression level of PurR indicated the drivers of regulator activity. The activity of the PurR-1 i-modulon was highly correlated (Pearson R = 0.86, p-value < 10^-10^) with the expression level of *purR* (Fig. 3e). A similar relationship was observed in 21 of 50 regulatory i-modulons (Table 1). In contrast, the activity of the PurR-2 i-modulon was uncorrelated with *purR* expression (Pearson R = 0.24, p-value = 8*10^-4^), and was likely controlled by UTP-dependent reiterative transcription^36^ (Fig. S4d).

The results presented in this section demonstrate that the i-modulon structure of the TRN revealed by ICA provides a deep understanding of its biological functions and offers a guide to discovery.

### I-modulons Capture Coordinated Biochemical Signals

Although most i-modulons were linked to single regulators, some i-modulons appeared to be influenced by more than one regulator. For example, the Tryptophan i-modulon was enriched for TrpR-regulated genes but also included two genes (*tnaA* and *tnaB*) that are regulated by tryptophan-mediated transcriptional attenuation^31^ and respond inversely to the other genes in the i-modulon (Fig. S4e). Similarly the Leucine/Isoleucine (Leu/Ile) i-modulon contained 13 genes in 4 operons that are regulated by unique combinations of transcriptional attenuation^32^ and TF binding (Fig. S4f).

This phenomenon enabled us to interpret the Pyruvate i-modulon, named for its high activity in strains grown with pyruvate as the primary carbon source (Fig. 4a). This i-modulon was dominated by the pyruvate transporter gene *btsT*^*35*^ and the putative pyruvate transporter gene *yhjX* (Fig. 4b). The gene coefficients were consistent with their reported regulatory strategies; the BtsSR two-component system regulates *btsT* and contains a high-affinity pyruvate receptor, whereas the YpdAB two-component system regulates *yhjX* at lower pyruvate concentrations^37^. Four additional genes were regulated by the pyruvate-responsive regulator PdhR. When the genes in the Pyruvate i-modulon were mapped to the reactions they catalyzed, we observed that the Pyruvate i-modulon responded to increased extracellular and intracellular pyruvate levels to increase fermentation and activate acetate overflow metabolism (Fig. 4c)^39^.

**Fig. 4:**
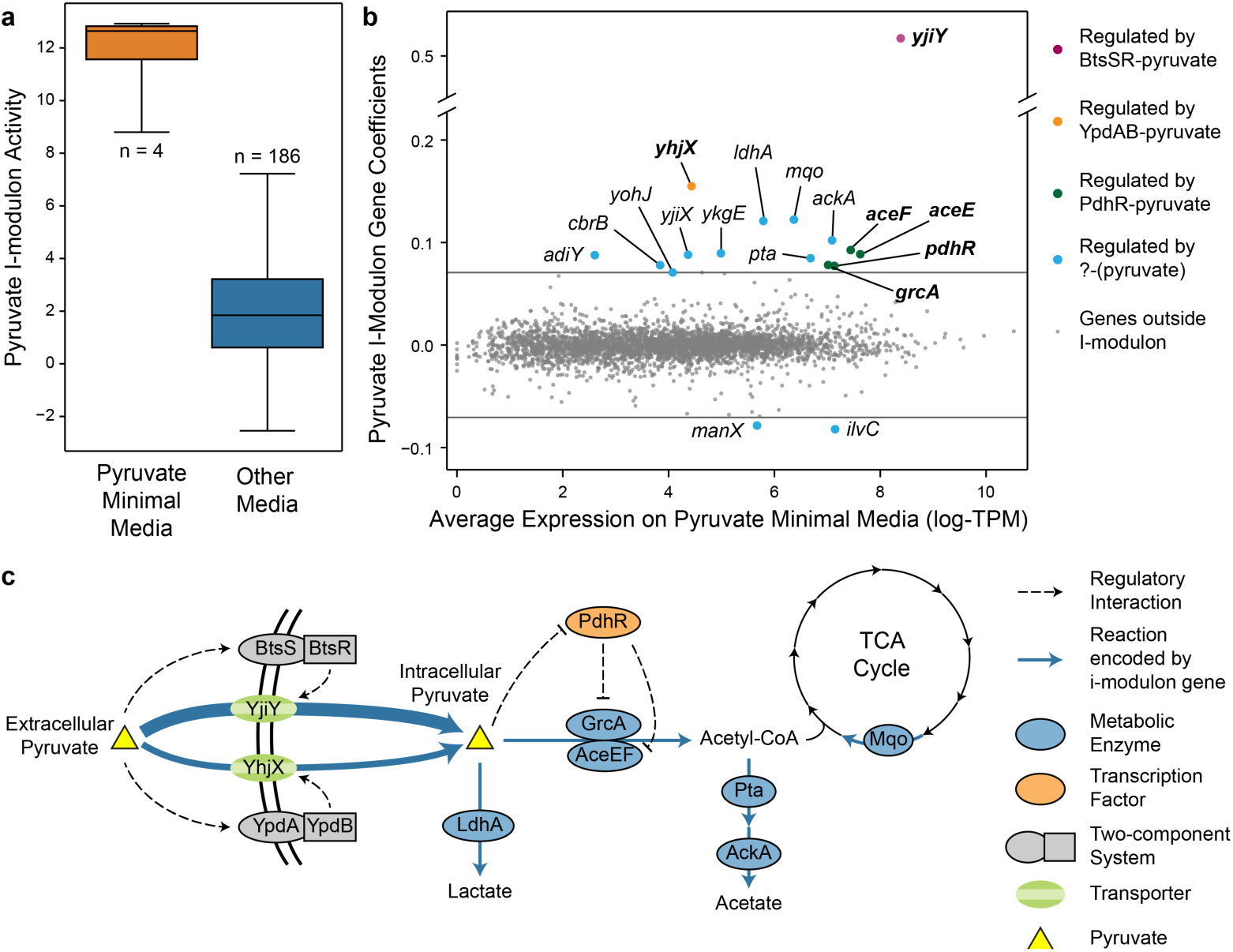
The Pyruvate I-modulon. (a) Boxplot of Pyruvate i-modulon activities. Number of expression profiles in each box are shown. (b) Scatter plot of Pyruvate i-modulon gene coefficients against average gene expression on pyruvate minimal media. I-modulon genes are colored by reported or predicted regulation. See Fig. S4e,f for other i-modulons associated to multiple related regulators. (c) Schematic illustration of the metabolic roles and transcriptional regulation of genes in the Pyruvate I-modulon. Extracellular pyruvate activates the two-component systems BtsSR and YpdAB to upregulate the expression of pyruvate transporter-encoding genes *yjiY* and *yhjX*. High levels of intracellular pyruvate antagonize the PdhR repressor, increasing conversion of pyruvate to acetyl-CoA. Although the regulatory strategy is unknown for the remaining genes, we hypothesize that they are controlled by pyruvate-sensing TFs.

### Two i-modulons characterize the ‘Fear vs. Greed’ Tradeoff

Can the i-modulon decomposition be utilized to understand a major genetic perturbation of the transcriptome? Adaptive laboratory evolution of *E. coli*^*40*^ revealed two distinct point mutations in the RNA polymerase subunit *β* that shift cellular resources towards growth-related functions (i.e. greed) away from stress-hedging functions (i.e. fear)^41^.

The quantitative i-modulon activities for two RpoB mutant strains reflected this trade-off (Fig. 5a). Three i-modulons whose activities significantly deviated from the wild-type strain were initially uncharacterized. Relaxation of the gene coefficient threshold revealed genes encoding translation machinery, such as ribosomal proteins, to comprise one of the uncharacterized i-modulons (Fig. 5b). The compendium-wide activity of this Translation i-modulon was correlated with growth rate (Pearson R = 0.67, p-value < 10^-10^, Fig. S4g), consistent with previous observations that growth is propelled by increased ribosomal catalytic activity^42–45^. The Translation i-modulon therefore represented the “greedy”, growth-related functions of the transcriptome.

**Fig. 5:**
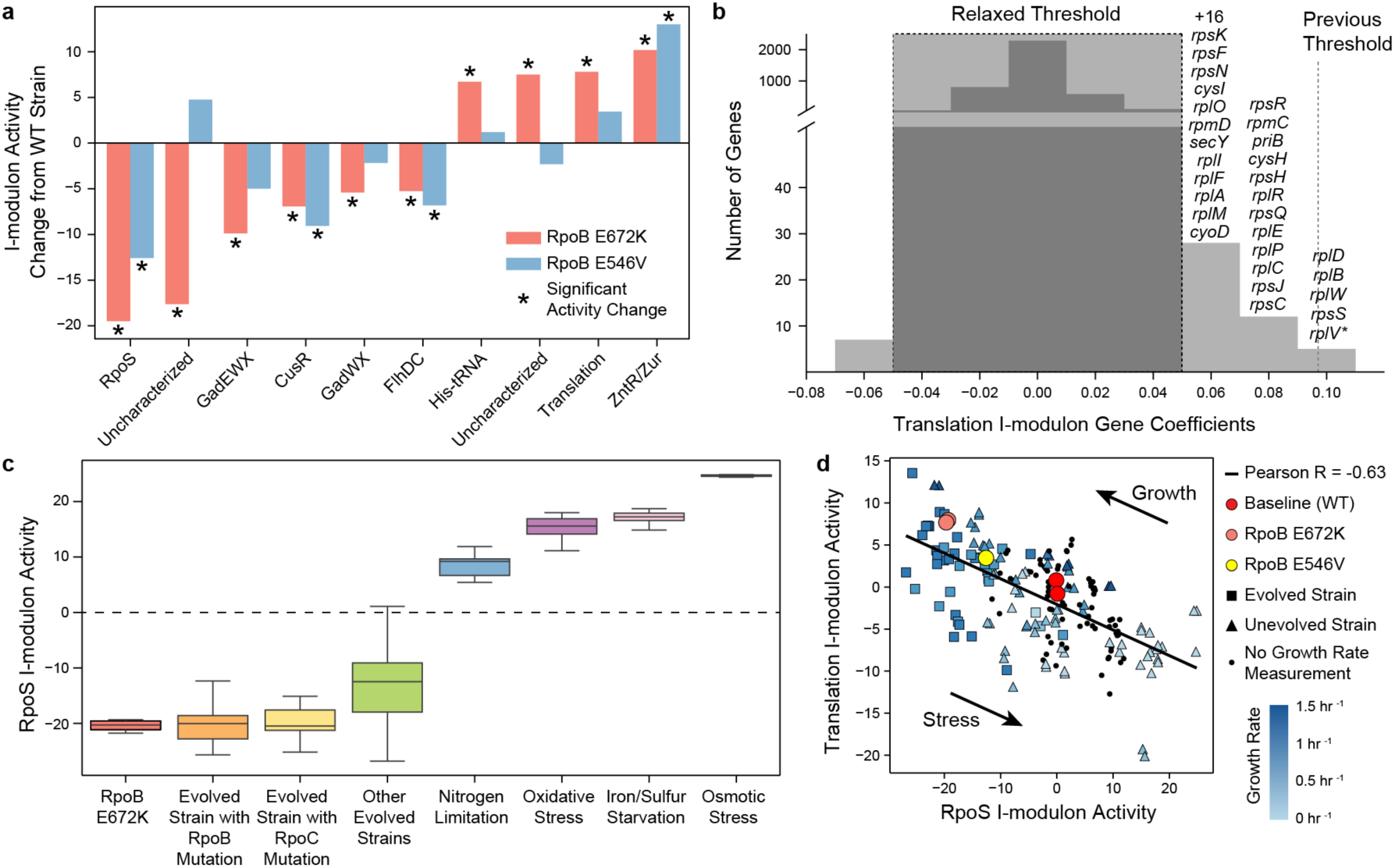
Two i-modulons characterize the ‘Fear vs. Greed’ Tradeoff. (a) Comparison of i-modulon activities in the RpoB E672K and RpoB E546V mutant strains grown on glucose minimal media against wild-type activities. Significant i-modulon activities are designated by asterisks. For detailed information about these i-modulons, see Supplemental Dataset 2. (b) Histogram of translation i-modulon gene coefficients. Gene names are shown for genes above relaxed threshold. The single gene outside the original threshold was *rplV*, marked with an asterisk. (c) The RpoS i-modulon activities revealed the stress level of the cell under various conditions. (d) The RpoS i-modulon activities were anti-correlated with the Translation i-modulon activities, highlighting the trade-off between stress-hedging and growth. Single nucleotide mutations in RpoB (in yellow and red) shift cellular resources along this line.

The i-modulon with the largest activity decrease in both variants was enriched in genes controlled by the stress response sigma factor (RpoS). The RpoS i-modulon activity was correlated (Pearson R = 0.67, p-value < 10^-10^, Fig. S4h) with the expression level of *rpoS* and revealed a quantitative measure of cellular stress across diverse conditions (Fig. 5c). Therefore, the RpoS i-modulon represented the “fearful” stress-hedging functions of the transcriptome.

PRECISE also contains expression data for over 30 adaptive laboratory evolution endpoint strains^46^, many of which contain mutations in genes encoding RNA polymerase subunits. All strains with mutated *rpoB* or *rpoC* genes exhibited low RpoS i-modulon activity, reflecting a reduction in stress-related expression. Further examination revealed that the RpoS i-modulon activity was anti-correlated with the Ribosomal i-modulon activity (Pearson R = −0.63, p < 10^-10^), illuminating the compendium-wide transcriptomic trade-off between fear and greed (Fig. 5d). The mutations in *rpoB* shift the strains along this line, increasing growth and reducing stress-related gene expression.

The results presented in this section show that the fear-greed tradeoff, and other transcriptome restructuring events, can be studied in great detail by decomposing the transcriptome into a summation of independent regulatory events.

### An I-modulon Identifies Transcriptional Differences Between Two Closely-Related *E. coli* Strains

The PRECISE database includes 30 RNA-seq datasets from *Escherichia coli* BW25113, a closely related strain to *E. coli* MG1655. The BW25113 strain was the background strain for the Keio collection of over 3,000 single-gene knock-outs^47^. The transcriptomic differences between these strains, resulting from 29 genetic variations^48^, have not been characterized. We identified a single i-modulon whose activities separated the transcriptomes of the two strains (Fig. 6a). The i-modulon gene coefficients elucidated twelve transcriptomic differences explained by genetic differences between the strains (Fig. 6b and Table S5).

**Fig. 6:**
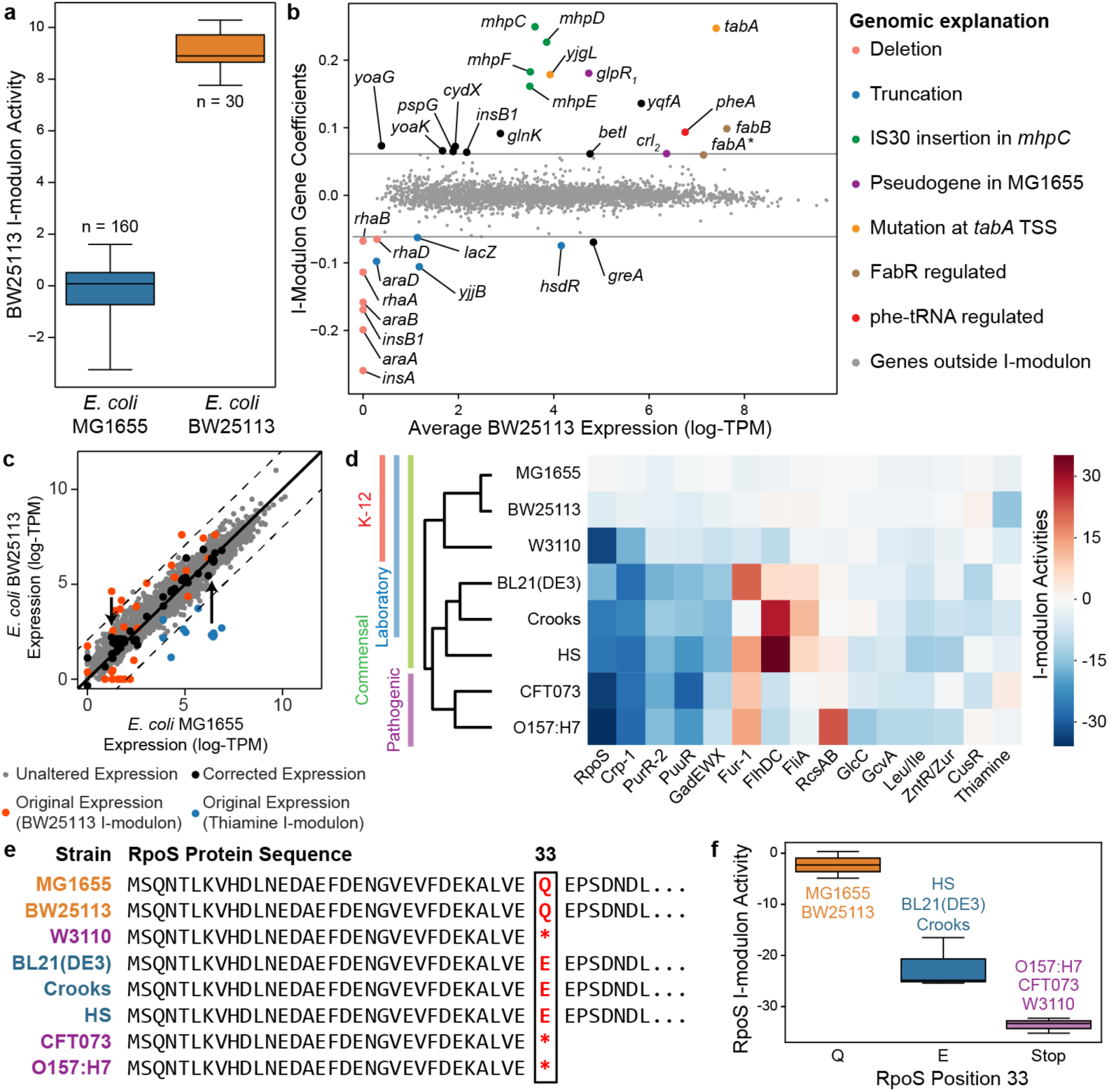
I-modulons Identify Differences in Transcriptional Regulation Across Multiple *E. coli* Strains. (a) BW25113 i-modulon activities separated by strain. Number of expression profiles from each strain is shown. (b) Scatterplot of average BW25113 expression against BW25113 i-modulon activity. Deletions and truncations in the BW25113 strain account for 11 of the 12 genes with negative coefficients. An insertion sequence (IS30) in the mhpC gene in the BW25113 strain corresponds to a large increase in expression of *mhpCDEF*, as IS30 contains a known promoter54. Two genes in the BW25113 strain, *crl* and *glpR*, are non-functional pseudogenes in MG1655 due to an internal frameshift. Their positive i-modulon gene coefficients imply an increase in expression when the genes are functional. Point mutations at the predicted transcription start site (TSS) of *tabA*, in the FabR regulator, and in the phenylalanine tRNA *pheV*, account for other genes with positive coefficients (See Table S5). Asterisks denote genes marginally within gene coefficient threshold. (c) Heatmap of estimated i-modulon activities for 8 *E. coli* strains grown on glucose minimal media (with added thiamine and ferric chloride for BW25113). Only significantly altered regulatory i-modulon activities are shown. Hierarchical clustering of the activities reflects strain-specific characteristics. (d) Subtraction of the BW25113 and Thiamine i-modulons from the *E. coli* BW25113 expression profile accounts for the major transcriptomic deviations from *E. coli* MG1655 grown without thiamine. Dashed lines indicate two-fold difference in TPM. (e) Sequence alignment of the RpoS protein across the eight *E. coli* strains. (f) RpoS activities of the eight strains grouped by position 33 in the RpoS protein sequence, as detailed in panel (e).

We then sought to use the summation of i-modulons to quantitatively account for these strain-specific differences. To this end, we analyzed two expression profiles in the compendium: the baseline condition (wild-type *E. coli* MG1655) and *E. coli* BW25113 with thiamine and ferric chloride supplementation. Only two i-modulons were differentially activated between the two conditions: the BW25113 i-modulon (described above) and the Thiamine i-modulon that controls thiamine biosynthesis through a riboswitch^30^. We accounted for the transcriptomic differences by subtracting the gene coefficients in the two i-modulons scaled by their respective i-modulon activities from the *E. coli* BW25113 expression profile. This increased the R^2^ between the two strains’ expression profiles from 0.93 to 0.94 (Fig. 6c), illustrating that the summation of the independently modulated genes captured the major expression differences between the two strains.

In the previous section we used the i-modulon structure to interpret the effects of a single mutation in *rpoB* on the transcriptome. Here, we illustrated the broader ability of the i-modulons to interpret and quantitatively account for the transcriptional differences between closely related strains.

### I-modulons Identify Differences in Transcriptional Regulation Across Multiple *E. coli* Strains

It has proven difficult to compare transcriptional regulation between different strains of the same species^49^. We examined the ability of i-modulons to provide a structured basis for such comparison. We grew and expression profiled a set of eight diverse *E. coli* strains in identical media (with additional supplements for BW251123 as described above) and calculated their i-modulon activities using the 70 previously identified i-modulons as a basis (Fig. 6d). Three strains (MG1655, BW25113 and W3110) diverged from the same ancestral K-12 strain with limited genetic differences; CFT0173 and O157:H7 EDL933 are pathogenic strains; and the remaining three strains (BL21(DE3), HS, and Crooks) are laboratory strains.

The expression profiles of the K-12 strains shared similar i-modulon activities, including higher activities in the pyrimidine-responsive PurR-2 i-modulon. This can be explained by a defect in the *rph-pyrE* operon, which leads to pyrimidine starvation in the K-12 strains^50^. The RpoS i-modulon activity was significantly suppressed in all strains except MG1655 and BW25113. Strains W3110, CFT073, and O157:H7 EDL933 have an amber mutation that results in a truncation in *rpoS* and is known to reduce its expression and activity^51,52^. Three other strains (BL21(DE3), HS, and Crooks) contain a divergent mutation at the same position that likely explains their reduced RpoS i-modulon activity (Fig. 6e,f).

These results demonstrate that the i-modulons derived from a single strain can provide a scaffold to analyze transcriptomes from other strains of the same species. Strain-specific differences in i-modulon activity can be traced to sequence variations in the associated regulator, thus providing a deep explanation for targeted strain differences.

## Discussion

We have demonstrated that the combination of (1) independent component analysis of high-quality RNAseq data and (2) high-resolution comprehensive regulator binding site information, identifies linear combinations of quantitative regulatory signals that reconstitute the *E. coli* transcriptome. This result suggests that a *principle of i-modulon addition* governs the composition of the *E. coli* transcriptome. Application of this principle provided a multidimensional understanding of the *E. coli* TRN, and uncovered detailed responses to environmental and genetic perturbations, optimality of adaptation to new conditions, and links between genotype and phenotype of *E. coli* strains. If this principle governs the composition of other prokaryotic transcriptomes, it provides a path to develop detailed understanding of transcriptional regulation in less-understood organisms.

We have shown that for the model prokaryote *E. coli*, 50 of the 70 identified i-modulons represent the effects of characterized transcriptional regulators. This coverage is a consequence of the quality and diversity of the RNA-seq compendium used, the extensive information available on *E. coli* transcriptional regulators, and the relative simplicity of the *E. coli* TRN. In principle, if we obtained expression data for every condition sensed by a prokaryote, we could decompose its expression state to a non-reducible set of regulatory signals. These fundamental signals, combined with high-throughput binding data for all TFs in an organism^38^, would lead to the establishment of a comprehensive quantitative transcriptional regulatory network.

## Methods

### RNA extraction and library preparation

Total RNA was sampled from duplicate cultures. All strains were grown in minimal salts (M9) medium at exponential phase, with complete growth conditions listed in Supplemental Dataset 1. Growth curve analysis were performed using Bioscreen C Reader system with 200 uL culture volume per well. Two biological replicates were used in the assay. Media components were purchased from Sigma-Aldrich (St. Louis, MO). For nitrate respiration cultures, a 35:50 ratio of carbon dioxide to nitrogen was bubbled through the media to deoxygenate. After inoculation and growth, 3 mL of cell broth (OD_600_ ∼ 0.5) was immediately added to 2 volumes Qiagen RNA-protect Bacteria Reagent (6 mL), vortexed for 5 seconds, incubated at room temperature for 5 min, and immediately centrifuged for 10 min at 17,500 RPMs. The supernatant was decanted, and the cell pellet was stored in the −80°C. Cell pellets were thawed and incubated with Readylyse Lysozyme, SuperaseIn, Protease K, and 20% SDS for 20 minutes at 37°C. Total RNA was isolated and purified using the Qiagen RNeasy Mini Kit columns and following vendor procedures. An on-column DNase-treatment was performed for 30 minutes at room temperature. RNA was quantified using a Nano drop and quality assessed by running an RNA-nano chip on a bioanalyzer. The rRNA was removed using Illumina Ribo-Zero rRNA removal kit for Gram Negative Bacteria. A KAPA Stranded RNA-Seq Kit (Kapa Biosystems KK8401) was used following the manufacturer’s protocol to create sequencing libraries with an average insert length of around ∼300 bp. Libraries were ran on a HiSeq4000 (Illumina).

### ChIP-exo preparation

ChIP-exo experimentation was performed following the procedures previously described ^55^. To activate each TF, cells were grown on relevant media: M9 minimal medium with 2 g/L glucose and 5 mM methionine for MetJ, and M9 minimal medium with 2 g/L glucose and 0.25 mM taurine for CysB. In brief, to identify binding maps for each TF, DNA bound to each TF from formaldehyde cross-linked E. coli cells were isolated by chromatin immunoprecipitation (ChIP) with the specific antibodies that specifically recognize myc tag (9E10, Santa Cruz Biotechnology), and Dynabeads Pan Mouse IgG magnetic beads (Invitrogen) followed by stringent washings as described previously ^56^. ChIP materials (chromatin-beads) were used to perform on-bead enzymatic reactions of the ChIP-exo method ^33^. Briefly, the sheared DNA of chromatin-beads was repaired by the NEBNext End Repair Module (New England Biolabs) followed by the addition of a single dA overhang and ligation of the first adaptor (5′-phosphorylated) using dA-Tailing Module (New England Biolabs) and NEBNext Quick Ligation Module (New England Biolabs), respectively. Nick repair was performed by using PreCR Repair Mix (New England Biolabs). Lambda exonuclease- and RecJf exonuclease-treated chromatin was eluted from the beads and overnight incubation at 65°C reversed the protein-DNA cross-link. RNAs- and Proteins-removed DNA samples were used to perform primer extension and second adaptor ligation with following modifications. The DNA samples incubated for primer extension as described previously were treated with dA-Tailing Module (New England Biolabs) and NEBNext Quick Ligation Module (New England Biolabs) for second adaptor ligation. The DNA sample purified by GeneRead Size Selection Kit (Qiagen) was enriched by polymerase chain reaction (PCR) using Phusion High-Fidelity DNA Polymerase (New England Biolabs). The amplified DNA samples were purified again by GeneRead Size Selection Kit (Qiagen) and quantified using Qubit dsDNA HS Assay Kit (Life Technologies). Quality of the DNA sample was checked by running Agilent High Sensitivity DNA Kit using Agilent 2100 Bioanalyzer (Agilent) before sequenced using HiSeq 2500 (Illumina) following the manufacturer’s instructions. ChIP-exo experiments were performed in biological duplicate.

### ChIP-exo processing

ChIP-exo was processed as described previously^55^. Briefly, sequence reads obtained from ChIP-exo experiments were mapped onto the E. coli reference genome (NC_000913.3) using bowtie^57^ with default options in order to generate SAM output files. MACE program^58^ was used to define peak candidates from biological duplicates for each experimental condition with sequence depth normalization. Then, each peak was assigned to the nearest operon on either side, using operon definitions from RegulonDB. Only operons 500 basepairs downstream of peak were considered. Final operons on forward strand were required to be in front of the peak, and operons on reverse strand were required to be behind the peak. Genome-scale data were visualized using MetaScope to manually curate peaks (http://systemsbiology.ucsd.edu/Downloads/MetaScope).

### Compilation of PRECISE

Raw sequencing reads were collected from GEO (See Supplemental Data 1 for accession numbers) or produced in the lab, and mapped to the reference genome (NC_000913.3 for strain MG1655 and CP009273 for BW25113) using bowtie 1.1.2^57^ with the following options “-X 1000 -n 2 −3 3”. Transcript abundance was quantified using *summarizeOverlaps* from the R *GenomicAlignments* package, with the following options “mode=“IntersectionStrict”, singleEnd=FALSE, ignore.strand=FALSE, preprocess.reads=invertStrand”^59^. Genes shorter than 100 nucleotides and genes with under 10 fragments per million mapped reads across all samples were removed before further analysis. Transcripts per Million (TPM) were calculated by *DESeq2*. The final expression compendium was log-transformed *log*_*2*_*(TPM+1)* before analysis, referred to as log-TPM. Biological replicates with R^2^ < 0.9 between log-TPM were removed to reduce technical noise.

### Compilation of the reported *E. coli* regulatory network

We compiled the global TRN using all interactions from RegulonDB 10.0^3^ for both transcription factor and sRNA binding sites. Binding sites were added from recent studies, as described in Fang et al.^27^, in addition to binding sites for Nac and NtrC^60^ and binding sites for 10 uncharacterized transcription factors ^38^. We also included sigma factor binding sites, riboswitch information, and transcriptional attenuation from Ecocyc^61^. When reported, mode of effect (i.e. activation or repression) was included. If the effect was unreported, or multiple effects were reported, effects were designated as unknown. All genes absent from PRECISE were removed from the final TRN.

### Computing robust independent components

We first normalized the compendium using wild-type *E. coli* MG1655 grown on glucose M9 minimal media as the baseline condition (labelled *base__wt_glc__1* and *base__wt_glc__2*). We subtracted the mean expression of each gene in these two samples from the compendium to calculate log_2_-fold-change (LFC) deviations from the baseline.

We used the Scikit-learn^62^ implementation of the FastICA algorithm^53^ to identify independent components. We executed FastICA 256 times with random seeds, a convergence tolerance of 10^-8^, log(cosh(x)) as the contrast function, and the parallel search algorithm. We constrained the number of components in each iteration to the number of components that reconstruct 99% of the variance as calculated by principal component analysis (138 components).

The resulting source components (**S**) from each run were clustered using the Scikit-learn implementation of the DBSCAN algorithm^63^ with epsilon of 0.1, and minimum of 50% of samples to create a cluster seed. DBSCAN does not require predetermination of the number of clusters, and does not require that all points belong to a cluster. The dimensionality of the dataset is therefore estimated by the number of clusters calculated by DBSCAN. The components computed by FastICA are standardized by default, with a mean of 0 and an L2-norm of 1. However, identical components from separate runs may have opposite signs. Therefore we used the following distance metric:

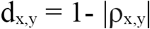

where ρ_x,y_ is the Pearson correlation between components x and y. Each component in a cluster was then inverted if necessary to ensure that the gene with the maximum absolute value in the component had a positive weight, creating sign-consistent clusters. The final independent components were defined as the centroid of each cluster in **S**, and the weightings were defined as the centroid of their corresponding weighting vectors in **A**.

In order to ensure that the final components were consistent across multiple runs, we computed the clustered components 100 times, and found that 70 components were identified in every run, which were the final robust components used in the analysis.

In order to confirm that the i-modulon structure was generally invariant to the composition of the expression database, we applied ICA to two subsets of PRECISE. The first subset consisted of all previously generated data from unevolved *E. coli* (48 profiles), and the second subset consisted of all previously generated data (170 profiles). We compared the resulting components using the absolute value of the pearson correlation coefficient. The resulting network was graphed using the Graphviz^64^ python library (Fig. S3c). Correlations below 0.5 were discarded as insignificant.

### Determination of the gene coefficient threshold

Each component in **S** contains the contributions of each gene to the statistically independent source of variation. Most of these values are near zero for a given component. In order to identify the most significant genes in each component, we modified the method proposed in Frigyesi et al.^65^. For each component, we iteratively removed genes with the largest absolute value and computed the kurtosis of the resulting distribution. Once the kurtosis fell below a cutoff, we designated the removed genes as significant.

To identify this cutoff, we performed a sensitivity analysis on the concordance between significant genes in each component and known regulons. Eleven components were initially identified to have high concordance (precision > 0.75 and recall > 0.25) with a single regulator, using an initial kurtosis cutoff of 4. We varied the kurtosis cutoff from 1 through 10 in increments of 0.5, and computed the F1-score (harmonic average between precision and recall) between each component and its linked regulator. The maximum value of the average F1-score across the 11 components occurred at a kurtosis cutoff of 4.5 (See Fig. S5a-c).

Since each set of significant genes represents a set of independently modulated genes, we henceforth refer to these gene sets as “i-modulons”. Since independent components have no canonical direction, we inverted i-modulons (and related activities) such that the number of positive genes in an i-modulon was always larger than the number of negative genes.

### Associating regulators to i-modulons

We compared the set of significant genes in each i-modulon to each regulon (defined as the set of genes regulated by any given regulator) using Fisher’s Exact Test (FDR < .01). Additionally, combined regulon enrichments were calculated to identify joint regulation of i-modulons (such as NtrC+RpoN and NagC/TyrR), using both intersection (+) and union (/) of up to four regulons. Final i-modulon-regulator associations were determined through manual curation of enriched regulators.

### Cumulative explained variance for ICA

Components were initially ordered by the L2-norm (sum of squares) of for each row in the **A** matrix for ICA. Cumulative explained variance was calculated for component *K*:

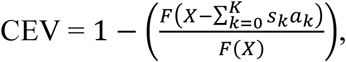

Where F(**A**) is the square of the Frobenius norm

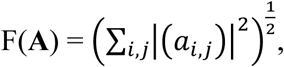

**X** is the original expression profile, *s*_*k*_ is the kth column in the **S** matrix, and *a*_*k*_ is the kth row in the **A** matrix.

### Comparison of microarray data and PRECISE using ICA and sparse-PCA

Microarray data was acquired from NCBI GEO Series GSE6836^22^. This dataset had similar size to PRECISE (212 experiments) to ensure comparability. Microarray data was processed using the RMA R package^66^. ICA was performed as described above for both datasets. PCA determined that 148 components reconstructed 99% of the variance in the microarray dataset. Sparse-PCA was performed using the *elastic net* R package^67^, searching for the same number of components as with ICA, and a vector of ones as thresholding parameters. The resulting components from sparse-PCA and ICA were compared against single regulators, using the kurtosis threshold with cutoff of 4.5 as discussed above. Regulators with highest F1-score were assigned to each component to ensure consistency in the comparison (no manual curation was used to generate the comparison figure).

### Differential activity analysis

We first computed the distribution of differences in i-modulon activities between biological replicates, and then fit a log-normal distribution to each distribution. We confirmed that the difference in activities between biological replicates followed a log-normal distribution for all i-modulons using the Kolmogorov-Smirnov test and validating through quantile-quantile plots (Fig. S5d-f).

To test for differential activity of an i-modulon between two different conditions, we first computed the average activity of the i-modulon between biological replicates, if available. We then computed the absolute value of the difference in i-modulon activities between the two conditions. This difference was compared against the log-normal distribution for the i-modulon to calculate a p-value. I-modulons were designated as significant if the absolute value of their activities was greater than 5, and p-value was below 0.01.

### I-modulon summation

We selected samples *base__wt_glc__1* and *base__wt_glc__2* to represent the wild-type cell, and samples *omics__bw_glc__1* and *omics__bw_glc__2* to represent the mutated strains to be corrected. The average activities between the replicates were used for the corrections. The corrections were applied to the BW25113 and Thiamine i-modulons.

The i-modulon decomposition is based on the equation **X=SA**, where *x*_*j*_ = Σ*s*_*i*_\**a*_*ij*_ for a particular expression profile *j*, where *i* represents an independent component. We aim to produce the correction (*x*_*2*_′) to the expression profile (*x*_*2*_) with respect to a baseline expression profile (*x*_*1*_) for all differentially activated i-modulons *i* ∈ I:

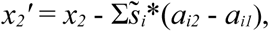

where 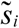 is a vector of zeros except for significant gene coefficients in i-modulon *i*, and *a*_*ij*_ is the activity of i-modulon *i* under condition *j*.

### RNA-seq processing and i-modulon projection for multiple strain comparison

Raw sequencing reads and transcriptome abundance were identified similar to as described in the section above, using the following reference genomes: NC_000913.3 (MG1655), CP009273 (BW25113), NC_007779.1 (W3110), NC_010468.1 (Crooks), NC_012971.2 (BL21(DE3)), NC_009800.1 (HS), CP008957.1 (O157:H7 EDL933), and NC_004431.1 (CFT073). Since gene composition varies across E. coli strains, we filtered the transcriptomes to only include the 1033 genes that were members of at least one i-modulon. Genes absent from a particular strain with respect to the reference strains (MG1655) were assumed to have zero expression. We calculated the log_2_(TPM + 1) values using the same normalization to baseline conditions as described above.

Thereafter, we calculated the i-modulon activities for the eight new *E. coli* expression profiles using the previously identified 70 independent components (including all gene coefficients). We projected the eight new expression profiles (**X′**) onto the previously computed basis (**S**):

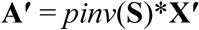

where **A′** represents the i-modulon activities for the eight strains, and *pinv* is the pseudo-inverse function. This represents the least-squares approximation of **A**.

### Automated characterization of i-modulons

Supplemental Dataset 2 contains detailed information on important i-modulons. Each page characterizes one i-modulon, which are sorted by name. The Cra i-modulon page is described in Fig. S6a-g.

The scatter plot (Fig. S6a) compares the i-modulon gene weights to the baseline expression (defined as the average log-TPM of sample IDs *wt_glc__wt_glc__1* and *wt_glc__wt_glc__2*). The horizontal dashed lines represent the threshold separating significant and non-significant genes in an i-modulon. Each gene is colored by its functional category, defined by clusters of orthologous groups of genes queried from EggNOG^68^. Gene names are provided on the scatterplot if total number of significant genes is below 25. Subscripts indicate split genes in *E. coli* MG1655, classified as pseudogenes.

The bar chart (Fig. S6b) displays the i-modulon activities across all conditions in the database, grouped by study. Each condition shares the same total width, regardless of number of biological replicates. The sample IDs correspond to the IDs in Supplemental Dataset 1.

The histograms (Fig. S6c) show the gene weights in an i-modulon on a log-scale, split by enriched regulon. The vertical bars represent the significance threshold for the component. The gray histogram captures genes that are not in an enriched regulon. The brown histogram captures genes that are in more than one enriched regulon.

The venn diagram (Fig. S6d) compares genes in an i-modulon (above the threshold) to genes in the enriched regulons. If the i-modulon name contains a slash (/), the regulon circle contains the union of genes in any enriched regulon (i.e. genes in either regulon). If the i-modulon name contains a plus sign (+), the regulon circle contains the intersection of the regulons (i.e. genes in both regulons). Numbers in parentheses indicate number of operons (complete or partial) in each category. Operons were defined using RegulonDB. This panel is hidden if no enriched regulons were identified.

The regulon scatter plot (Fig. S6e) compares the expression level of enriched regulators to the i-modulon activity. Each point represents an expression profile. Two lines were fit to each scatter plot, a simple line and a broken line. The broken line represents a minimal expression level required before a correlation is observed between the i-modulon activity and the regulator expression. Only the line with the highest *R*^*2*^_*adj*_ is shown, where

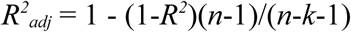

The broken line is modeled as:

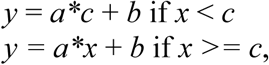

where *x* is the expression level of the gene encoding the TF, and *y* is the i-modulon activity level. Parameter *c* is optimized using the *optimize.curve_fit* function in the Python *scipy* package^69^. Three initial values for *c* were tested to identify the optimal fit: minimum expression, maximum expression, and mean expression.

This panel is hidden if no enriched regulons were identified, or if expression levels were not measured for the regulator (see Thiamine i-modulon). If there are two or more enriched regulons with expression levels, the two TFs with the strongest association to the i-modulon are shown.

### Motif Search

The motifs (Fig. S6f) were identified by searching upstream sequences using MEME^70^. Genes in each i-modulon were grouped into operons. A stringent upstream sequence was defined as the region from -300 to the start of the operon. We searched for zero or one motif per sequence using an E-value threshold of 10^-3^, searching for motifs with all widths between 6 and 30 basepairs for all non-genomic i-modulons. For i-modulons with no enriched motifs in the stringent upstream sequence, we searched for motifs in a broader upstream sequence of −600 to +100, keeping all other parameters constant. The E-value of the motif and the percent of i-modulon operons with the upstream motif are listed below the motif.

The motif comparison (Fig. S6g) was generated by comparing i-modulon motifs against known motifs from RegulonDB using TOMTOM^71^, with an E-value threshold of 10^-3^ and allowing incomplete matches. The E-value of the comparison is shown below the figure.

For regulatory i-modulons, genes with identified motif sites but no known regulation by the associated TF are reported in Table S4.

## Acknowledgements

We thank Dr. Joe Pogliano for biological insights, Dr. David Heckmann, Dr. James Yurkovich, Dr. Jared Broddrick, Erol Kavvas, Jean-Christophe Lachance, and Dr. Ke Chen for helpful discussions and Marc Abrams for editorial comments. This research used resources of the National Energy Research Scientific Computing Center, a DOE Office of Science User Facility supported by the Office of Science of the U.S. Department of Energy under Contract No. DE-AC02-05CH11231. This work was funded by the Novo Nordisk Foundation Center for Biosustainability and the Technical University of Denmark (grant number NNF10CC1016517).

## Author Contributions

A.V.S. designed the study and performed the analysis, with assistance from L.Y. and B.O.P. Y.G., R.S., Y.H., and S.X. performed expression profiling experiments. Y.G. performed ChIP-exo experiments. D.K. performed expression profiling for multi-strain expression analysis, and K.S.C. analyzed the multi-strain data. L.Y. and Z.A.K. provided technical support and conceptual advice. A.V.S. and B.O.P. wrote the manuscript.

## Competing Interests

The authors declare no competing interests

## Data availability

Unprocessed RNA-seq and ChIP-exo data reported in this paper are deposited in NCBI GEO with primary accession codes GSE122211, GSE122295, GSE122296, and GSE122320.

## Supplemental Information

Supplemental Methods

Figure S1: Summary of data

Figure S2: Overview of i-modulons

Figure S3: Validation of i-modulon regulator relationships Figure

Figure S4: Additional properties of i-modulons

Figure S5: Significance thresholds for components and differential activity

Figure S6: Descriptive Characteristics of the Cra I-modulon with Explanations

Table S1: ChIP-exo binding sites for MetJ

Table S2: ChIP-exo binding sites for CysB

Table S3: Computationally-detected novel TF binding sites upstream of i-modulon genes

Table S4: I-modulon membership for uncharacterized genes as defined by Ghatak et al.^72^

Table S5: Genes in BW25113 i-modulon

Supplemental Dataset 1: Expression levels, experimental conditions, and i-modulon decomposition of the PRECISE dataset

Supplemental Dataset 2: Descriptive characteristics of 30 selected i-modulons

