## Supplemental Information for "The *Escherichia coli* Transcriptome Mostly Consists of Independently Regulated Modules"

### Supplemental Figures:

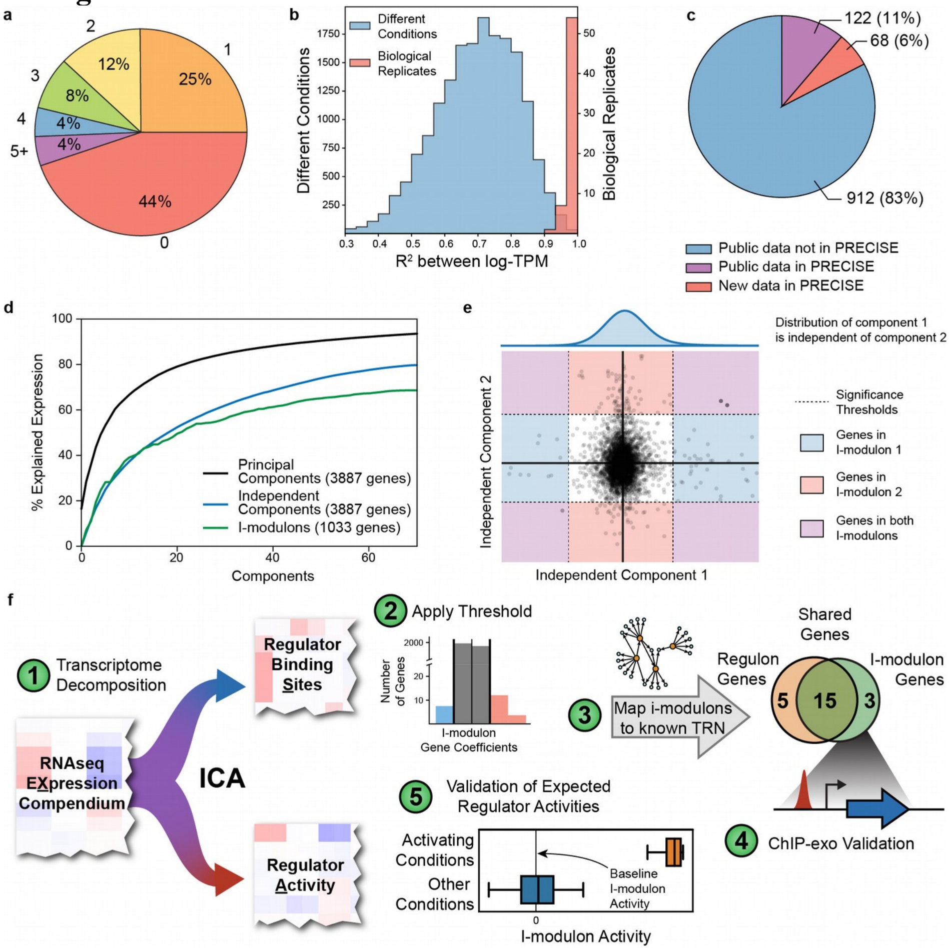


**Fig. S1: Summary of data** (a) Distribution of TF connectivity in *Escherichia coli*. The pie chart indicates the fraction of genes that are regulated by the indicated number of TFs. (b) Histogram of data quality, as measured by the coefficient of determination ($R^2$) between log-TPM. Comparison between biological replicates are shown in red, whereas comparison between all pairwise non-replicates are shown in blue. (c) Pie chart of all published RNA-seq data for *E. coli* MG1655 or BW25113 on NCBI GEO as of January 2018. (d) Cumulative explained variance for the first 70 components calculated by principal component analysis (black) and ICA (blue). Cumulative explained variance using only significant genes in each i-modulon is shown in green. (e) Scatterplot of two independent components, illustrating statistical independence. Given any value of independent component 2, the distribution of values across independent component 1 will be nearly identical to the distribution displayed above the plot. Thus, the coefficient of a gene in independent component 1 does not depend on the coefficient of the gene in independent component 2. The plot also illustrates that a gene may be significant in multiple components, and therefore a member of multiple i-modulons. (f) Schematic illustration of data processing pipeline. Validation steps (4 and 5) are discussed in results section “Validation of I-modulon-Regulator Relationships”.

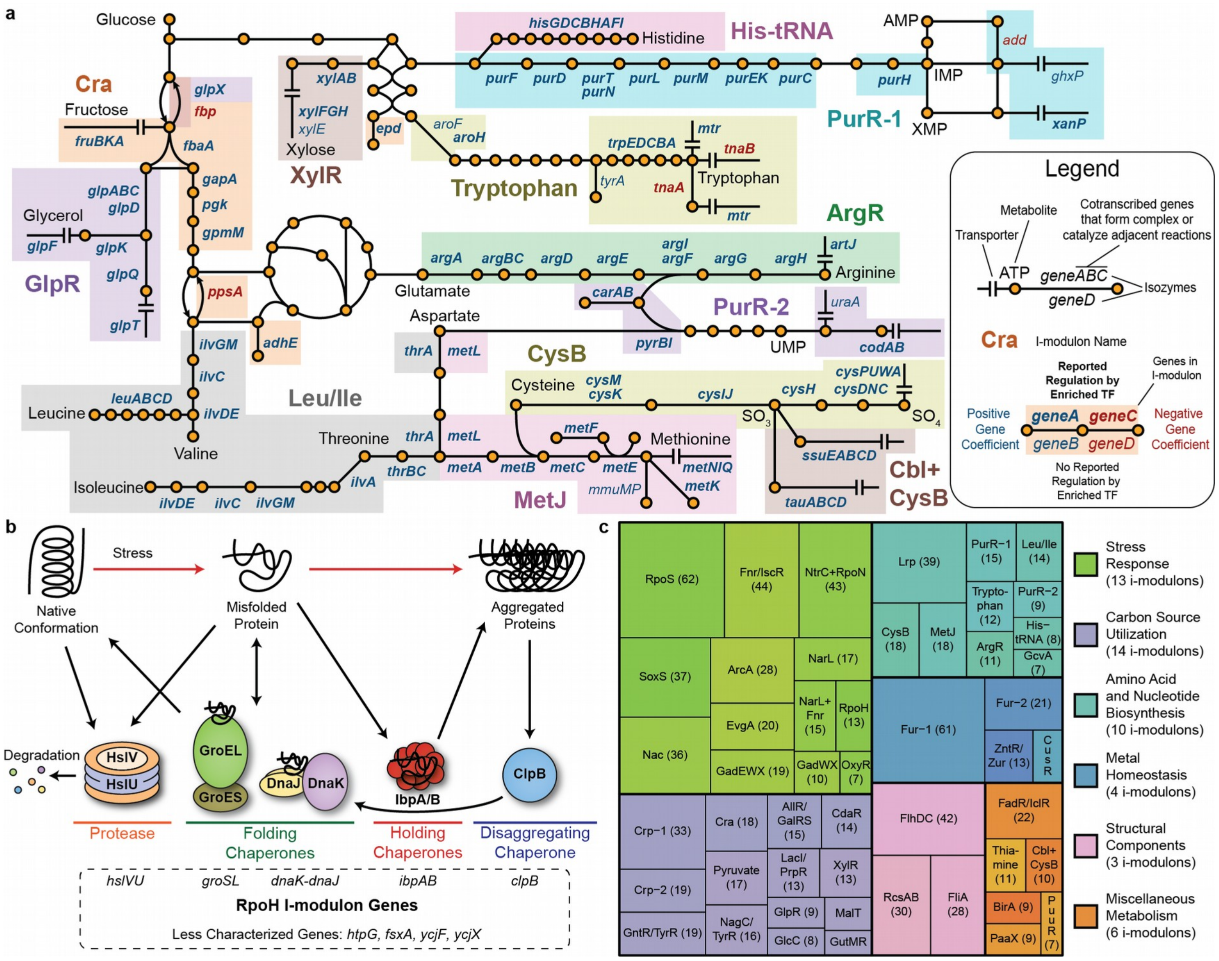

**Fig. S2: Overview of i-modulons**

(a) Schematic illustration of metabolic pathways captured by 12 regulatory i-modulons. Gene names within a colored box are members of the corresponding i-modulon. Co-transcribed genes whose products form a complex or catalyze adjacent reactions share gene names (e.g., trpEDCBA), and isozymes (i.e., enzymes that catalyze identical reactions) share reaction lines (e.g., glpABC/glpD). Genes with positive i-modulon weights are colored blue, and genes with negative weights are red. Bold font indicates that the genes belong to the associated regulon. (b) Simplified schematic illustration of the protein misfolding response in *E. coli* emphasizing genes in the RpoH i-modulon. Adapted from Baneyx and Mujacic, 2004[73]. (c) Illustration of the 50 regulatory i-modulons, colored by functional category. Number of genes in each i-modulon are indicated.

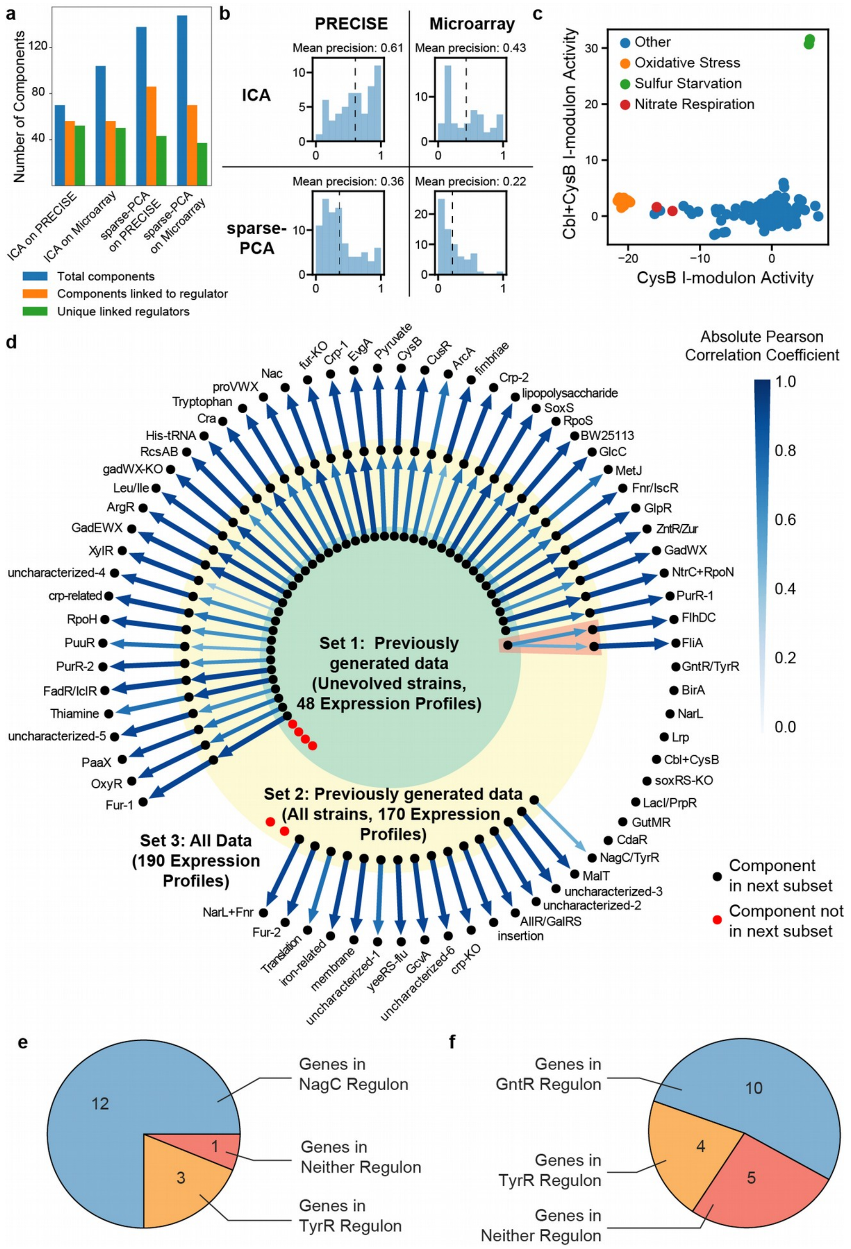


**Fig. S3: Validation of i-modulon regulator relationships**

(a) Evaluation of ICA and sparse-PCA on PRECISE and a microarray dataset (NCBI GEO GSE6836)[22]. Although ICA of PRECISE identifies the fewest number of components, almost all of these components are linked to regulators. In addition, ICA of PRECISE identifies more unique regulators than any other compared decomposition. (b) ICA of PRECISE reveals the strongest associations between regulators and components. (c) Scatterplot of CysB i-modulon activities against Cbl+CysB i-modulon activities. The Cbl+CysB i-modulon was only active under sulfur starvation conditions. (d) Comparison of ICA results on three subsets of the PRECISE dataset. Each node represents a component. Components are linked if their gene coefficients are correlated (Pearson R > 0.5). Red nodes were lost upon addition of new data. New data may also increase the resolution of previously identified components, such as splitting the chemotaxis-related FliA i-modulon from the flagella-related FlhDC i-modulon (pink box). (e) Distribution of gene regulation in the combined NagC/TyrR i-modulon. (f) Distribution of gene regulation in the combined GntR/TyrR i-modulon.

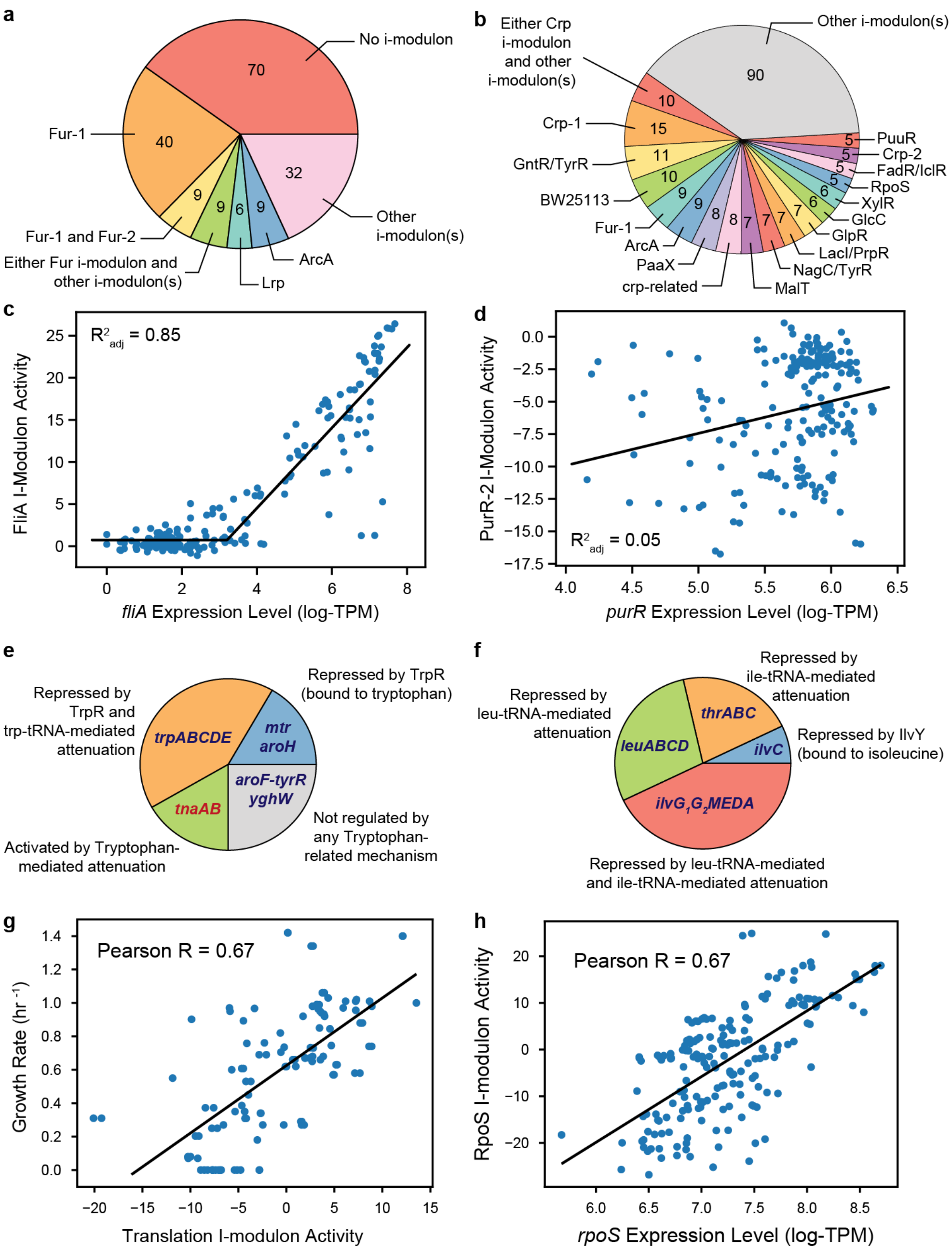


**Fig. S4: Additional properties of i-modulons**

(a,b) I-modulon composition of the reported Fur and Crp regulons. 353 genes in the reported Crp regulon were not in any i-modulon and are not shown. (c) The FliA i-modulon activity level was highly correlated with the log-transformed *fliA* expression level above a certain expression level. (d) Unlike the PurR-1 i-modulon, the PurR-2 i-modulon activity level was not correlated with *purR* expression. (e) Distribution of tryptophan-related regulatory mechanisms in the TrpR i-modulon. Gene names in blue indicate positive gene coefficients, and gene names in red indicate negative gene coefficients. (f) Distribution of leucine- and isoleucine-related regulatory mechanisms in the Leu/Ile i-modulon. High concentrations of leucine or isoleucine either charge tRNAs that pause transcription to form termination loops, or activate the transcription factor IlvY to repress transcription of *ilvC*. Subscripts indicate pseudogene fragments. (g) The translation i-modulon activity level was correlated with growth rate. (h) The RpoS i-modulon activity level was correlated with *rpoS* gene expression.

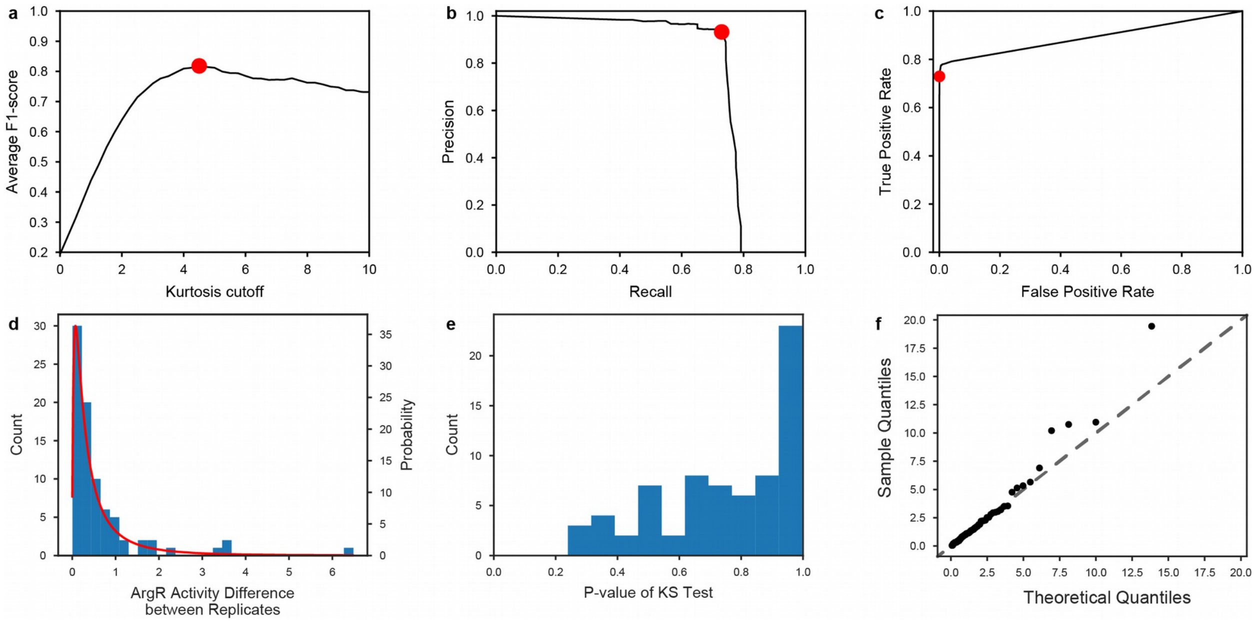


**Fig. S5: Significance thresholds for components and differential activity**

(a) Sensitivity analysis on the kurtosis cutoff. Red dot indicates optimal value. (b) Precision-recall curve and (c) receiver operating characteristic (ROC) curve from varying the kurtosis cutoff. (d) The difference in ArgR i-modulon activities between biological replicates follows a log-normal distribution (shown in red). (e) P-values from K-S tests on each i-modulon are higher than 0.1, indicating that an inability to reject the null hypothesis that the activity differences between replicates are from a log-normal distribution. (f) Quantile-quantile plot of difference in ArgR i-modulon activities between biological replicates provides further evidence of log-normal distribution.

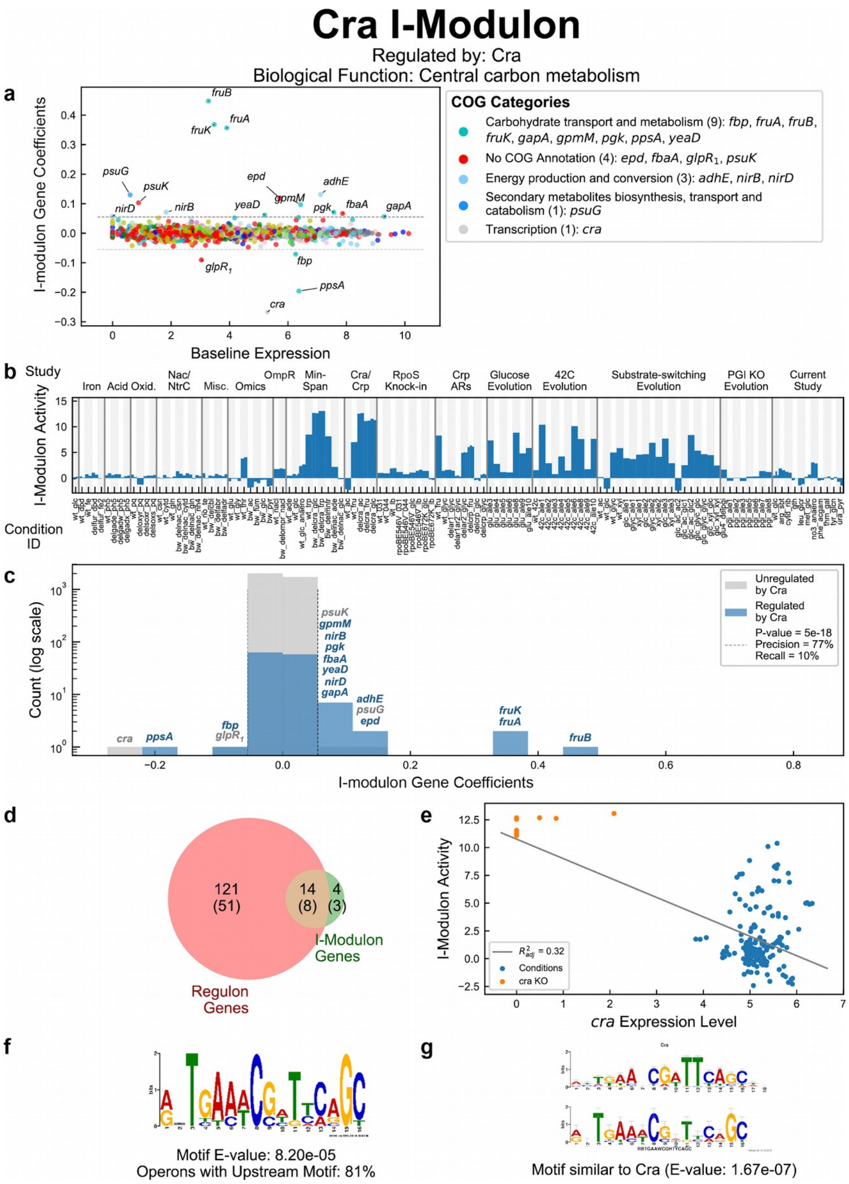


**Fig. S6: Descriptive Characteristics of the Cra I-modulon with Explanations**

Similar characteristics are shown for 30 selected i-modulons in Supplemental Dataset 2. (a) Scatterplot of i-modulon gene coefficients against gene expression (log-TPM) in the baseline condition (*E. coli* MG1655 grown on glucose M9 minimal medium). Genes are colored by their cluster of orthologous groups (COG) categories. (b) I-modulon activities across the entire PRECISE compendium, grouped by original study. Supplemental Dataset 1 contains experimental metadata (e.g. carbon source, pH, growth rate, GEO accession, etc.) for each condition ID. Each condition occupies a constant width, regardless of the number of biological replicates. (c) Histogram of i-modulon gene coefficients on a logarithmic count scale. Two histograms are overlaid, one for genes with reported regulation by the associated regulator, and one for genes without reported regulation. If multiple regulators are associated, additional histograms are overlaid. Gene names are listed above bars outside of the significance threshold (dashed vertical line). Statistics on regulator enrichment are reported in the legend. (d) Venn diagram of genes in associated regulon vs. significant genes in i-modulon. Numbers in parenthesis indicate number of operons in each category. (e) Scatterplot of i-modulon activity against expression level of associated regulator. Regulator knock-outs, if applicable, are indicated in orange. The $R^2_{adj}$ accounts for minimum expression levels, as discussed in Fig. S4c and Methods. (f) Motif identified upstream of i-modulon genes. (g) Similar motif in RegulonDB to the motif identified in (f).

### Supplemental Tables:

#### Table S1: ChIP-exo binding sites for MetJ

| Peak Name | Binding peak location | Peak Intensity | Transcription Unit (+ strand) | Gene Loci (+ strand) | Transcription Unit (- strand) | Gene Loci (- strand) |
|---|---|---|---|---|---|---|
| MetJ-1 | 83550 | 35 | | | leuLABCD | b0071;b0072;b0073;b0074;b0075 |
| MetJ-2 | 179560 | 23 | dgt | b0160 | mtn-btuF-yadS | b0157;b0158;b0159 |
| MetJ-3 | 222715 | 11 | gmhB | b0200 | metNIQ | b0197;b0198;b0199 |
| MetJ-4 | 275303 | 56 | mmuPM | b0260;b0261 | insH-1 | b0259 |
| MetJ-5 | 633492 | 63 | ybdL | b0600 | ybdH | b0599 |
| MetJ-6 | 1154392 | 27 | pabC-mltG-tmk-holB-ycfH | b1096;b1097;b1098;b1099;b1100 | | |
| MetJ-7 | 1398937 | 34 | | | uspE | b1333 |
| MetJ-8 | 1459265 | 43 | paaABCDEFGHIJK | b1388;b1389;b1390;b1391;b1392;b1393;b1394;b1395;b1396;b1397;b1398 | | |
| MetJ-9 | 1610723 | 36 | | | uxaB | b1521 |
| MetJ-10 | 1632086 | 27 | | | ydfJ_2 | b4600_2 |
| MetJ-11 | 1714512 | 17 | gstA | b1635 | | |
| MetJ-12 | 2132965 | 26 | | | wza-wzb-wzc-wcaAB | b2058;b2059;b2060;b2061;b2062 |
| MetJ-13 | 2154128 | 37 | mdtABCD-baeSR | b2074;b2075;b2076;b2077;b2078;b2079 | ibsB | b4668 |
| MetJ-14 | 2563581 | 46 | yffQR | b2448;b2449 | | |
| MetJ-15 | 2564631 | 29 | yffS | b2450 | | |
| MetJ-16 | 3086219 | 18 | yqgC;metK | b2940;b2942 | speAB;yqgB | b2937;b2938;b2939 |
| MetJ-17 | 3152198 | 78 | metC | b3008 | exbBD | b3005;b3006 |
| MetJ-18 | 3243319 | 39 | | | uxaCA | b3091;b3092 |
| MetJ-19 | 3353095 | 58 | | | arcB | b3210 |
| MetJ-20 | 3458579 | 49 | gspCDEFGHIJKLMO | b3324;b3325;b3326;b3327;b3328;b3329;b3330;b3331;b3332;b3333;b3334;b3335 | | |
| MetJ-21 | 3492425 | 67 | tsgA | b3364 | ppiA | b3363 |
| MetJ-22 | 3719905 | 39 | cspA | b3556 | | |
| MetJ-23 | 3891520 | 105 | mdtL | b3710 | | |
| MetJ-24 | 4011713 | 126 | | | | |
| MetJ-25 | 4012917 | 76 | metE | b3829 | metR | b3828 |
| MetJ-26 | 4128305 | 119 | metBL | b3939;b3940 | yiiX;metJ | b3937;b3938 |
| MetJ-27 | 4128579 | 105 | metBL | b3939;b3940 | metJ | b3938 |
| MetJ-28 | 4132531 | 98 | metF | b3941 | | |
| MetJ-29 | 4214175 | 75 | metA | b4013 | yjaB | b4012 |
| MetJ-30 | 4496913 | 26 | intB | b4271 | | |
| MetJ-31 | 4572359 | 38 | yjiT | b4342 | | |

### Table S2: ChIP-exo binding sites for CysB

| Peak Name | Binding peak location | Peak Intensity | Transcription Unit (+ strand) | Gene Loci (+ strand) | Transcription Unit (- strand) | Gene Loci (- strand) |
|---|---|---|---|---|---|---|
| CysB-1 | 50380 | 25 | | | | |
| CysB-2 | 70016 | 15 | araC | b0064 | araBAD | b0061;b0062;b0063 |
| CysB-3 | 113246 | 22 | guaC | b0104 | coaE-zapD-yacG | b0101;b0102;b0103 |
| CysB-4 | 135689 | 28 | | | yacC-speED | b0120;b0121;b0122 |
| CysB-5 | 169892 | 120 | fhuACDB | b0150;b0151;b0152;b0153 | | |
| CysB-6 | 252209 | 229 | dinB-yafNOP | b0231;b0232;b0233;b0234 | | |
| CysB-7 | 385122 | 158 | tauABCD | b0365;b0366;b0367;b0368 | | |
| CysB-8 | 401038 | 216 | iraP | b0382 | ddlA | b0381 |
| CysB-9 | 431231 | 58 | | | | |
| CysB-10 | 451813 | 168 | | | cyoABCDE | b0428;b0429;b0430;b0431;b0432 |
| CysB-11 | 516238 | 34 | fetAB | b0490;b0491 | qmcA-ybbJ | b0488;b0489 |
| CysB-12 | 609875 | 27 | | | fepA-entD | b0583;b0584 |
| CysB-13 | 754783 | 34 | sdhCDAB-sucABCD | b0721;b0722;b0723;b0724;b0726;b0727;b0728;b0729 | gltA | b0720 |
| CysB-14 | 815328 | 26 | | | | |
| CysB-15 | 850236 | 23 | ompX | b0814 | rhtA | b0813 |
| CysB-16 | 866398 | 235 | iaaA-gsiABCD | b0828;b0829;b0830;b0831;b0832 | moeAB | b0826;b0827 |
| CysB-17 | 904593 | 26 | ybjQ-amiD | b0866;b0867 | ybjP | b0865 |
| CysB-18 | 943784 | 52 | dmsABC | b0894;b0895;b0896 | | |
| CysB-19 | 997106 | 227 | | | ssuEADCB | b0933;b0934;b0935;b0936;b0937 |
| CysB-20 | 1053409 | 23 | | | yccM | b0992 |
| CysB-21 | 1098910 | 33 | ycdXY | b1034;b1035 | | |
| CysB-22 | 1165389 | 12 | hinT-ycfL-lpoB-thiK-nagZ-ycfP | b1103;b1104;b1105;b1106;b1107;b1108 | | |
| CysB-23 | 1195322 | 14 | icd | b1136 | rluE-nudJ | b1134;b1135 |
| CysB-24 | 1250937 | 32 | dhaR | b1201 | dhaKLM | b1198;b1199;b1200 |
| CysB-25 | 1333725 | 39 | cysB | b1275 | | |
| CysB-26 | 1350219 | 68 | | | yciW | b1287 |
| CysB-27 | 1526120 | 273 | yncG | b1454 | ansP | b1453 |
| CysB-28 | 1555778 | 15 | | | maeA | b1479 |
| CysB-29 | 1567237 | 117 | | | dosCP | b1489;b1490 |
| CysB-30 | 1600329 | 25 | | | lsrRK | b1511;b1512 |
| CysB-31 | 1667332 | 6 | | | mlc-ynfK | b1593;b1594 |
| CysB-32 | 1706881 | 12 | ydgK-rsxABCDGE-nth | b1626;b1627;b1628;b1629;b1630;b1631;b1632;b1633 | | |
| CysB-33 | 1810832 | 58 | ydjN | b1729 | | |
| CysB-34 | 1818923 | 13 | | | chbBCARFG | b1733;b1734;b1735;b1736;b1737;b1738 |
| CysB-35 | 1842359 | 41 | gdhA | b1761 | ynjH | b1760 |
| CysB-36 | 1924627 | 44 | holE | b1842 | yobA-yebZY | b1839;b1840;b1841 |
| CysB-37 | 1977895 | 273 | | | flhDC | b1891;b1892 |
| CysB-38 | 2032198 | 12 | | | dcm-vsr | b1960;b1961 |
| CysB-39 | 2060953 | 122 | | | cbl | b1987 |
| CysB-40 | 2084561 | 69 | | | yeeED | b2012;b2013 |
| CysB-41 | 2085611 | 113 | | | yeeED | b2012;b2013 |
| CysB-42 | 2110587 | 12 | | | rfbBDACX | b2037;b2038;b2039;b2040;b2041 |
| CysB-43 | 2133482 | 17 | | | wza-wzb-wzc-wcaAB | b2058;b2059;b2060;b2061;b2062 |
| CysB-44 | 2192823 | 32 | | | yehE | b2112 |
| CysB-45 | 2240873 | 21 | | | | |
| CysB-46 | 2249630 | 26 | yeiH | b2158 | yeiE | b2157 |
| CysB-47 | 2413410 | 33 | ackA-pta | b2296;b2297 | yfbV | b2295 |
| CysB-48 | 2434800 | 29 | | | pdxB-usg-truA-dedA | b2317;b2318;b2319;b2320 |
| CysB-49 | 2489195 | 9 | | | yfdX-frc-oxc-yfdVE | b2371;b2372;b2373;b2374;b2375 |
| CysB-50 | 2532389 | 194 | cysK | b2414 | | |
| CysB-51 | 2543583 | 227 | | | cysPUWAM | b2421;b2422;b2423;b2424;b2425 |

| | | | | | | |
|---|---|---|---|---|---|---|
| CysB-52 | 2551298 | 36 | | | yfeYX;ypeA-yfeZ | b2431;b2432;b2433;b2434 |
| CysB-53 | 2563572 | 21 | yffQR | b2448;b2449 | | |
| CysB-54 | 2616204 | 11 | bepA-yfgD | b2494;b2495 | yfgO | b2493 |
| CysB-55 | 2630311 | 36 | yfgHI | b2505;b2506 | | |
| CysB-56 | 2876380 | 287 | iap | b2753 | cysDNC | b2750;b2751;b2752 |
| CysB-57 | 2891912 | 189 | queD | b2765 | cysJIH | b2762;b2763;b2764 |
| CysB-58 | 2971189 | 26 | ygdR;tas | b2833;b2834 | | |
| CysB-59 | 3029593 | 8 | | | | |
| CysB-60 | 3263874 | 9 | | | tdcABCDEFG | b4471;b3113;b3114;b3115;b3116;b3117;<br>b3118 |
| CysB-61 | 3354689 | 15 | gltBDF | b3212;b3213;b3214 | | |
| CysB-62 | 3367136 | 19 | yhcE_2F | b4569_2;b3219 | insH-10 | b3218 |
| CysB-63 | 3373600 | 11 | | | nanATEK-yhcH | b3221;b3222;b3223;b3224;b3225 |
| CysB-64 | 3384357 | 42 | argR | b3237 | mdh | b3236 |
| CysB-65 | 3418320 | 15 | yhdV | b3267 | | |
| CysB-66 | 3558468 | 9 | rtcR | b3422 | rtcBA | b4475;b3421 |
| CysB-67 | 3712010 | 23 | | | yhjX | b3547 |
| CysB-68 | 3828710 | 32 | xanP | b3654 | gltS | b3653 |
| CysB-69 | 3859989 | 39 | yidL | b3680 | yidKJ | b3678;b3679 |
| CysB-70 | 3839993 | 41 | yicS | b4555 | nlpA | b3661 |
| CysB-71 | 3867710 | 17 | yidQ | b3688 | ibpAB | b3686;b3687 |
| CysB-72 | 3891605 | 293 | mdtL | b3710 | | |
| CysB-73 | 4077596 | 8 | yihXY-dtd-yiiD | b3885;b3886;b3887;b3888 | | |
| CysB-74 | 4092321 | 18 | | | frvABXR | b3897;b3898;b3899;b3900 |
| CysB-75 | 4108796 | 195 | sbp | b3917 | | |
| CysB-76 | 4214123 | 86 | metA | b4013 | yjaB | b4012 |
| CysB-77 | 4270438 | 8 | yjbQR | b4056;b4057 | | |
| CysB-78 | 4337654 | 5 | | | adiY | b4116 |
| CysB-79 | 4556320 | 26 | | | yjiC | b4325 |

### Table S3: Computationally-detected novel TF binding sites upstream of i-modulon genes

| Identified Motif[1] | I-modulon | Operon | Locus tag(s) | Site p-value | Site start position | Site sequence |
|---|---|---|---|---|---|---|
| 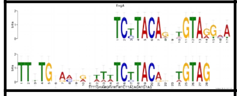 | EvgA | ypdI | b2376 | 3.37E-15 | 2494486 | TTTTGAAAGGGTTATCTTACAG TTGTAG |
| 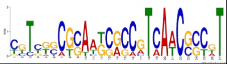 | FlhDC | fliC | b1923 | 3.33E-13 | 2003656 | CGTCGGCTCAATCGCCGTCAAC CCTGT |
| 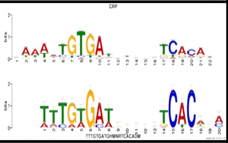 | Crp-1 | nmpC | b0553_1 | 1.61E-06 | 576933 | TTTGTGAAGTAGATCTCTAT |
| | | pepE | b4021 | 3.56E-06 | 4230350 | TTTATGCTGCCCGACTCATC |
| | | ydeNM | b1498 | 1.04E-07 | 1582801 | TATGTGATGGATGTCACTTA |
| | | ymfED | b1138 | 6.78E-07 | 1198360 | ATTGTGATCAGTAGCACGTA |
| 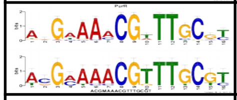 | PurR-1 | ghxP | b4064 | 1.14E-08 | 4278206 | ACGATAACGTTTGCGC |
| 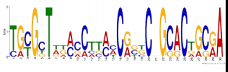 | CdaR | srlAEBD-gutM-srlR-gutQ | b2703,b2705,b2706,b 2702,b2704 | 2.01E-10 | 2825611 | TATGGTAAAGCATCACGCCCC GCACAAGGA |
| 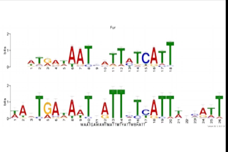 | Fur-2 | bfd-bfr | b3337 | 2.57E-11 | 3467020 | AAATGAAAATAGTTCTTATTTC AATT |
| | | preTA | b2146 | 9.14E-08 | 2233859 | TATTGAGAATAATTATTACTTC ACCT |
| | | ydhYVWXUT | b1674,b1673,b1672 | 1.82E-09 | 1754721 | AACTGATATTTATTATCATTTG AAAT |
| 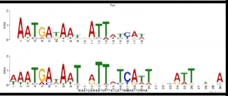 | Fur-1 | bfd-bfr | b3336,b3337 | 5.55E-09 | 3467020 | AAATGAAAATAGTTCTTATTTC AATTACGG |
| 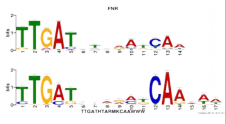 | Fnr/IscR | ghoST | b4128 | 6.83E-05 | 4352566 | TTGATAGGAGTCATATT |
| | | iscRSUA | b2531,b2530 | 1.19E-04 | 2662361 | CTGATAAGACGCATTAC |
| | | ttdR | b3060 | 6.19E-06 | 3206313 | CTGCTCTGGCGCAATAT |
| | | ycbJ | b0919 | 2.10E-05 | 971518 | TTGCTGTTGCTCAGGAA |
| | | yhgH-nfuA | b3414 | 6.39E-05 | 3545505 | TTGCTTTTACGCAATGG |
| | | yoeA_1 | b4582_1 | 2.10E-05 | 2068504 | ATGATTAAGGTCAAAAA |
| | | yqhD-dkgA | b3011 | 1.68E-06 | 3155193 | TTGATTGTGCGCAATTT |
| 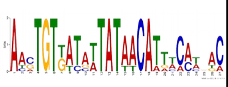 | ZntR/Zur | mepM | b1856 | 6.66E-10 | 1941849 | AACTGTGTTCTTATTTCATGTA AACGC |
| | | pliG | b1178 | 2.81E-12 | 1227503 | AACTGTTATAATATAACAATCC CTAAC |
| 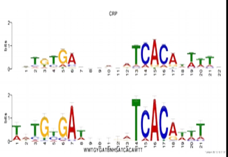 | crp-related | fliLMNOPQR | b1949 | 1.73E-05 | 2022414 | TTTCAGATAGGCTTCACGATT |
| | | mscS | b2924 | 2.86E-06 | 3069899 | TTTGCCAAATAGATCACAGAT |
| | | nepI | b3662 | 8.55E-06 | 3841792 | TGTGTGACGCATTTAACGTTT |
| | | ybiJ | b0802 | 8.74E-07 | 837972 | GATGTGATGAGTATCACGTTT |
| | | ydeA | b1528 | 5.78E-09 | 1616934 | AACGCGATCCAGATCACAAAT |
| | | ydhP | b1657 | 2.19E-05 | 1737392 | TTTGCCATTTTGCTAACAAAC |
| | | yedRJ | b1963 | 1.83E-05 | 2033750 | AATGTGCGCCTGATCACACCA |
| | | yjcB | b4060 | 9.23E-09 | 4275169 | TTTGTGAACTATATCACAATT |
| | | yneE | b1520 | 1.13E-07 | 1609056 | AACGTGATTACGATCACATTC |
| 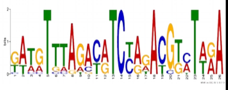 | MetJ | ybdH | b0599 | 1.48E-11 | 633496 | TTTATTTAGACATCTAAACGTC TTGA |
| | | metJ | b3938 | 6.14E-12 | 4128577 | GAAGTTTAGATGTCCAGATGTA TTGA |
| | | mmuPM | b0261,b0260 | 1.08E-11 | 275291 | GATGTTTAGATGTCCATACGTT TAGA |
| | | ybdL | b0600 | 1.48E-11 | 633496 | TTTATTTAGACATCTAAACGTC TTGA |
| 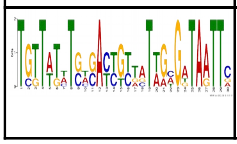 | XylR | xylE | b4031 | 1.81E-11 | 4242357 | TCTTTTCTGTGATCTTAATTGTG ATAATTA |
| | | yiaB | b3563 | 1.74E-11 | 3728000 | TGGTAGATGCGACTGTTCTAAC GGTAGTTG |
| | | yicJI | b3656,b3657 | 5.56E-13 | 3836034 | TGTTTTATGTGATCGTGGTAGC |

| | | | | | | |
|---|---|---|---|---|---|---|
| | | | | | | GTTAATTC |
| 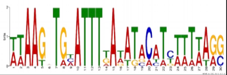 | NarL | hmp | b2552 | 3.69E-13 | 2685783 | ATAAGATGCATTTGAGATACATCAATTAAG |
| 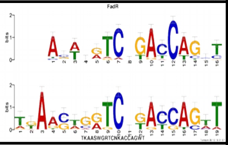 | FadR/IclR | ybfA | b0699 | 4.71E-07 | 729018 | ATAAGTGGTCGGATGAGTA |
| | | yqaE-kbp | b2666 | 7.49E-07 | 2797094 | TTAGCGGAGCCTGCCAGTT |
| 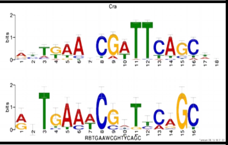 | Cra | glpEGR | b3423_1 | 2.50E-06 | 3560722 | AATCAAACCATCCGGC |
| 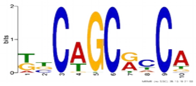 | RpoS | adhP | b1478 | 2.55E-06 | 1554161 | TTCAGCACCA |
| | | ahr | b4269 | 1.38E-04 | 4496479 | TTCGGCACCA |
| | | amyA | b1927 | 1.21E-04 | 2005911 | TGCTGCGCCT |
| | | bfd-bfr | b3336 | 3.81E-04 | 3466818 | GGCAGCTGCA |
| | | ggt | b3447 | 2.68E-04 | 3587285 | ATCAGCAGCA |
| | | yahK | b0325 | 4.81E-04 | 342521 | GCCTGCAACG |
| | | yahO | b0329 | 2.38E-05 | 346024 | TGCTGCACCA |
| | | ybaY | b0453 | 7.56E-05 | 475461 | GCCAGCACCG |
| | | ybdK | b0581 | 9.73E-05 | 607591 | TTCAGCTTCA |
| | | ybhP-clsB-ybhN | b0788,b0789,b0790 | 8.40E-07 | 824801 | TTCAGCGCCA |
| | | ycaC | b0897 | 3.84E-05 | 945515 | GGCAGCATCA |
| | | ycgB | b1188 | 4.08E-06 | 1237775 | TCCAGCGCCA |
| | | ydhS | b1668 | 4.81E-04 | 1747191 | TACAGCAACA |
| | | yeaGH | b1784,b1783 | 3.22E-04 | 1866379 | GCCTGCGCCC |
| | | yebV | b1836 | 2.82E-05 | 1921555 | TCCAGCATCA |
| | | yedP | b1955 | 1.21E-04 | 2025027 | GTCAGCGTCG |
| | | yegP | b2080 | 5.74E-05 | 2164675 | TGCAGCAACA |
| | | yegS | b2086 | 1.30E-05 | 2168117 | TTCTGCACCA |
| | | ygaM | b2672 | 4.37E-04 | 2799630 | GGCATCGCCA |
| | | yghA | b3003 | 8.40E-07 | 3149373 | TTCAGCGCCA |
| | | yhbO | b3153 | 9.73E-05 | 3298670 | ACCTGCGCCA |
| | | yhcO | b3239 | 2.55E-06 | 3386494 | GTCAGCGCCA |
| | | ymgE | b1195 | 8.06E-06 | 1244366 | TCCAGCACCA |
| | | yqjCDEK | b3100 | 1.64E-05 | 3250016 | TGCTGCGCCA |
| | | ytjA | b4568 | 3.84E-05 | 4611737 | GTCAGCGACA |
| 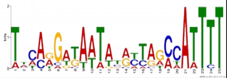 | Lrp | yqeG | b2845 | 1.38E-07 | 2985792 | TTTATGGTAAATTGCCCTCCATTTT |
| 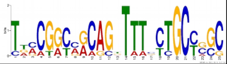 | GntR/TyrR | ptsG | b1101 | 4.83E-09 | 1157935 | TGCCTATCGCAGGTATTCTGCTGGG |

[1] Motifs were identified by searching all upstream sequences of genes in an i-modulon, and compared to reported motifs in RegulonDB (See Methods). If a match was found, the top motif is the RegulonDB motif, and the bottom motif is the motif found upstream of i-modulon genes.

**Table S4: I-modulon membership for uncharacterized genes as defined by Ghatak et al. 2018[72]. Additional annotations are derived from i-modulon membership.**

| b-number | Gene Name | I-modulon | Current annotation from Ecocyc | Additional annotation from i-modulon membership |
|---|---|---|---|---|
| b4662 | sgrT | AllR/GalRS | putative inhibitor of the PtsG glucose transporter | |
| b0975 | hyaD | ArcA | putative hydrogenase 1 maturation protease HyaD | |
| b0976 | hyaE | ArcA | putative HyaA chaperone | |
| b0977 | hyaF | ArcA | protein HyaF | |
| b1020 | phoH | ArcA | ATP-binding protein PhoH | |
| b2713 | hydN | ArcA | putative electron transport protein HydN | |
| b2725 | hycA | ArcA | regulator of the transcriptional regulator FhlA | |
| b1541 | ydfZ | ArcA, Fnr/IscR | putative selenoprotein YdfZ | |
| b1256 | ompW | ArcA, Fnr/IscR, BirA | outer membrane protein W | |
| b2788 | gudX | CdaR | glucarate dehydratase-related protein | |
| b2789 | gudP | CdaR | galactarate/glucarate/glycerate transporter GudP | |
| b1780 | yeaD | Cra | putative aldose 1-epimerase YeaD | |
| b2166 | psuK | Cra | putative pseudouridine kinase | |
| b1138 | ymfE | Crp-1 | e14 prophage; uncharacterized protein YmfE | |
| b1143 | ymfI | Crp-1 | e14 prophage; protein YmfI | |
| b1145 | cohE | Crp-1 | e14 prophage; putative repressor protein YmfK | |
| b1426 | ydcH | Crp-1 | protein YdcH | |
| b3358 | yhfK | Crp-1 | putative transporter YhfK | |
| b4357 | lgoR | Crp-1 | putative DNA-binding transcriptional regulator LgoR | |
| b1498 | ydeN | Crp-1, NagC/TyrR | putative sulfatase | |
| b0791 | ybhQ | Crp-2 | putative inner membrane protein | |
| b1423 | ydcJ | Crp-2 | DUF1338 domain-containing protein YdcJ | |
| b2535 | csiE | Crp-2 | stationary phase inducible protein CsiE | |
| b4068 | yjcH | Crp-2 | conserved inner membrane protein YjcH | |
| b1441 | ydcT | Crp-2, Nac | putative ABC transporter ATP-binding protein YdcT | |
| b1287 | yciW | CysB | putative oxidoreductase | Sulfur oxidoreductase |
| b1466 | narW | EvgA | NarW, putative private chaperone for NarZ nitrate reductase subunit | |
| b1501 | ydeP | EvgA | putative oxidoreductase YdeP | |
| b2082 | ogrK | EvgA | prophage P2 late control protein OgrK | |
| b2085 | yegR | EvgA | uncharacterized protein YegR | |
| b2367 | emrY | EvgA | tripartite efflux pump membrane subunit EmrY | |
| b2368 | emrK | EvgA | tripartite efflux pump membrane fusion protein EmrK | |
| b2372 | yfdV | EvgA | putative transport protein YfdV | |
| b2375 | yfdX | EvgA | protein YfdX | |
| b2376 | ypdI | EvgA | colanic acid synthesis putative lipoprotein YpdI | |
| b4518 | ymdF | EvgA | conserved protein YmdF | |
| b0557 | borD | FadR/IclR | DLP12 prophage; prophage lipoprotein BorD | Related to fatty acid degradation |
| b0699 | ybfA | FadR/IclR | DUF2517 domain-containing protein YbfA | Related to fatty acid degradation |
| b3466 | yhhL | FadR/IclR | DUF1145 domain-containing protein YhhL | Related to fatty acid degradation |
| b2666 | yqaE | FadR/IclR, insertion | Pmp3 family protein | Related to fatty acid degradation |
| b4315 | fimI | fimbriae | putative fimbrial protein FimI | |
| b1044 | ymdA | FlhDC | uncharacterized protein YmdA | Flagellar protein |
| b1072 | flgA | FlhDC | flagellar basal body P-ring formation protein FlgA | |
| b1075 | flgD | FlhDC | flagellar biosynthesis, initiation of hook assembly | |
| b1081 | flgJ | FlhDC | putative peptidoglycan hydrolase FlgJ | |
| b1878 | flhE | FlhDC | flagellar protein | |
| b1904 | yecR | FlhDC | lipoprotein YecR | Flagellar protein |
| b1943 | fliK | FlhDC | flagellar hook-length control protein | |
| b1070 | flgN | FlhDC, FliA | flagellar biosynthesis protein FlgN | |
| b1925 | fliS | FlhDC, FliA | flagellar biosynthesis protein FliS | |
| b1926 | fliT | FlhDC, FliA | flagellar biosynthesis protein FliT | |
| b1566 | flxA | FliA | Qin prophage; protein FlxA | Chemotaxis protein |
| b1742 | ves | FliA | HutD family protein Ves | Chemotaxis protein |
| b1760 | ynjH | FliA | DUF1496 domain-containing protein YnjH | Chemotaxis protein |
| b4110 | yjcZ | FliA | uncharacterized protein YjcZ | Chemotaxis protein |
| b0715 | abrB | Fnr/IscR | putative regulator | |
| b0836 | bssR | Fnr/IscR | regulator of biofilm formation | |

| | | | | |
|---|---|---|---|---|
| b0919 | ycbJ | Fnr/IscR | putative phosphotransferase YcbJ | |
| b1587 | ynfE | Fnr/IscR | putative selenate reductase YnfE | |
| b1593 | ynfK | Fnr/IscR | putative dethiobiotin synthetase | |
| b1750 | ydjX | Fnr/IscR | DedA family protein YdjX | |
| b1751 | ydjY | Fnr/IscR | 4Fe-4S ferredoxin-type domain-containing protein YdjY | |
| b1785 | yeaI | Fnr/IscR | putative c-di-GMP binding protein CdgI | |
| b1906 | yecH | Fnr/IscR | DUF2492 domain-containing protein YecH | |
| b3158 | yhbU | Fnr/IscR | putative peptidase YhbU | |
| b3159 | yhbV | Fnr/IscR | putative peptidase YhbV | |
| b3211 | yhcC | Fnr/IscR | radical SAM family oxidoreductase YhcC | |
| b0468 | ybaN | Fur-1 | conserved inner membrane protein YbaN | |
| b0587 | fepE | Fur-1 | polysaccharide co-polymerase family protein FepE | |
| b0803 | ybiI | Fur-1 | zinc finger domain-containing protein YbiI | |
| b0804 | ybiX | Fur-1 | PKHD-type hydroxylase YbiX | |
| b1452 | yncE | Fur-1 | PQQ-like domain-containing protein YncE | |
| b1494 | pqqL | Fur-1 | putative zinc peptidase | |
| b1496 | yddA | Fur-1 | ABC transporter family protein YddA | |
| b1705 | ydiE | Fur-1 | PF10636 family protein YdiE | |
| b4567 | yjjZ | Fur-1 | protein YjjZ | |
| b1672 | ydhW | Fur-2 | protein YdhW | |
| b1673 | ydhV | Fur-2 | putative oxidoreductase | |
| b1674 | ydhY | Fur-2, Fnr/IscR | putative 4Fe-4S ferredoxin-type protein | |
| b3337 | bfd | Fur-2, Fur-1 | bacterioferritin-associated ferredoxin | |
| b3410 | feoC | Fur-2, OxyR | ferrous iron transport protein FeoC | |
| b3362 | yhfG | GadEWX | DUF2559 domain-containing protein YhfG | Acid resistance protein |
| b3491 | yhiM | GadEWX | inner membrane protein with a role in acid resistance | |
| b3508 | yhiD | GadEWX | inner membrane protein YhiD | Acid resistance protein |
| b3511 | hdeD | GadEWX | acid-resistance membrane protein | |
| b4377 | yjjU | GadEWX | putative patatin-like phospholipase YjjU | Acid resistance protein |
| b3507 | dctR | GadEWX, GadWX | putative DNA-binding transcriptional regulator DctR | |
| b0486 | ybaT | GadWX | putative transporter YbaT | Acid resistance protein |
| b2902 | ygfF | GcvA | putative NAD(P)-binding oxidoreductase with NAD(P)-binding Rossmann-fold domain | |
| b3646 | yicG | GcvA | conserved inner membrane protein YicG | |
| b2977 | glcG | GlcC | putative heme-binding protein GlcG | |
| b2979 | glcD | GlcC | glycolate dehydrogenase, putative FAD-linked subunit | |
| b4467 | glcF | GlcC | glycolate dehydrogenase, putative iron-sulfur subunit | |
| b4468 | glcE | GlcC | glycolate dehydrogenase, putative FAD-binding subunit | |
| b4705 | mntS | GlcC | small protein MntS | |
| b2377 | yfdY | insertion | DUF2545 domain-containing protein YfdY | |
| b3691 | dgoT | LacI/PrpR | putative D-galactonate transporter | |
| b1137 | ymfD | lipopolysaccharide | e14 prophage; putative SAM-dependent methyltransferase | Lipopolysaccharide-related protein |
| b1550 | gnsB | lipopolysaccharide | Qin prophage; protein GnsB | Lipopolysaccharide-related protein |
| b2032 | wbbK | lipopolysaccharide | putative lipopolysaccharide biosynthesis protein | |
| b2035 | wbbH | lipopolysaccharide | putative O-antigen polymerase | |
| b2145 | yeiS | lipopolysaccharide | DUF2542 domain-containing protein YeiS | Lipopolysaccharide-related protein |
| b2642 | yfjW | lipopolysaccharide | CP4-57 prophage; uncharacterized protein YfjW | Lipopolysaccharide-related protein |
| b3618 | yibB | lipopolysaccharide | protein HtrL | Lipopolysaccharide-related protein |
| b4253 | yjgL | lipopolysaccharide | protein YjgL | Lipopolysaccharide-related protein |
| b4279 | yjhB | lipopolysaccharide | putative sialic acid transporter | |
| b2352 | gtrS | Lipopolysaccharide, GcvA | CPS-53 (KpLE1) prophage; serotype specific glucosyl transferase | |
| b1534 | ydeE | Lrp | dipeptide exporter | |
| b1605 | ydgI | Lrp | putative arginine:ornithine antiporter | |
| b2845 | yqeG | Lrp | putative transporter YqeG | Amino acid or peptide transporter |
| b4045 | yjbJ | Lrp | putative stress response protein | |
| b4217 | ytfK | Lrp | DUF1107 domain-containing protein YtfK | |
| b3523 | yhjE | Lrp, Nac | putative transporter YhjE | Amino acid or peptide transporter |
| b4037 | malM | MalT | maltose regulon periplasmic protein | |
| b1828 | yebQ | membrane | putative transporter YebQ | |
| b2971 | yghG | membrane | lipoprotein YghG | |
| b2972 | pppA | membrane | prepilin peptidase | |
| b3151 | yraQ | membrane | permease family protein YraQ | |

| | | | | |
|---|---|---|---|---|
| b4466 | sslE | membrane | putative lipoprotein YghJ | |
| b2012 | yeeD | membrane, CysB | putative sulfurtransferase YeeD | |
| b2013 | yeeE | membrane, CysB | inner membrane protein YeeE | |
| b1495 | yddB | membrane, Fur-1 | putative TonB-dependent receptor | |
| b3937 | yiiX | MetJ | putative lipid binding hydrolase | Methionine or folate related hydrolase |
| b1339 | abgR | Nac | putative LysR-type DNA-binding transcriptional regulator AbgR | |
| b1375 | ynaE | Nac | Rac prophage; uncharacterized protein YnaE | Nitrogen starvation protein |
| b1442 | ydcU | Nac | putative ABC transporter membrane subunit YdcU | Putative peptide transporter |
| b1443 | ydcV | Nac | putative ABC transporter membrane subunit YdcV | Putative peptide transporter |
| b3043 | ygiL | Nac | putative fimbrial protein YgiL | Nitrogen starvation protein |
| b3446 | yrhB | Nac | putative heat shock chaperone | Nitrogen starvation protein |
| b3712 | yieE | Nac | putative phosphopantetheinyl transferase | Nitrogen starvation protein |
| b1497 | ydeM | NagC/TyrR | putative anaerobic sulfatase maturation enzyme YdeM | |
| b1796 | yoaG | NarL | DUF1869 domain-containing protein YoaG | |
| b1552 | cspI | NarL+Fnr | Qin prophage; cold shock protein CspI | |
| b2208 | napF | NarL+Fnr, Fnr/IscR | ferredoxin-type protein | |
| b1797 | yeaR | NarL+Fnr, NarL | DUF1971 domain-containing protein YeaR | |
| b4506 | ykgO | NarL+Fnr, ZntR/Zur | putative ribosomal protein | |
| b1008 | rutE | NtrC+RpoN | putative malonic semialdehyde reductase | |
| b1009 | rutD | NtrC+RpoN | putative aminoacrylate hydrolase | |
| b1010 | rutC | NtrC+RpoN | putative aminoacrylate peracid reductase | |
| b1483 | ddpF | NtrC+RpoN | putative D,D-dipeptide ABC transporter ATP-binding subunit DdpF | |
| b1484 | ddpD | NtrC+RpoN | putative D,D-dipeptide ABC transporter ATP-binding subunit DdpD | |
| b1485 | ddpC | NtrC+RpoN | putative D,D-dipeptide ABC transporter membrane subunit DdpC | |
| b1486 | ddpB | NtrC+RpoN | putative D,D-dipeptide ABC transporter membrane subunit DdpB | |
| b1487 | ddpA | NtrC+RpoN | putative D,D-dipeptide ABC transporter periplasmic binding protein | |
| b1596 | ynfM | NtrC+RpoN | putative transporter YnfM | Putative peptide transporter |
| b2870 | ygeW | NtrC+RpoN | putative carbamoyltransferase YgeW | Nitrogen starvation protein |
| b2875 | yqeB | NtrC+RpoN | XdhC-CoxI family protein YqeB | Nitrogen starvation protein |
| b2876 | yqeC | NtrC+RpoN | uncharacterized protein YqeC | Nitrogen starvation protein |
| b3269 | yhdX | NtrC+RpoN | putative ABC transporter membrane subunit YhdX | Putative peptide transporter |
| b3270 | yhdY | NtrC+RpoN | putative ABC transporter membrane subunit YhdY | Putative peptide transporter |
| b3271 | yhdZ | NtrC+RpoN | putative ABC transporter ATP-binding subunit YhdZ | Putative peptide transporter |
| b1932 | yedL | NtrC+RpoN, Nac | putative acetyltransferase YedL | Nitrogen starvation protein |
| b1389 | paaB | PaaX | phenylacetyl-CoA 1,2-epoxidase subunit B | |
| b1391 | paaD | PaaX | phenylacetate degradation protein | |
| b1660 | ydhC | PurR-1 | putative transporter YdhC | Purine/purine precursor efflux pump |
| b2313 | cvpA | PurR-1 | colicin V production protein | Purine biosynthetic protein |
| b0306 | ykgE | Pyruvate | putative lactate utilization oxidoreductase YkgE | |
| b2141 | yohJ | Pyruvate | PF03788 family membrane protein YohJ | Pyruvate-responsive protein |
| b3547 | yhjX | Pyruvate | putative pyruvate transporter | |
| b3716 | cbrB | Pyruvate | putative inner membrane protein | Pyruvate-responsive protein |
| b4353 | yjiX | Pyruvate | conserved protein YjiX | Pyruvate-responsive protein |
| b0527 | ybcI | RcsAB | conserved inner membrane protein YbcI | |
| b2044 | wcaL | RcsAB | putative colanic biosynthesis glycosyl transferase | |
| b2045 | wcaK | RcsAB | putative colanic acid biosynthesis pyruvyl transferase | |
| b2046 | wzxC | RcsAB | G7097-MONOMER | |
| b2050 | wcaI | RcsAB | putative colanic biosynthesis glycosyl transferase | |
| b2054 | wcaF | RcsAB | putative acyl transferase | |
| b2055 | wcaE | RcsAB | putative colanic acid biosynthesis glycosyl transferase | |
| b2056 | wcaD | RcsAB | putative colanic acid polymerase | |
| b2057 | wcaC | RcsAB | putative colanic acid biosynthesis glycosyl transferase | |
| b2058 | wcaB | RcsAB | putative colanic acid biosynthesis acyl transferase | |
| b2059 | wcaA | RcsAB | putative colanic acid biosynthesis glycosyl transferase | |
| b4026 | yjbE | RcsAB | uncharacterized protein YjbE | |
| b4027 | yjbF | RcsAB | lipoprotein YjbF | |
| b4028 | yjbG | RcsAB | capsule biosynthesis GfcC family protein YjbG | |

| | | | | |
|---|---|---|---|---|
| b4029 | yjbH | RcsAB | YjbH family protein | Colanic acid-related protein |
| b3563 | yiaB | RcsAB, XylR | conserved inner membrane protein YiaB | Putative xylose or xyloside transporter |
| b1321 | ycjX | RpoH | DUF463 domain-containing protein YcjX | Misfolding response |
| b1322 | ycjF | RpoH | conserved inner membrane protein YcjF | Misfolding response |
| b4140 | fxsA | RpoH | protein FxsA | Misfolding response |
| b0329 | yahO | RpoS | DUF1471 domain-containing protein YahO | Stress-related protein |
| b0453 | ybaY | RpoS | PF09619 family lipoprotein YbaY | Stress-related protein |
| b0707 | ybgA | RpoS | DUF1722 domain-containing protein YbgA | Stress-related protein |
| b0788 | ybhN | RpoS | conserved inner membrane protein YbhN | Stress-related protein |
| b0790 | ybhP | RpoS | endonuclease/exonuclease/phosphatase domain-containing protein YbhP | Stress-related protein |
| b0897 | ycaC | RpoS | putative hydrolase | Stress-related protein |
| b1003 | yccJ | RpoS | PF13993 family protein YccJ | Stress-related protein |
| b1051 | msyB | RpoS | acidic protein that suppresses heat sensitivity of a <i>secY</i> mutant | Stress-related protein |
| b1188 | ycgB | RpoS | PF04293 family protein YcgB | Stress-related protein |
| b1195 | ymgE | RpoS | PF04226 family protein YmgE | Stress-related protein |
| b1536 | ydeI | RpoS | BOF family protein YdeI | Stress-related protein |
| b1668 | ydhS | RpoS | FAD/NAD(P) binding domain-containing protein YdhS | Stress-related protein |
| b1784 | yeaH | RpoS | DUF444 domain-containing protein YeaH | Stress-related protein |
| b1836 | yebV | RpoS | protein YebV | Stress-related protein |
| b1955 | yedP | RpoS | putative mannosyl-3-phosphoglycerate phosphatase | Stress-related protein |
| b2080 | yegP | RpoS | DUF1508 domain-containing protein YegP | Stress-related protein |
| b2266 | elaB | RpoS | tail anchored inner membrane protein | Stress-related protein |
| b2672 | ygaM | RpoS | DUF883 domain-containing protein YgaM | Stress-related protein |
| b3100 | yqjK | RpoS | conserved protein YqjK | Stress-related protein |
| b3239 | yhcO | RpoS | putative barnase inhibitor | Stress-related protein |
| b3361 | fic | RpoS | putative adenosine monophosphate—protein transferase Fic | Stress-related protein |
| b3555 | yiaG | RpoS | putative DNA-binding transcriptional regulator YiaG | Stress-related regulator |
| b4149 | blc | RpoS | outer membrane lipoprotein Blc | Stress-related protein |
| b4568 | ytjA | RpoS | DUF1328 domain-containing protein YtjA | Stress-related protein |
| b0379 | yaiY | SoxS | inner membrane protein | Oxidative stress-related protein |
| b0389 | yaiA | SoxS | protein YaiA | Oxidative stress-related protein |
| b0448 | mdlA | SoxS | ABC transporter family protein MdlA | |
| b0449 | mdlB | SoxS | ABC transporter family protein MdlB | |
| b0850 | ybjC | SoxS | DUF1418 domain-containing protein YbjC | Oxidative stress-related protein |
| b1580 | rspB | SoxS | putative zinc-binding dehydrogenase RspB | Oxidative stress-related protein |
| b1581 | rspA | SoxS | mandelate racemase/muconate lactonizing enzyme family protein RspA | Oxidative stress-related protein |
| b2160 | yeiI | SoxS | putative sugar kinase YeiI | Oxidative stress-related protein |
| b2237 | inaA | SoxS | putative lipopolysaccharide kinase InaA | Oxidative stress-related protein |
| b2468 | aegA | SoxS | putative oxidoreductase Fe-S subunit AegA | Oxidative stress-related protein |
| b2898 | ygfZ | SoxS | folate-binding protein | Oxidative stress-related protein |
| b2962 | yggX | SoxS | putative Fe<sup>2+</sup>-trafficking protein | Oxidative stress-related protein |
| b3238 | yhcN | SoxS | DUF1471 domain-containing stress-induced protein YhcN | Oxidative stress-related protein |
| b3898 | frvX | SoxS | peptidase M42 family protein | Oxidative stress-related protein |
| b4379 | yjjW | SoxS | putative glycyl-radical enzyme activating enzyme YjjW | Oxidative stress-related protein |
| b4536 | yobH | SoxS | protein YobH | Oxidative stress-related protein |
| b4380 | yjjI | SoxS, Fnr/IscR | DUF3029 domain-containing protein YjjI | |
| b2998 | yghW | Tryptophan, ArcA, GcvA | DUF2623 domain-containing protein YghW | |
| b0270 | yagG | XylR | putative D-xylonate transporter YagG | |
| b0271 | yagH | XylR | CP4-6 prophage; putative xylosidase/arabinosidase | |
| b3562 | yiaA | XylR | conserved inner membrane protein YiaA | Putative xylose or xyloside transporter |
| b3657 | yicJ | XylR | putative xyloside transporter YicJ | |
| b2001 | yeeR | yeeRS-flu | CP4-44 prophage; inner membrane protein YeeR | |
| b2002 | yeeS | yeeRS-flu | CP4-44 prophage; RadC-like JAB domain-containing protein YeeS | |
| b0296 | ykgM | ZntR/Zur | putative ribosomal protein | |
| b1974 | yodB | ZntR/Zur | putative cytochrome | |

**Table S5: Genes in the BW25113 i-modulon**

| I-modulon Gene/Operon | I-modulon Gene Coefficient(s) | Genotypic Change | Phenotypic Change |
|---|---|---|---|
| *araBAD* | -0.16; -0.20; -0.10 | Deletion of *araBA*, Truncation of *araD* | No expression in BW25113 |
| *rhaBAD* | -0.07; -0.11; -0.07 | Deletion | No expression in BW25113 |
| *lacZ* | -0.06 | Truncation | Low expression in BW25113 |
| *hsdR* | -0.07 | Truncation | Low expression in BW25113 |
| *yjjB* | -0.11 | Truncation | Low expression in BW25113 |
| *insAB-1* | -0.26; -0.17 | Deletion | No expression in BW25113 |
| *mhpCDEF* | 0.25; 0.23; 0.16; 0.18 | Insertion of IS30 in *mhpC* | IS30 contains internal promoter[54] leading to increased expression in BW25113 |
| *crl* | 0.06 | Pseudogene in MG1655, repaired in BW25113 | Increased expression in BW25113 |
| *glpR* | 0.18 | Pseudogene in MG1655, repaired in BW25113 | Increased expression in BW25113 |
| *tabA-ygjL* | 0.25; 0.18 | SNP at transcription start site (A→G. 4,474,834) | Increased expression in BW25113 |
| *fabAB* | 0.06; 0.10 | SNP in FabR | Decreased repression of *fabAB* by FabR |
| *pheA* | 0.09 | SNP in *pheV* tRNA | Altered tRNA-mediated attenuation via phe-tRNA |
