## Supplemental Data 2 for "The *Escherichia coli* Transcriptome Mostly Consists of Independently Regulated Modules"

### ArcA I-Modulon

Regulated by: ArcA

Biological Function: Electron transport chain

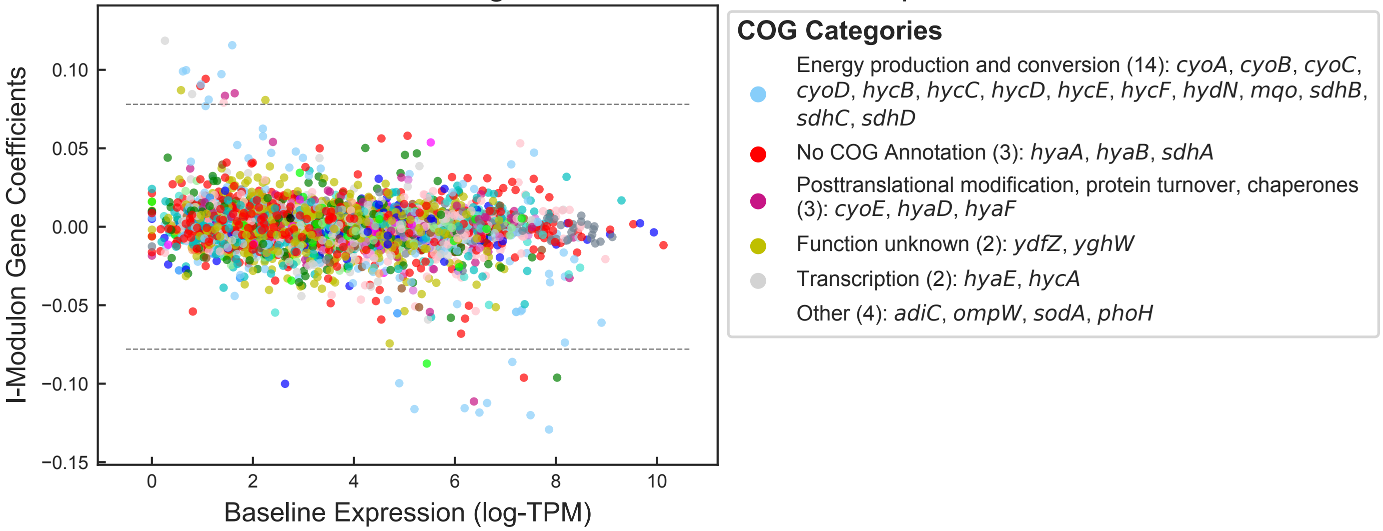


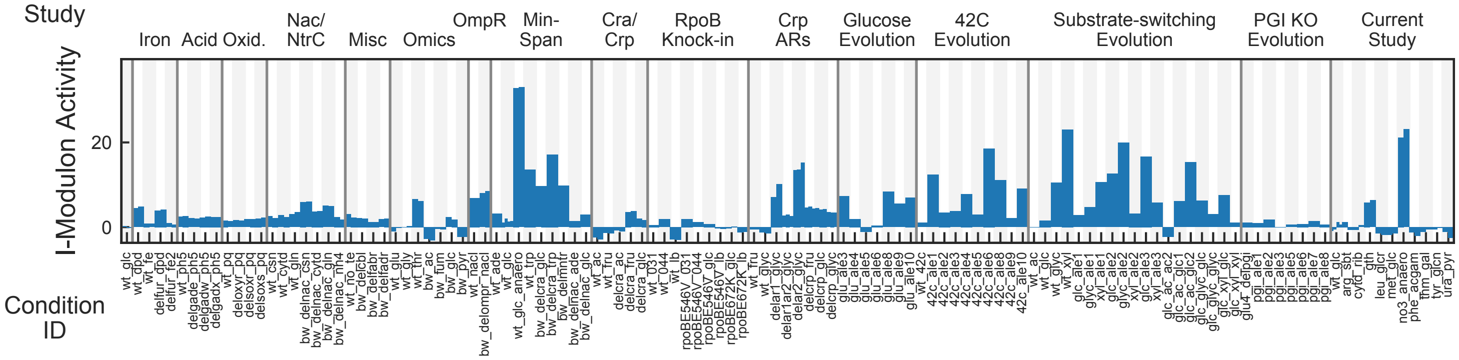


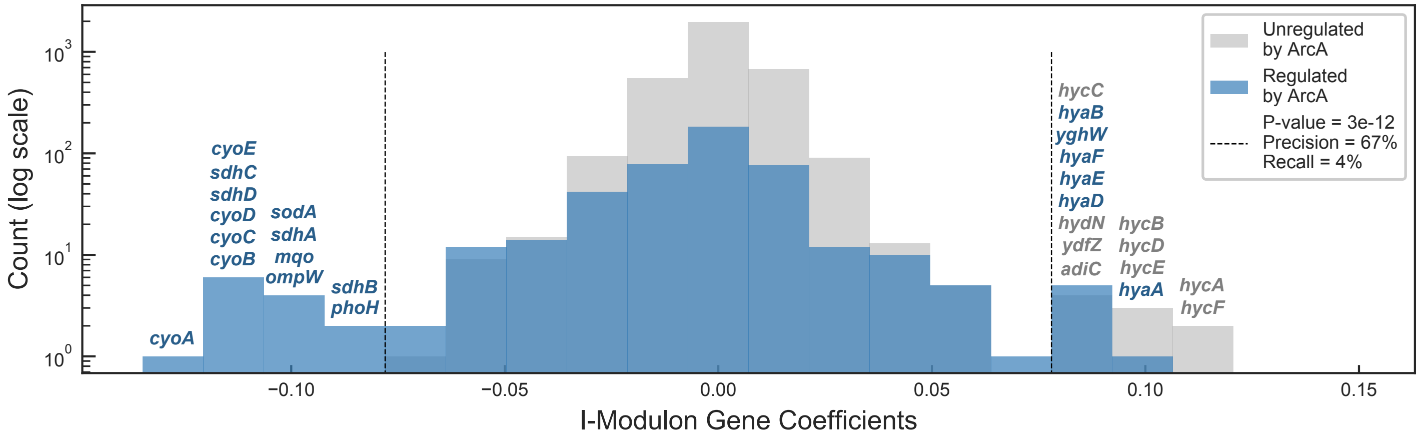


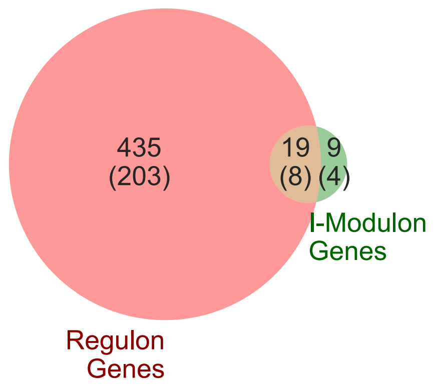


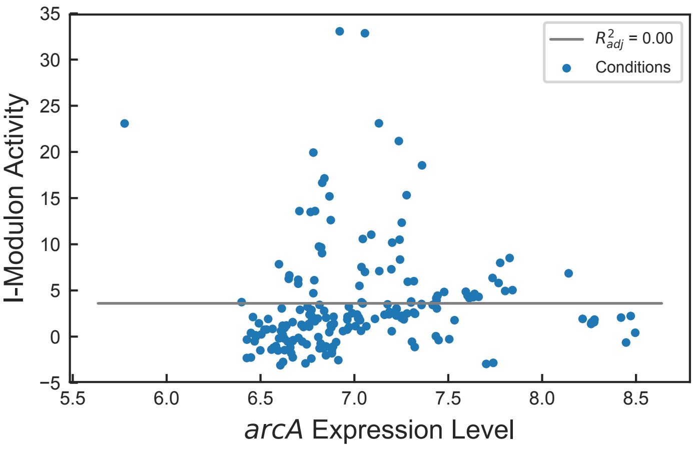

### BW25113 I-Modulon

Biological Function: Transcriptional difference between BW25113 and MG1655

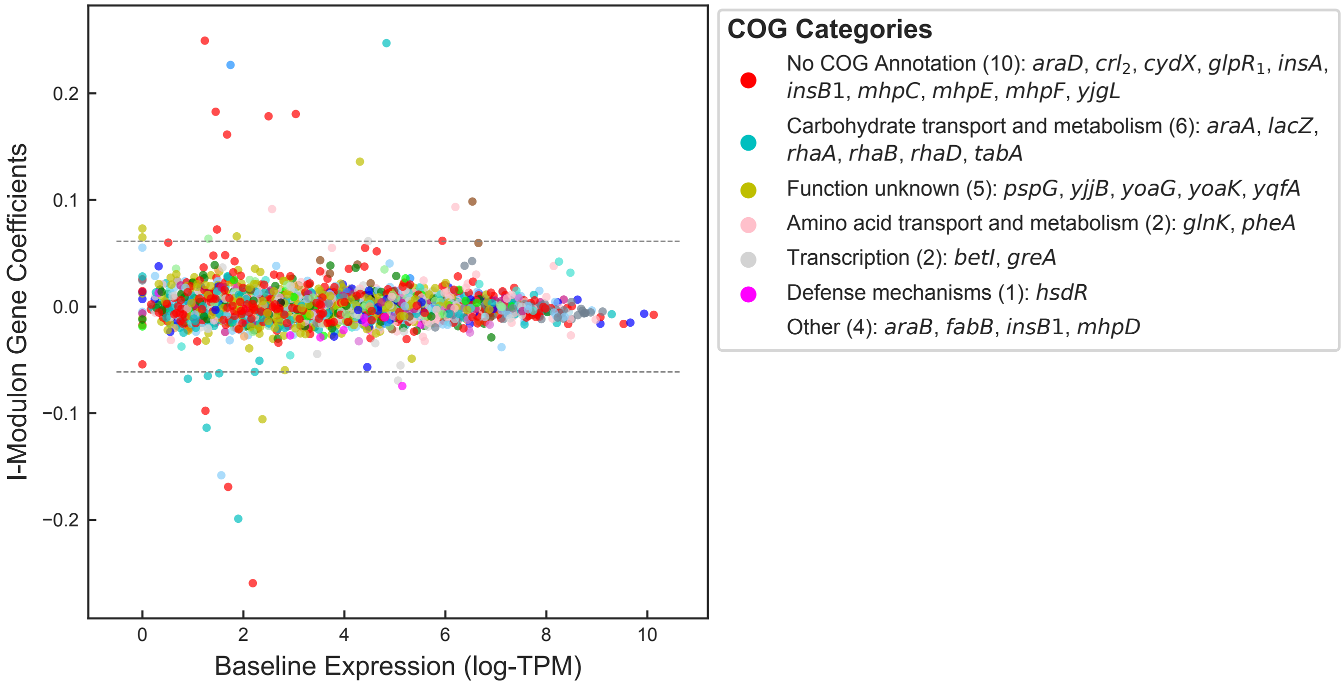


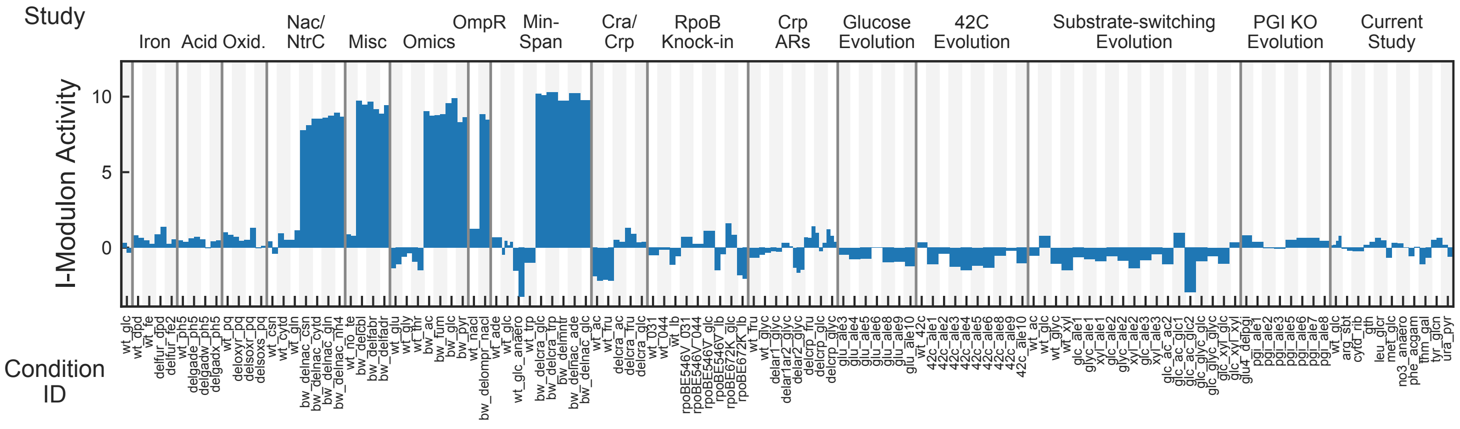


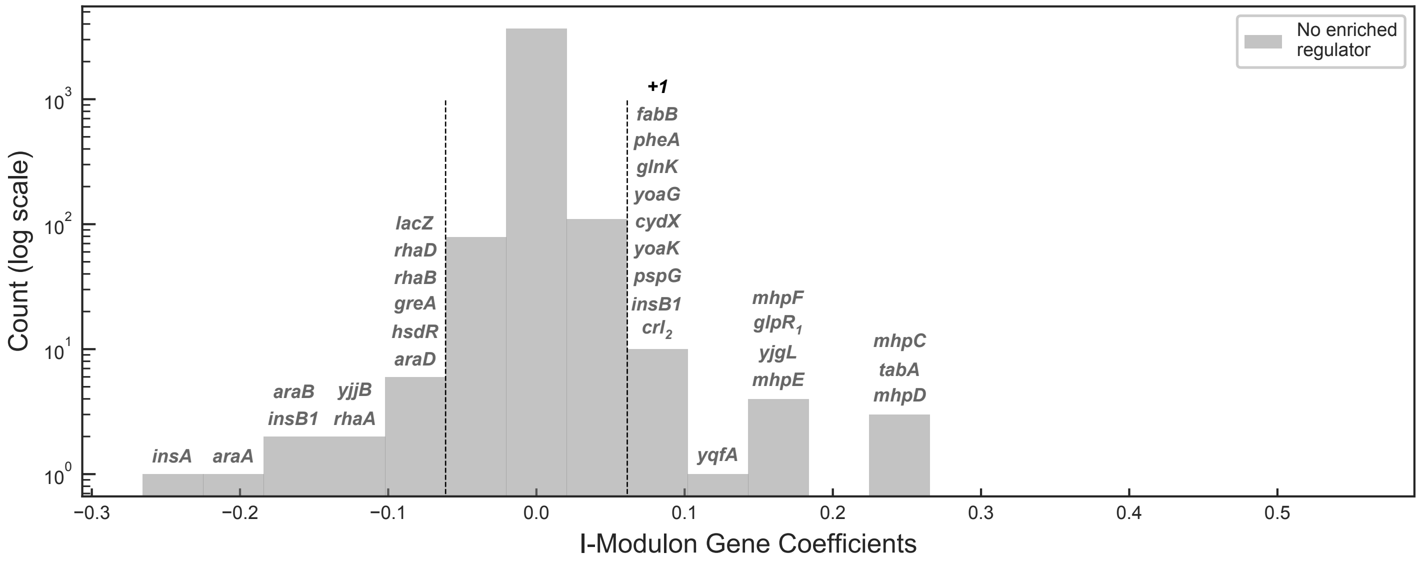

### Cbl + CysB I-Modulon

Regulated by: Cbl and CysB

Biological Function: Aliphatic sulfonate utilization

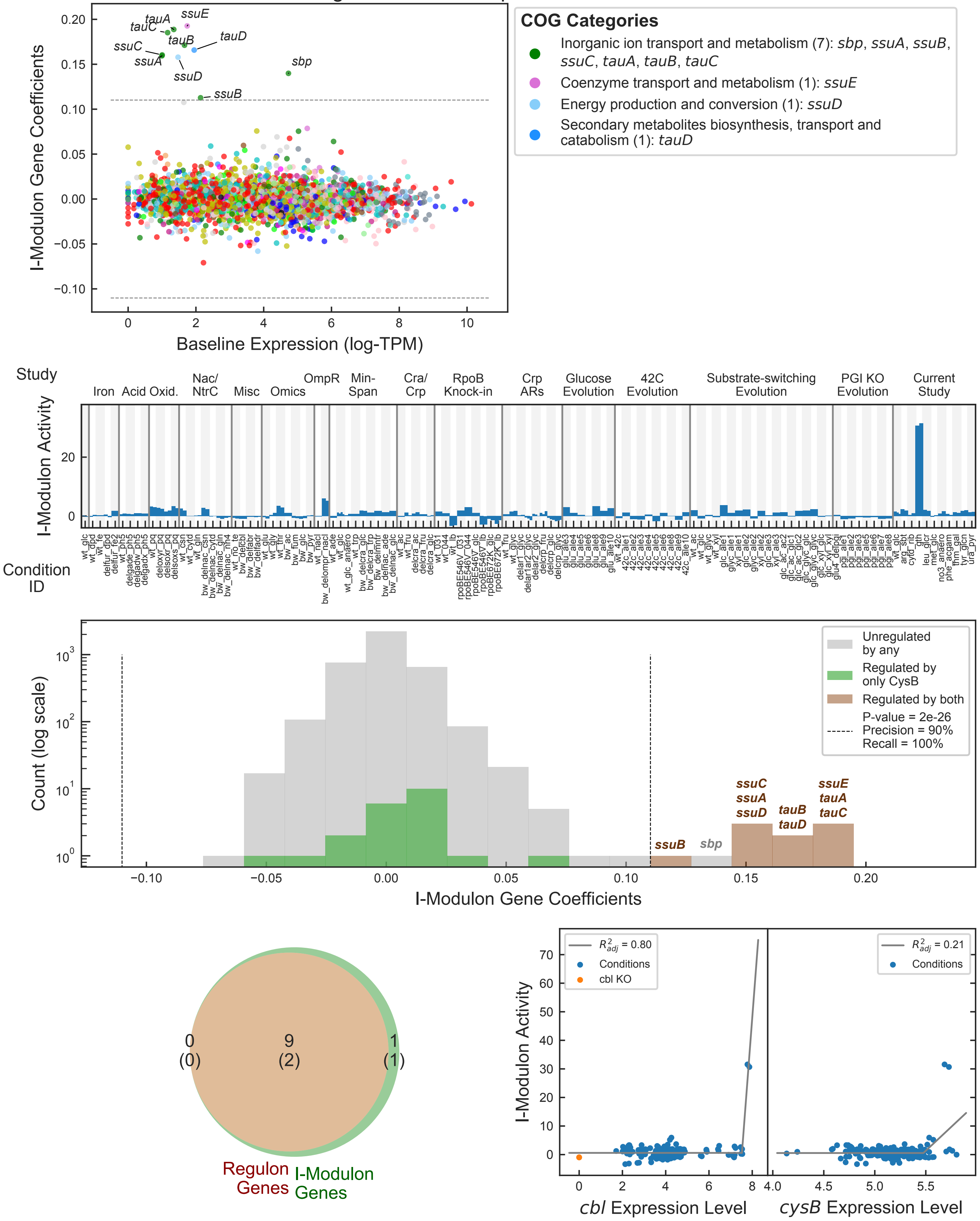

### CdaR I-Modulon

Regulated by: CdaR

Biological Function: Glucarate catabolism

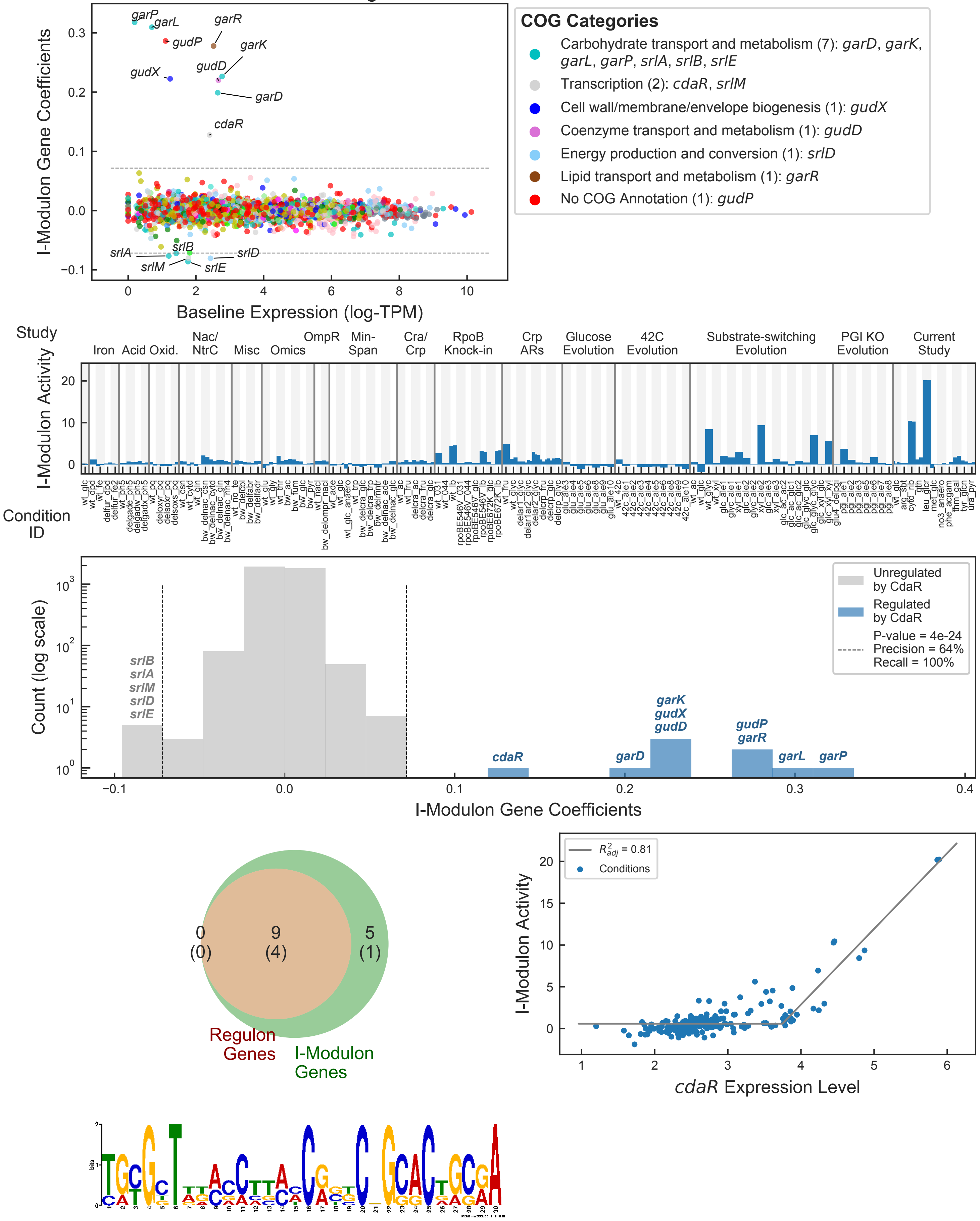


Motif E-value: 1.30e-04

Operons with Upstream Motif: 100%

### Cra I-Modulon

Regulated by: Cra

Biological Function: Central carbon metabolism

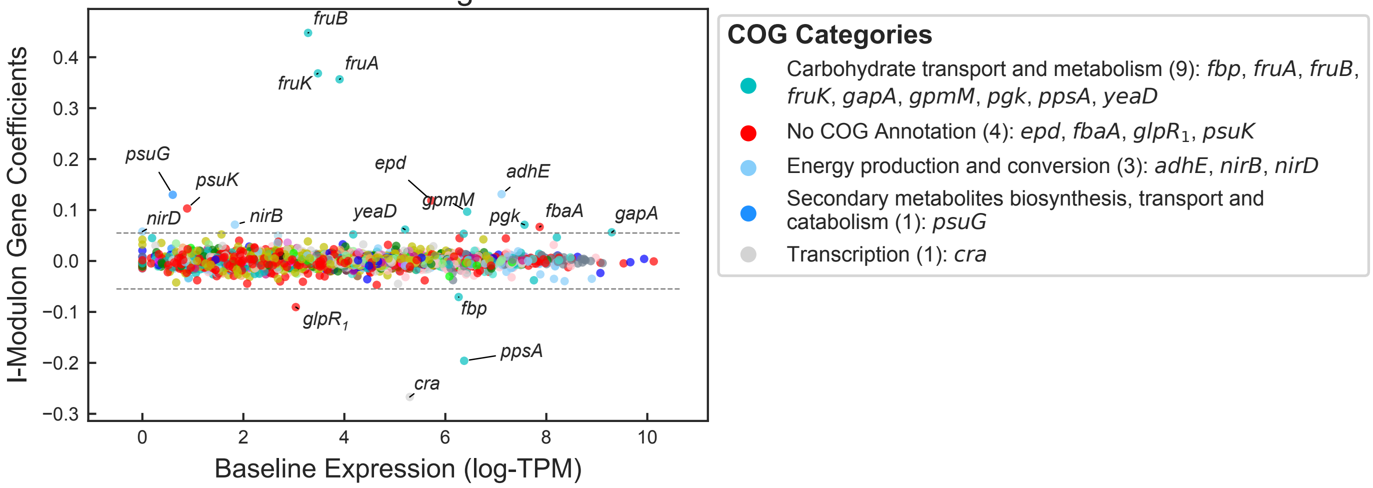


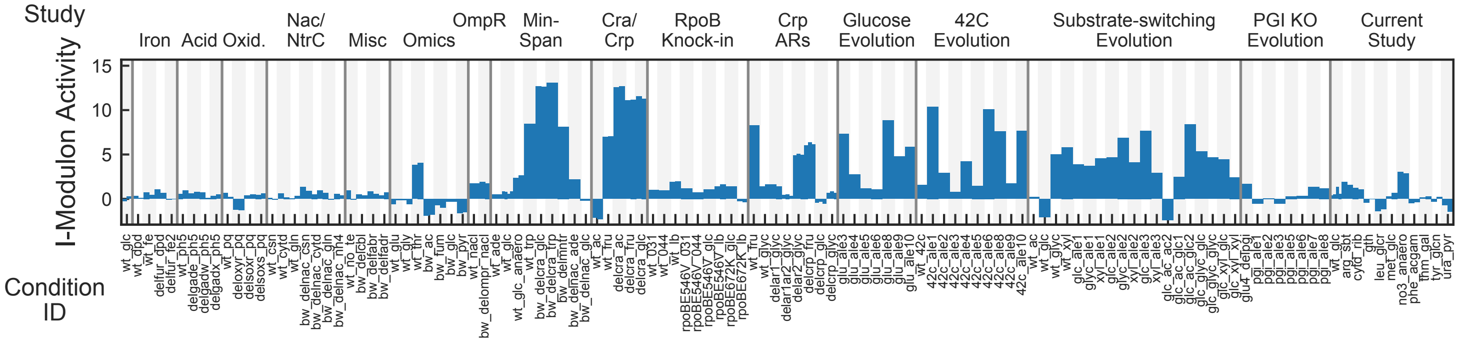


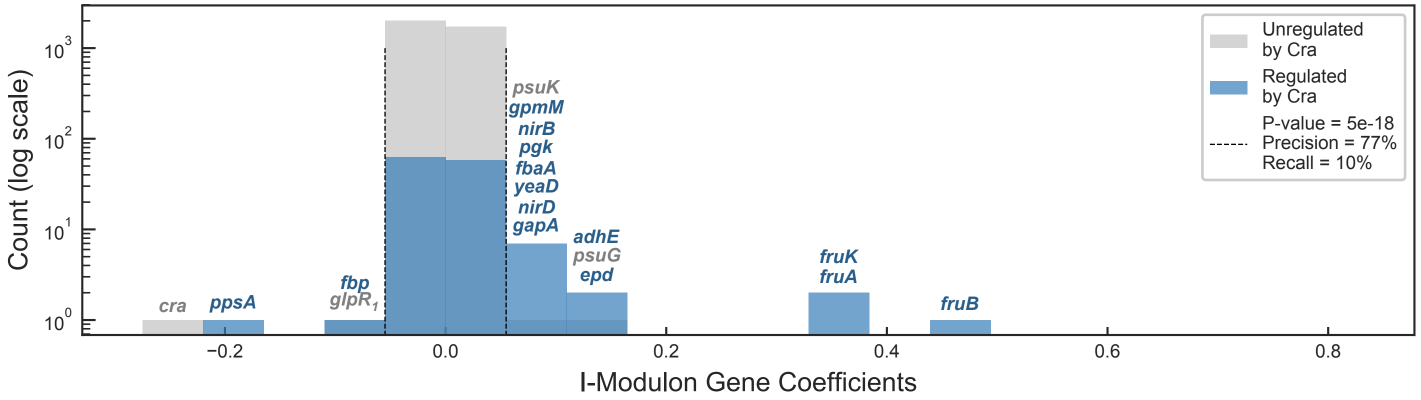


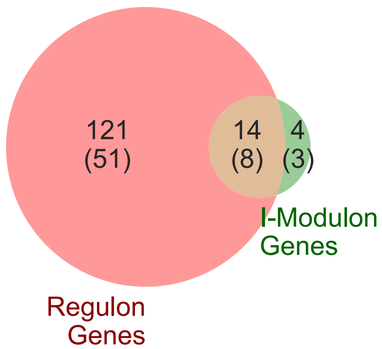


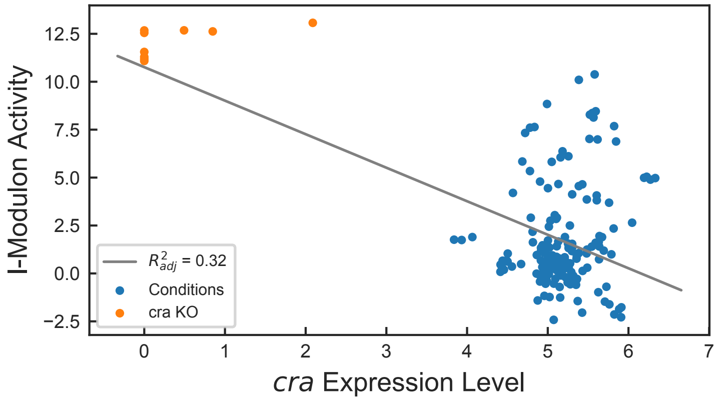


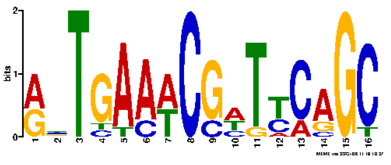

Motif E-value: 8.20e-05

Operons with Upstream Motif: 81%

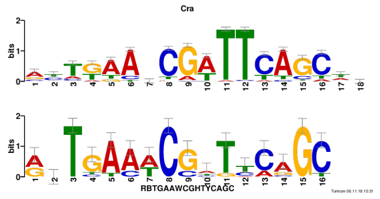


Motif similar to Cra (E-value: 1.67e-07)

### Crp – 1 I-Modulon

Regulated by: Crp

Biological Function: Carbon source catabolism

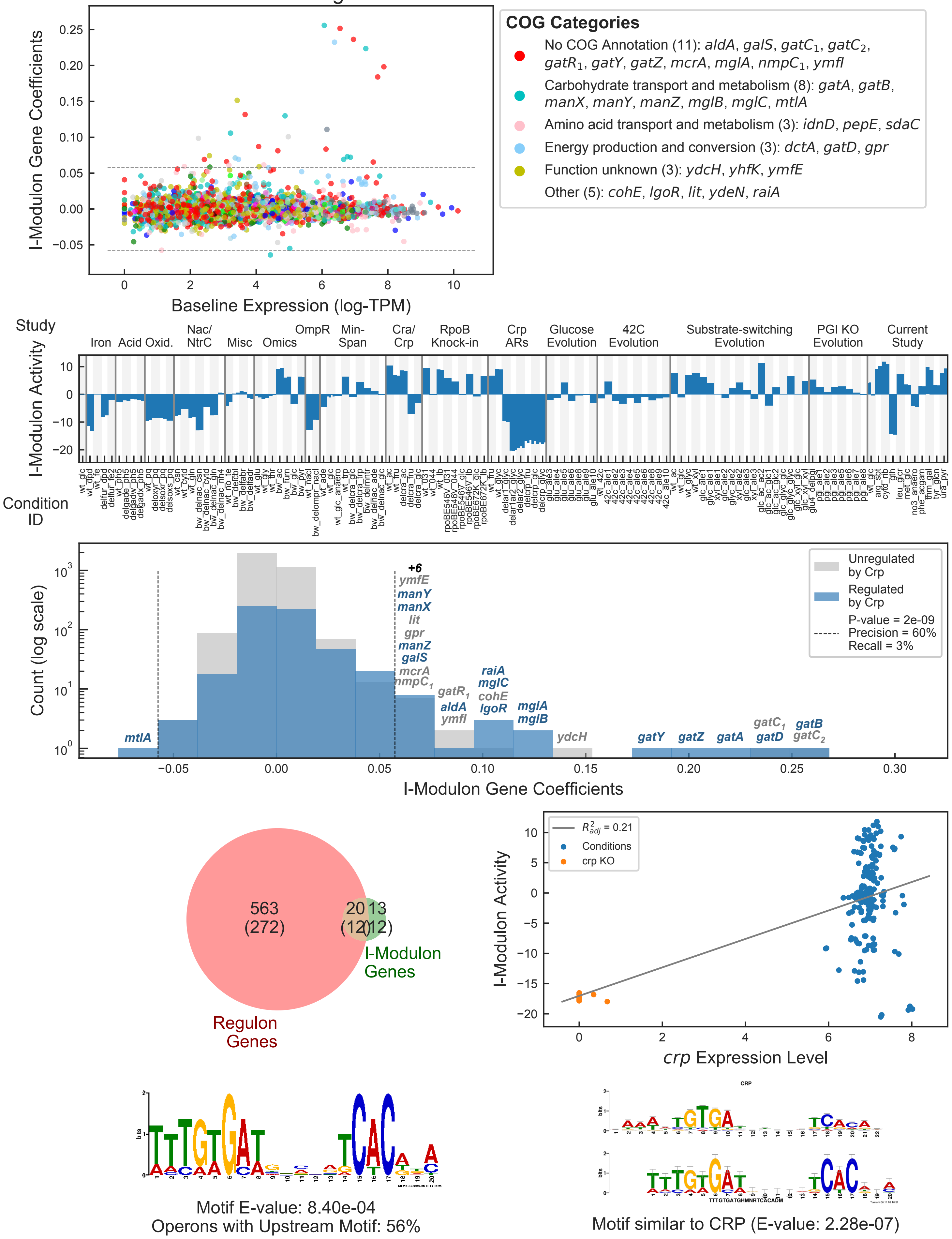

### Crp−2 I-Modulon

Regulated by: Crp

Biological Function: Carbon source catabolism

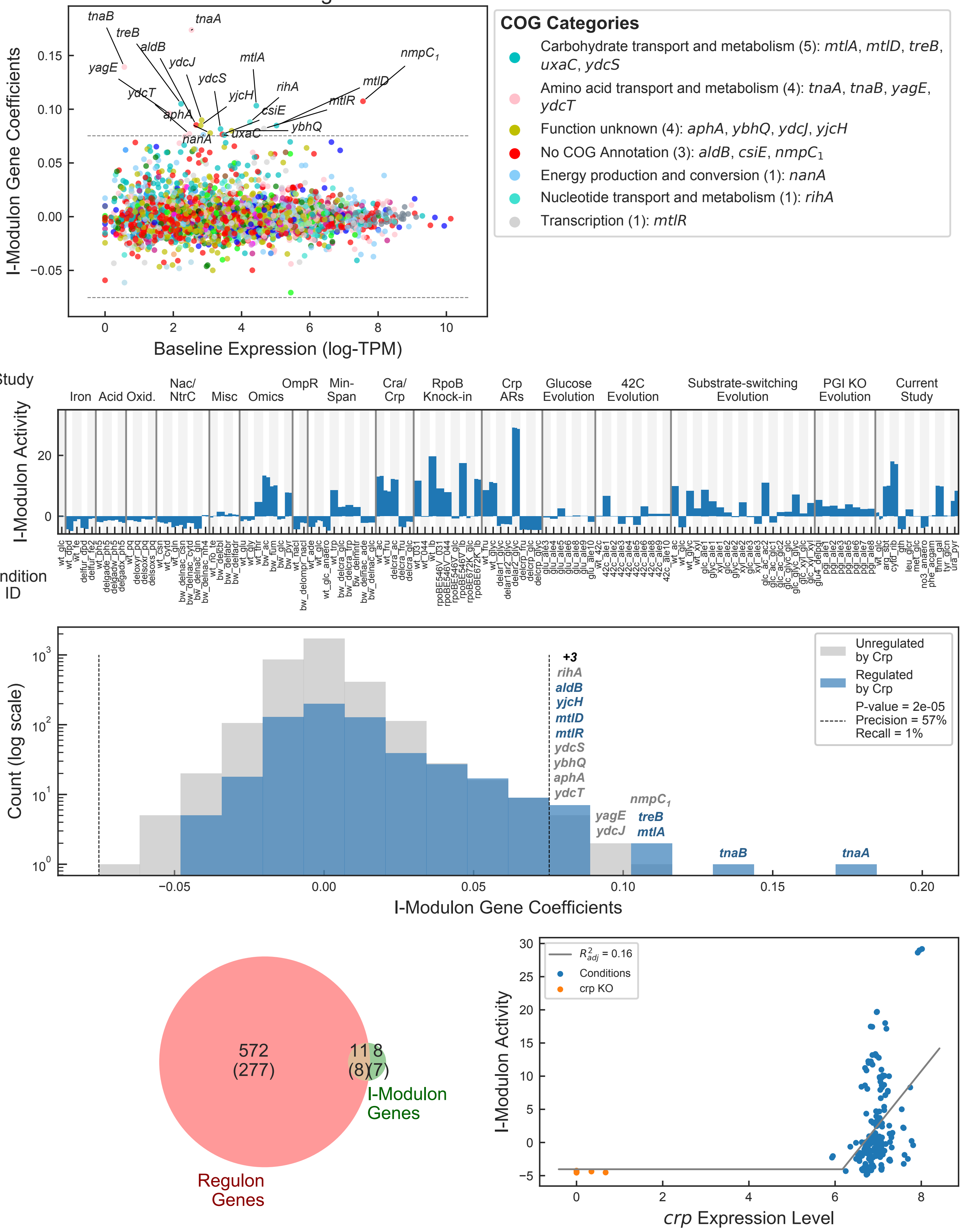

### CysB I-Modulon

Regulated by: CysB

Biological Function: Inorganic sulfate assimilation

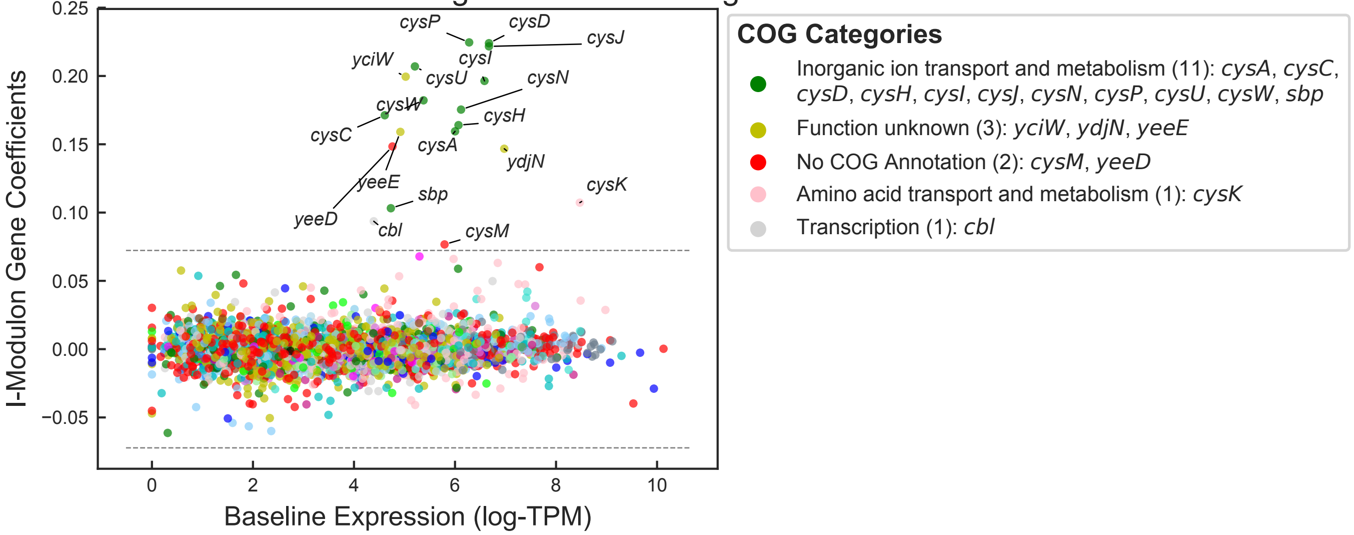


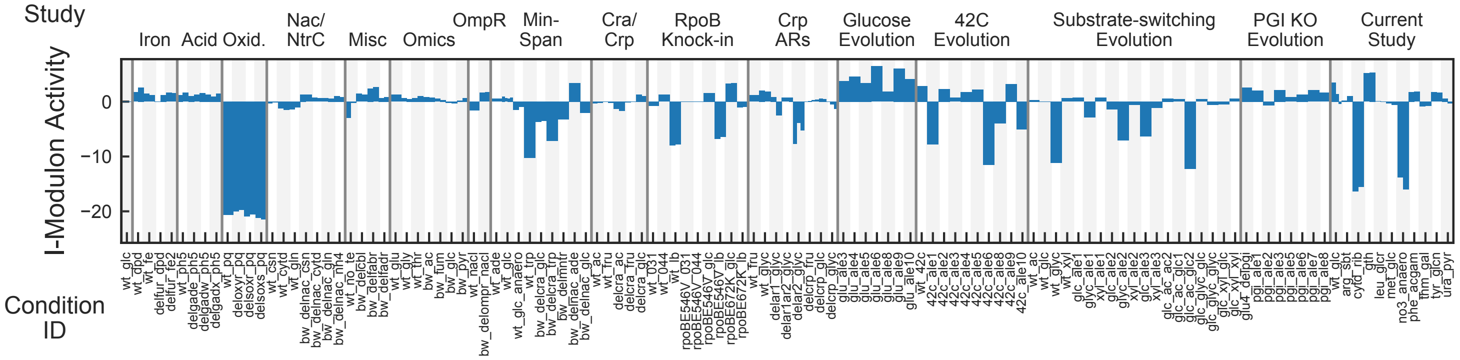


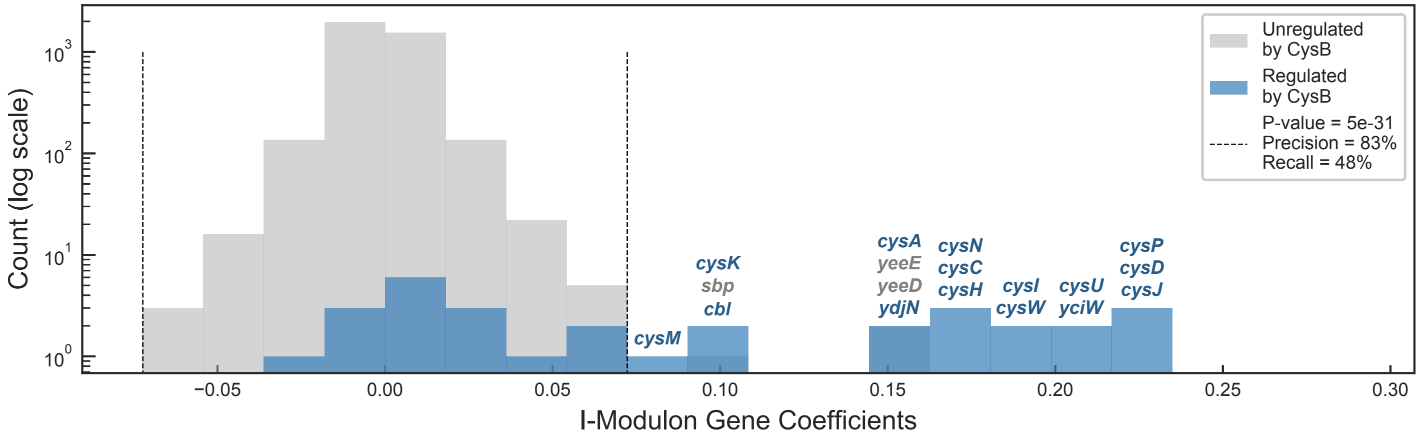


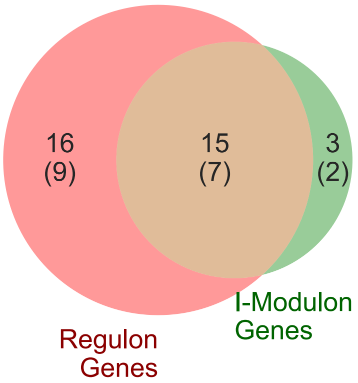


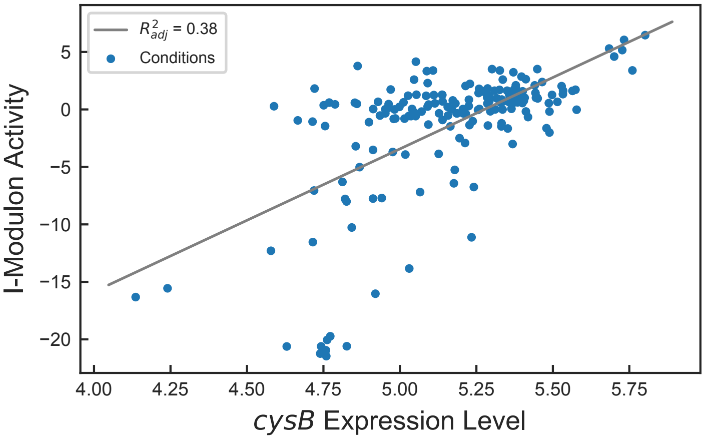

### EvgA I-Modulon

Regulated by: EvgA

Biological Function: Acid and osmotic stress response

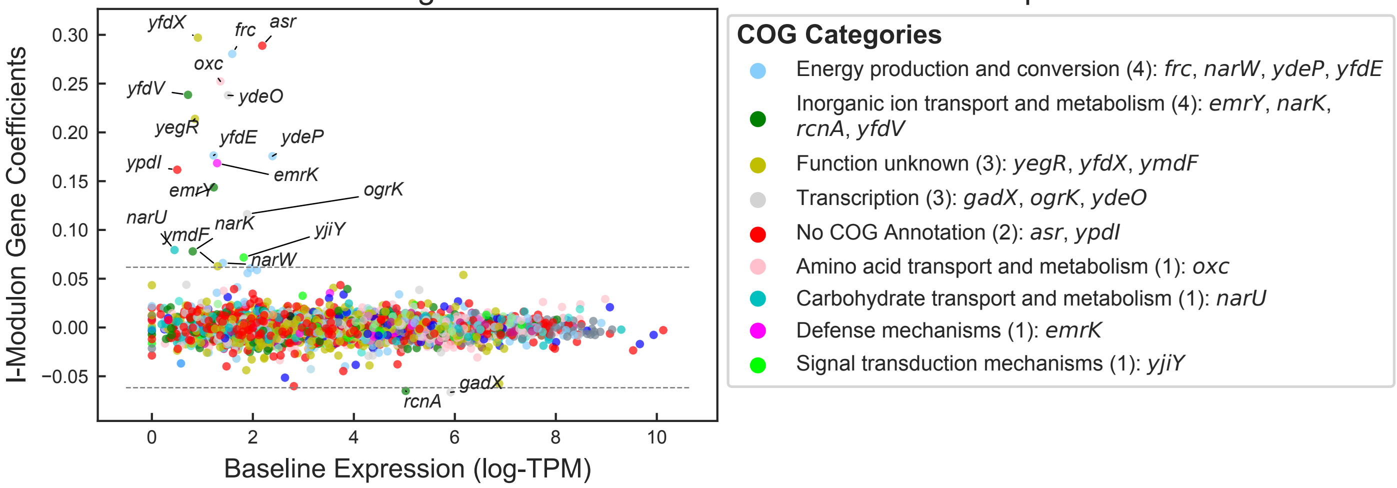


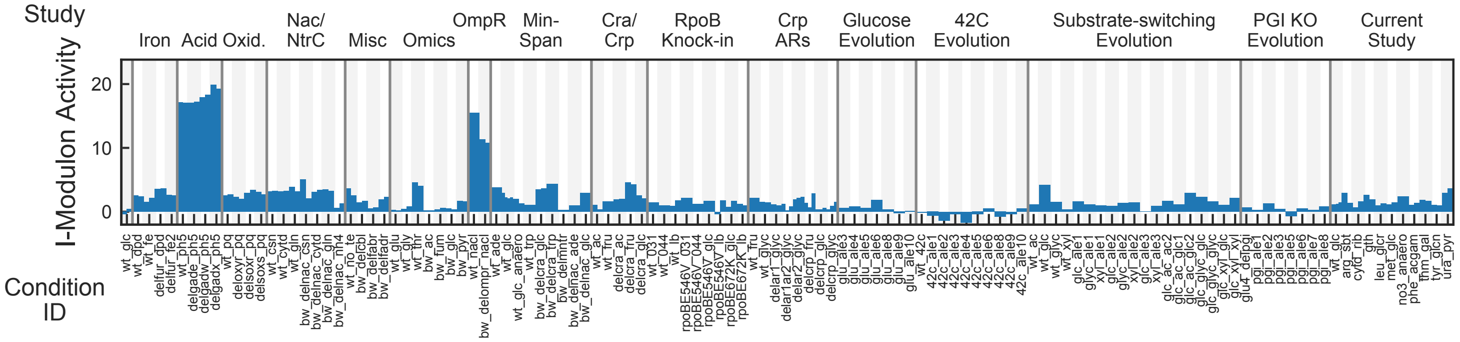


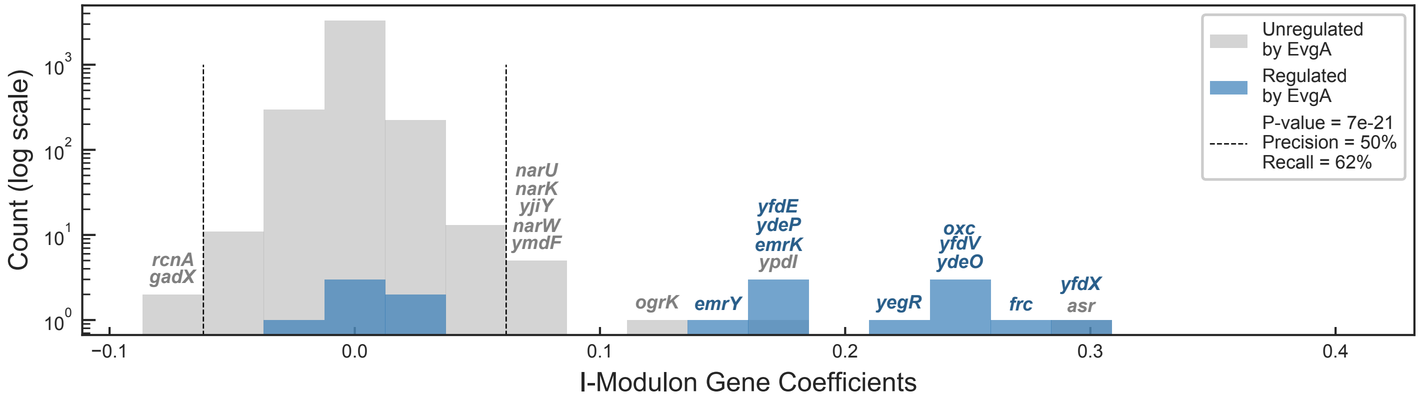


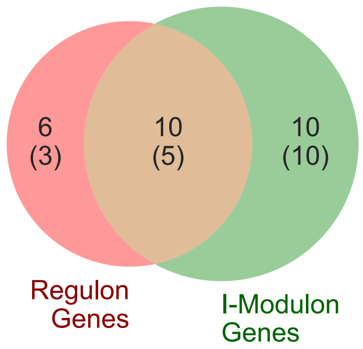


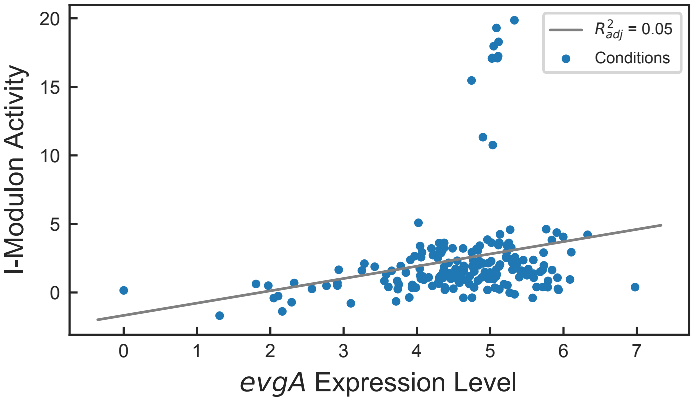


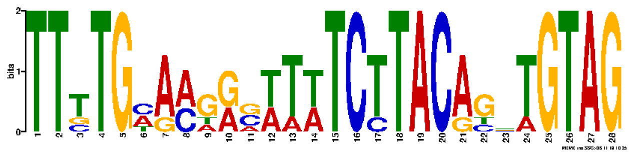

Motif E-value: 3.00e-04

Operons with Upstream Motif: 33%

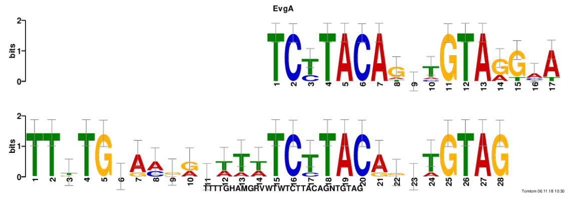


Motif similar to EvgA (E-value: 2.10e-11)

### FlhDC I-Modulon

Regulated by: FlhDC
Biological Function: Flagella assembly

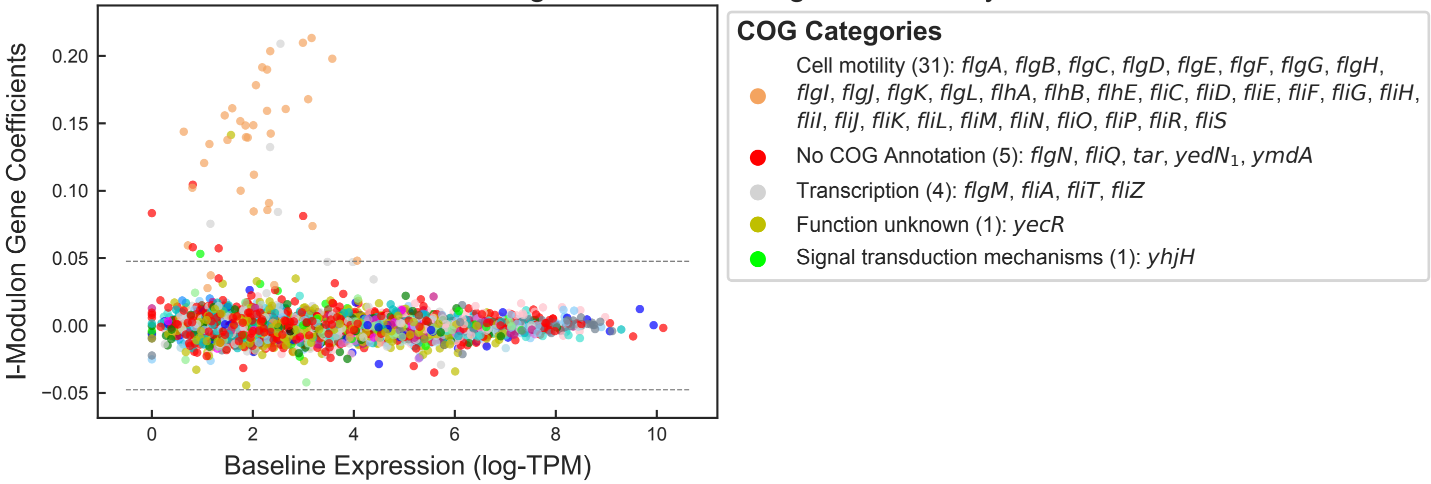


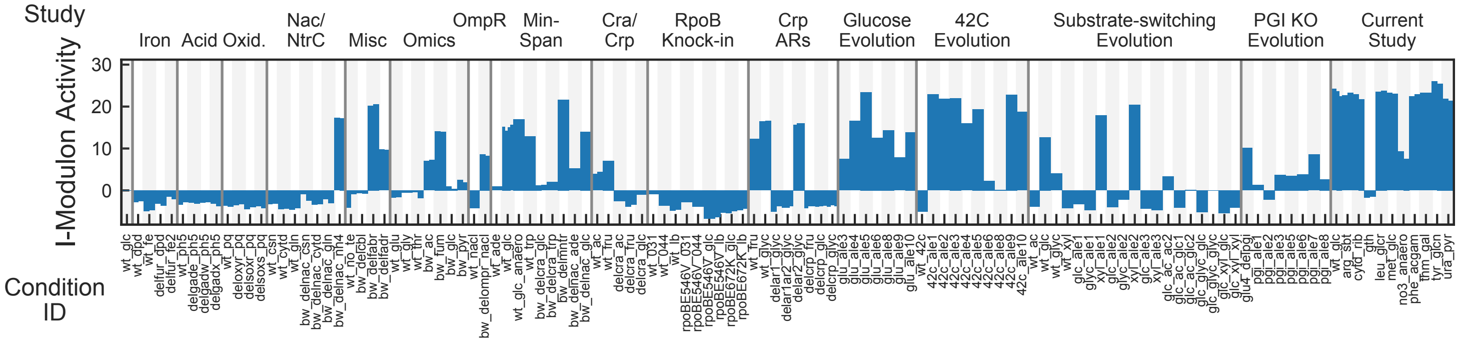


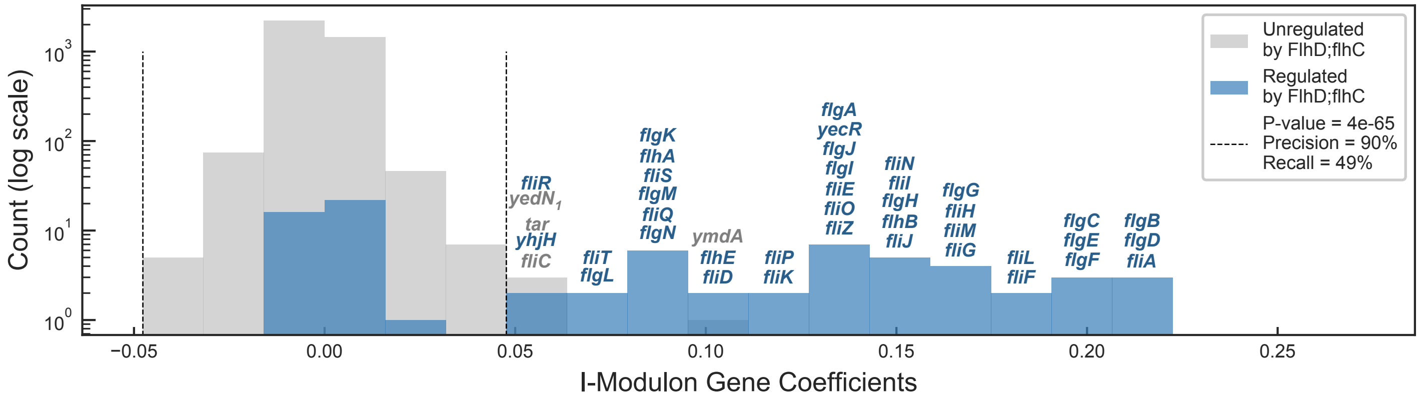


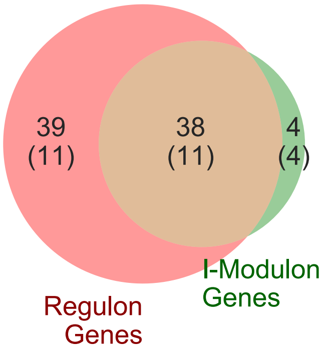


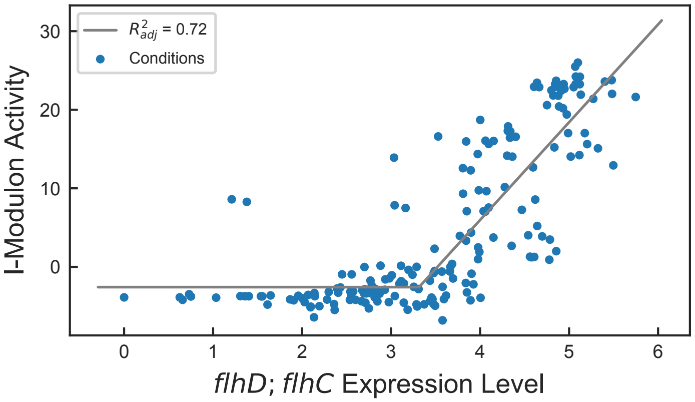


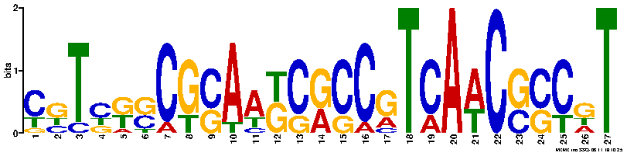

Motif E-value: 7.00e-11
Operons with Upstream Motif: 53%

### FliA I-Modulon

Regulated by: FliA

Biological Function: Chemotaxis

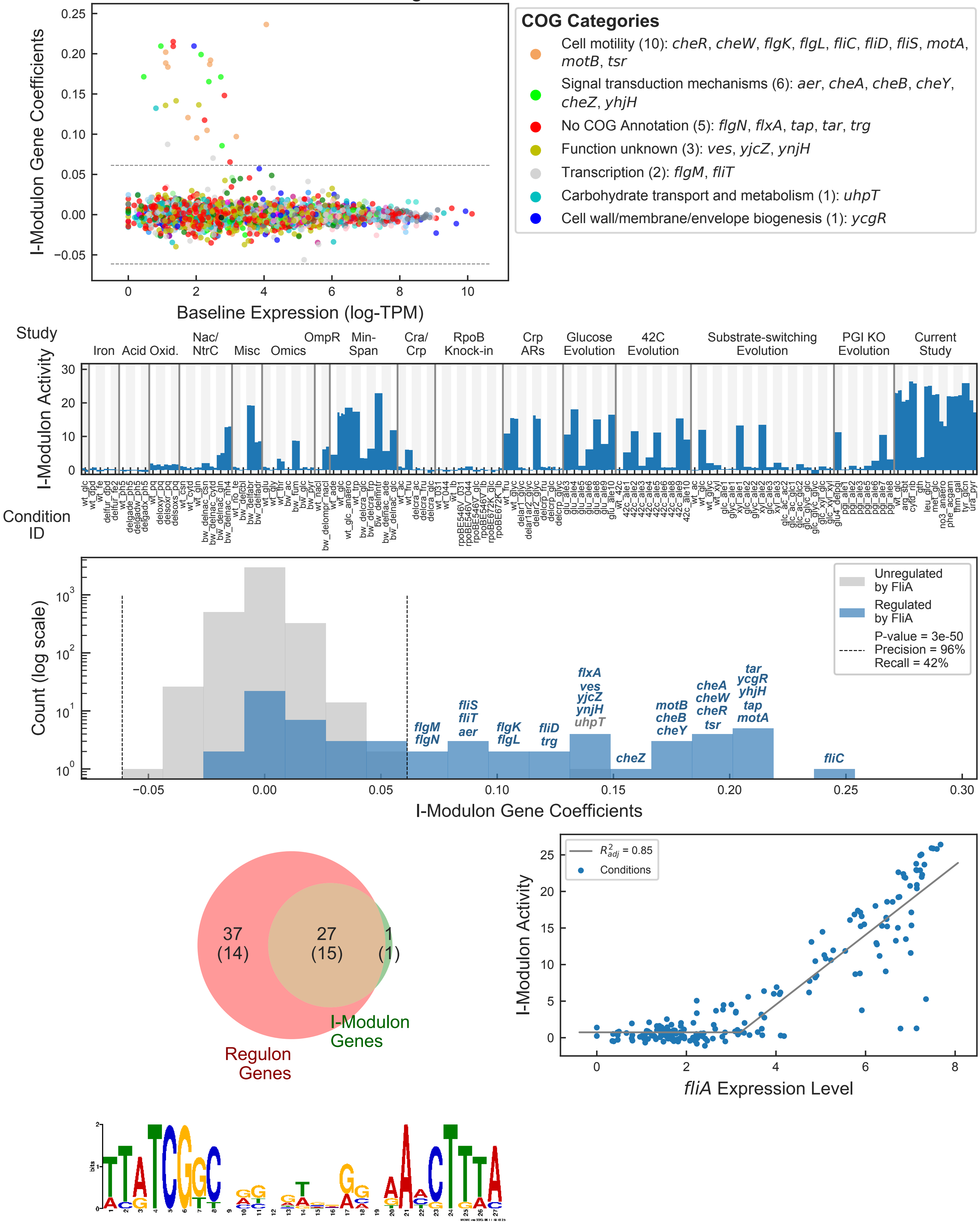


Motif E-value: 3.20e-09

Operons with Upstream Motif: 56%

### Fur – 1 I-Modulon

Regulated by: Fur

Biological Function: Iron homeostasis

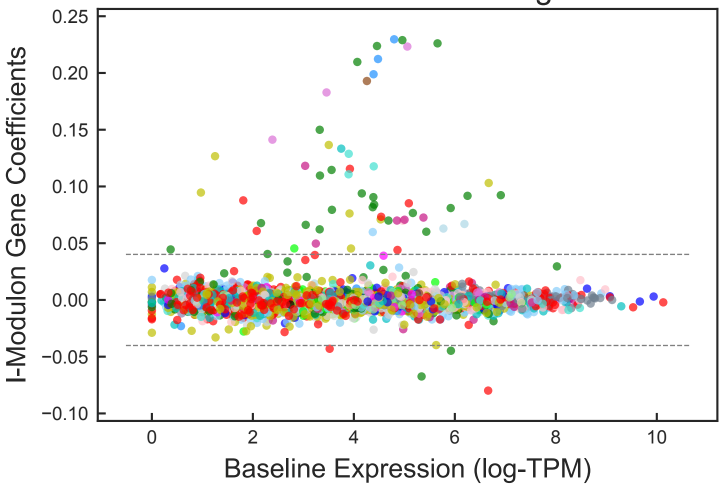


**COG Categories**

- Inorganic ion transport and metabolism (25): *bfr*, *cirA*, *efeB*, *efeO*, *fecA*, *fecC*, *fecR*, *fepA*, *fepB*, *fepD*, *fepE*, *fepG*, *fes*, *fhuA*, *fhuB*, *fhuC*, *fhuD*, *fhuE*, *fhuF*, *fiu*, *ftnA*, *tonB*, *yddA*, *yddB*, *yqjH*
- No COG Annotation (8): *efeU*$_2$, *fecI*, *fepC*, *gadW*, *sodB*, *ydiE*, *yoeA*$_1$, *yoeG*$_1$
- Function unknown (7): *sufA*, *sufE*, *ybaN*, *ybdZ*, *ybiX*, *yjjZ*, *yncE*
- Posttranslational modification, protein turnover, chaperones (5): *nrdH*, *pqqL*, *sufB*, *sufC*, *sufD*
- Other (16): *entC*, *entD*, *entH*, *nrdE*, *nrdF*, *nrdI*, *entB*, *entE*, *entF*, *exbB*, *exbD*, *sufS*, *entS*, *bfd*, *entA*, *ybiI*

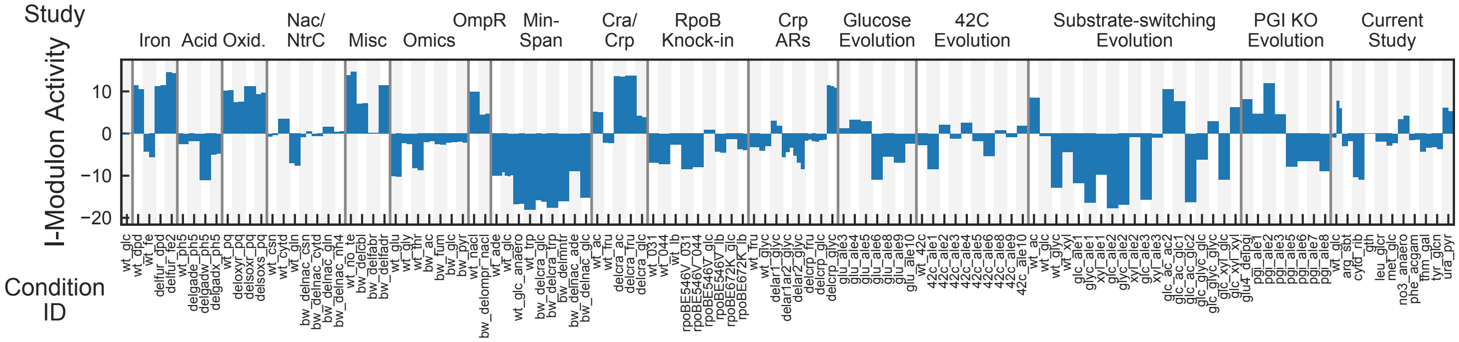


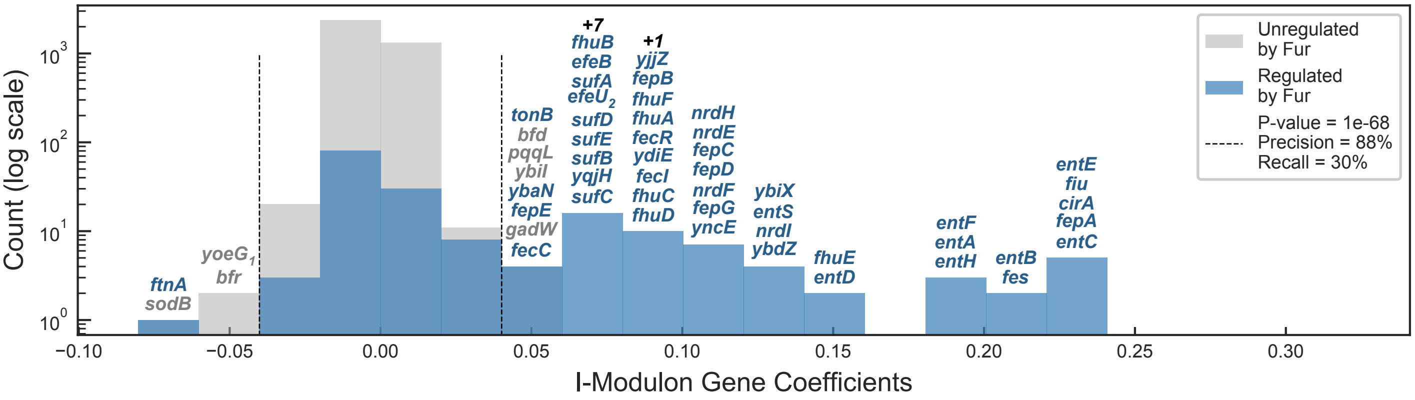


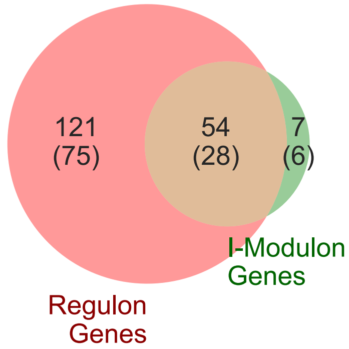


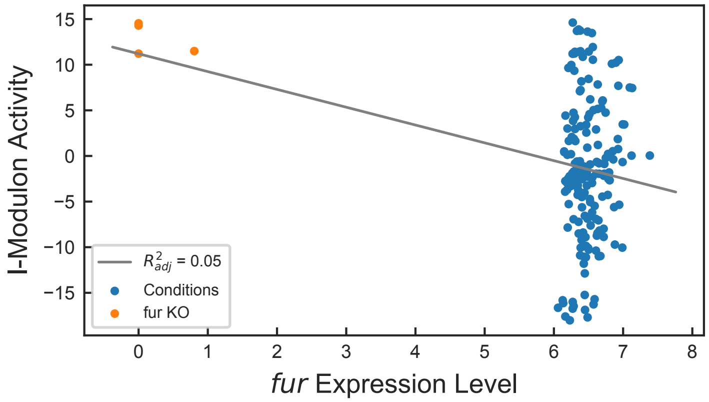


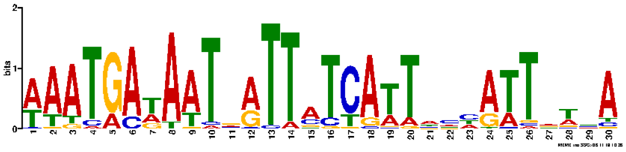

Motif E-value: 6.70e-55

Operons with Upstream Motif: 73%

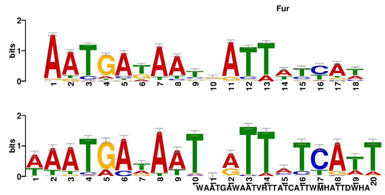


Motif similar to Fur (E-value: 2.36e-10)

### Fur – 2 I-Modulon

Regulated by: Fur

Biological Function: Iron homeostasis

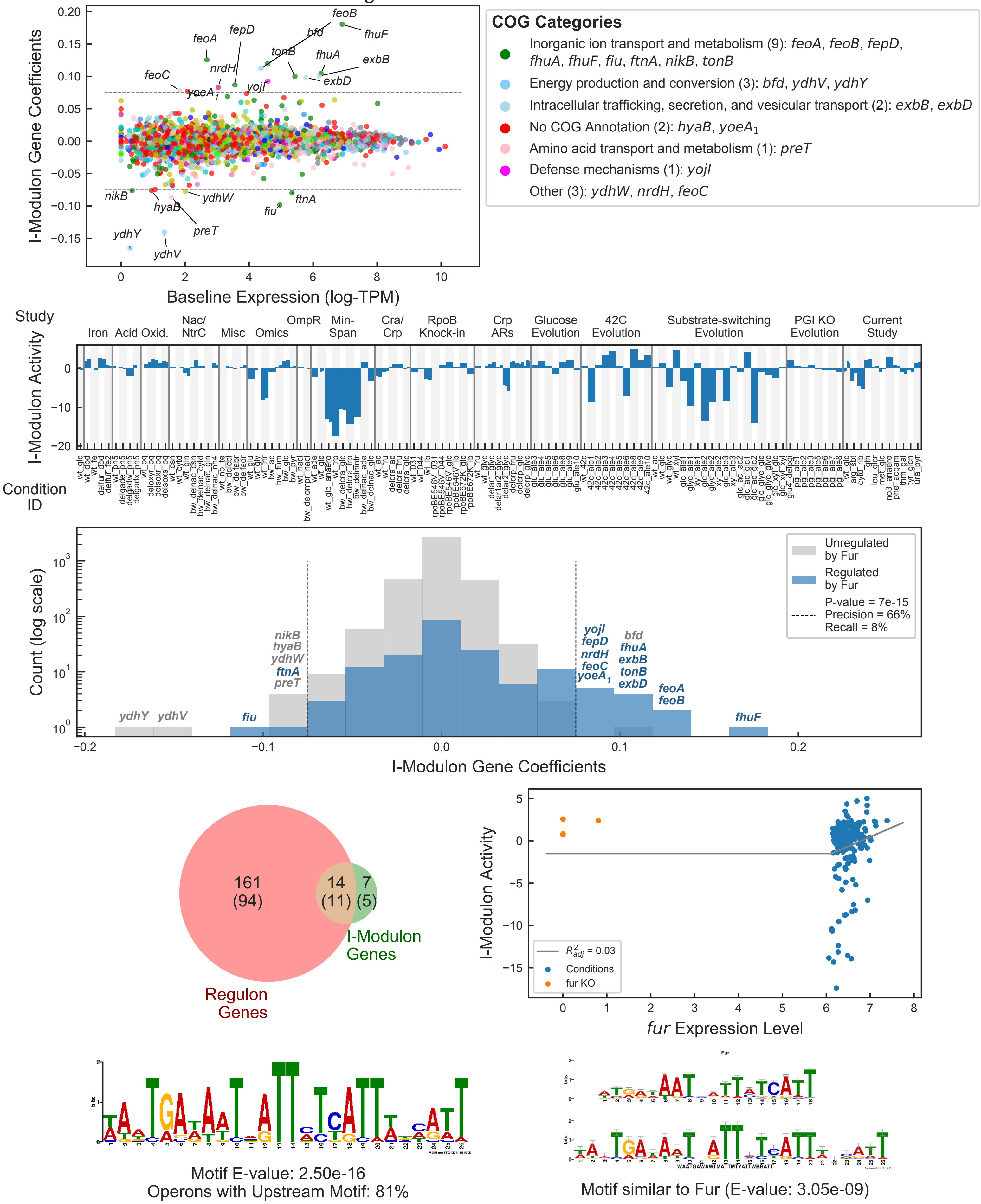

### GlpR I-Modulon

Regulated by: GlpR

Biological Function: Glycerol catabolism

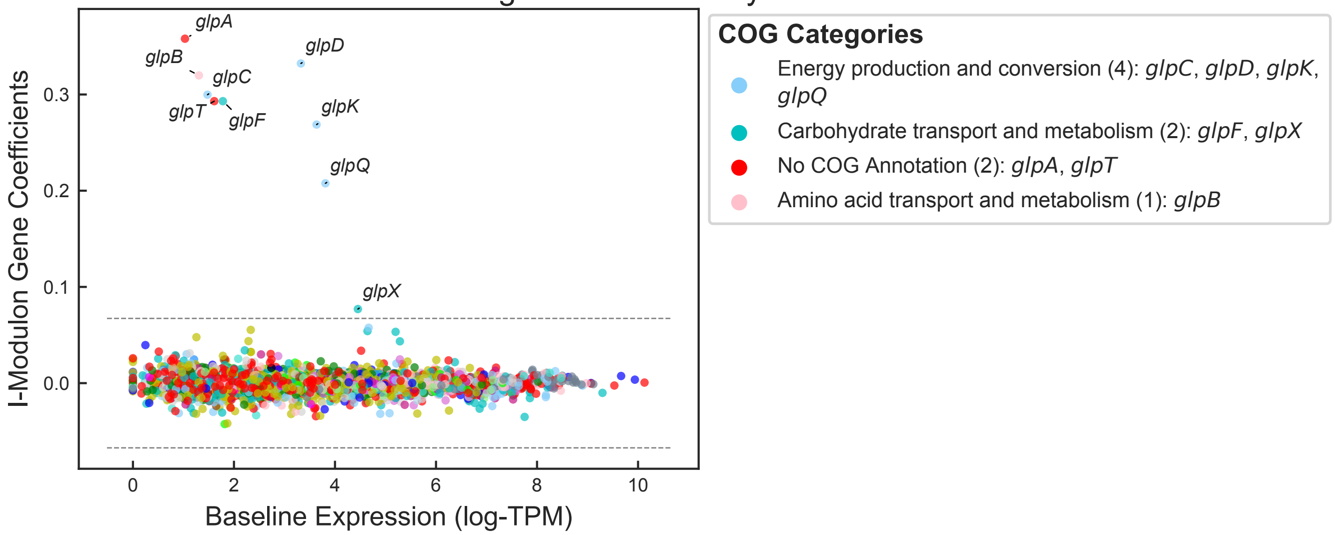


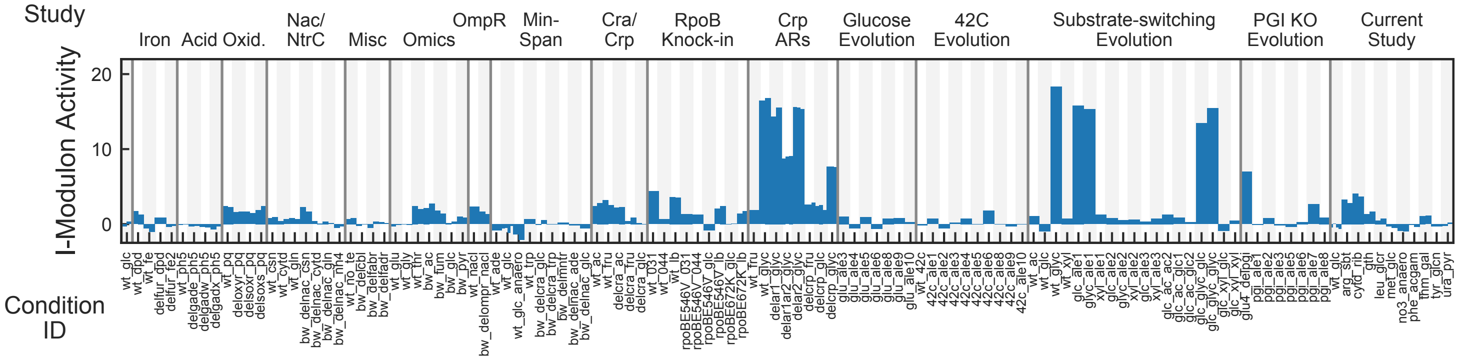


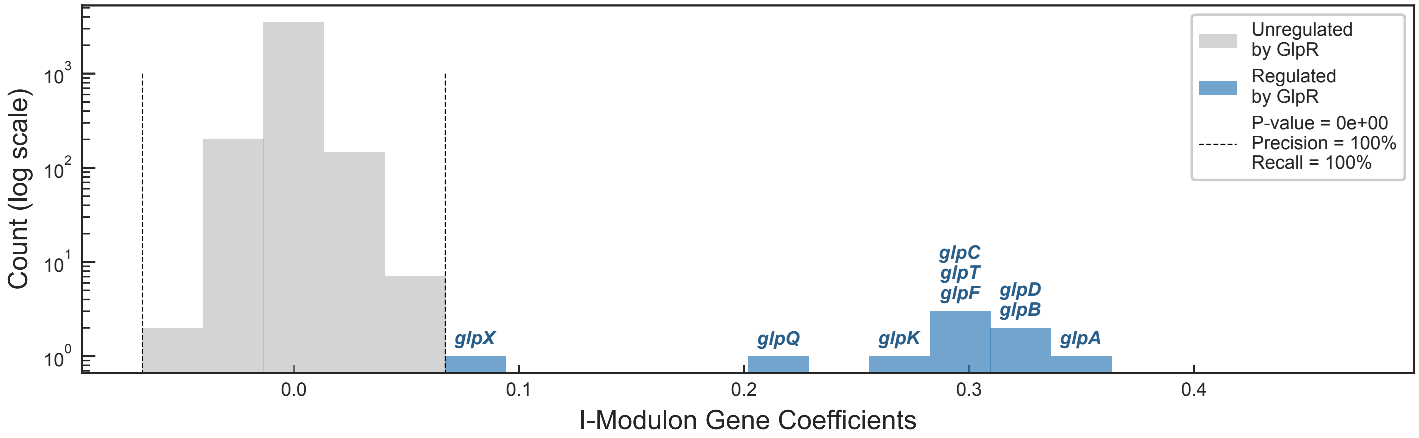


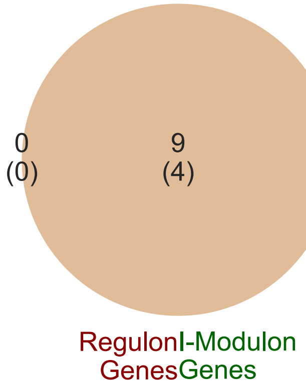

### GntR/TyrR I-Modulon

Regulated by: GntR or TyrR

Biological Function: Gluconate catabolism and tyrosine biosynthesis

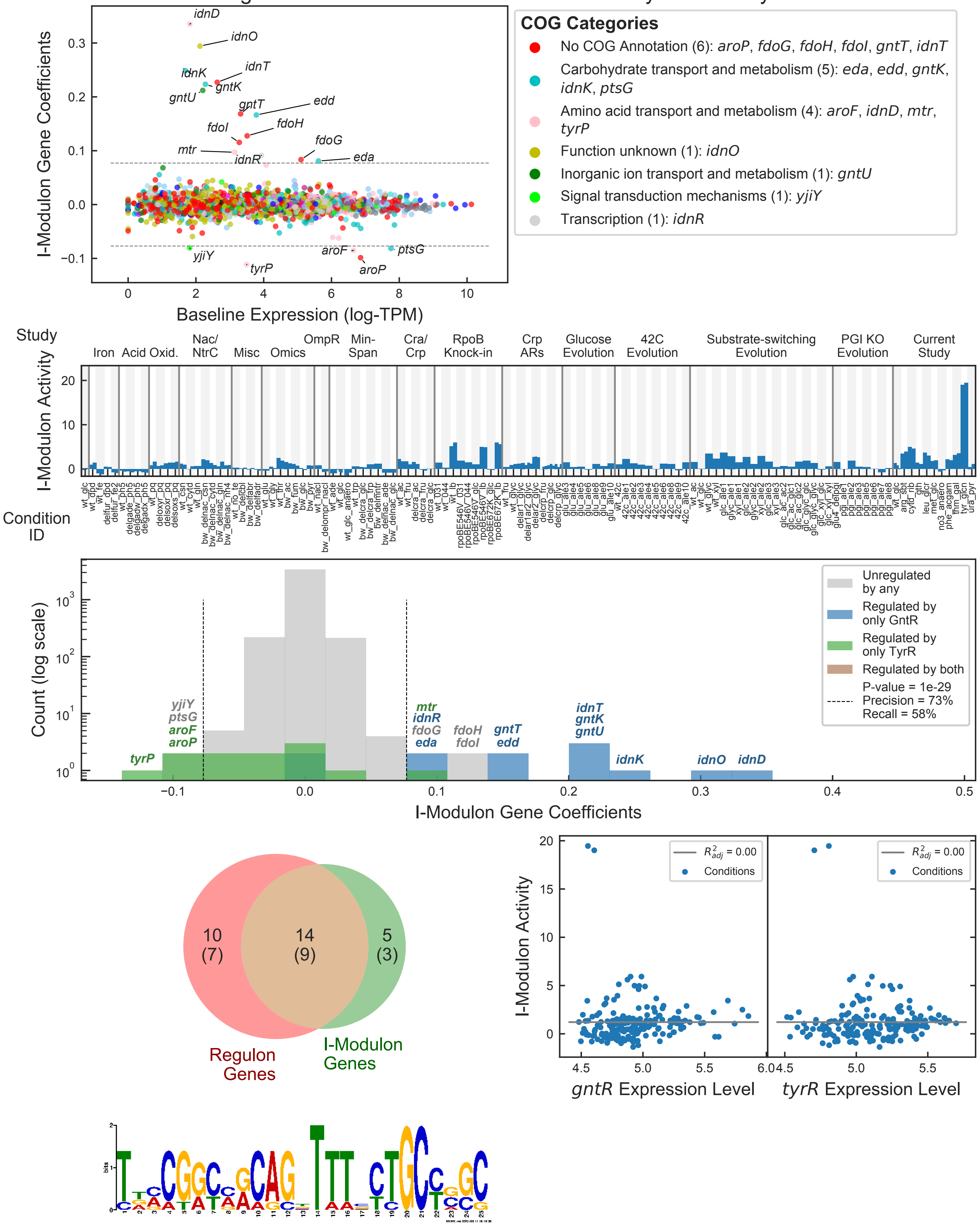


Motif E-value: 2.60e-04

Operons with Upstream Motif: 58%

### GutMR I-Modulon

Regulated by: GutM and GutR

Biological Function: Sorbitol catabolism

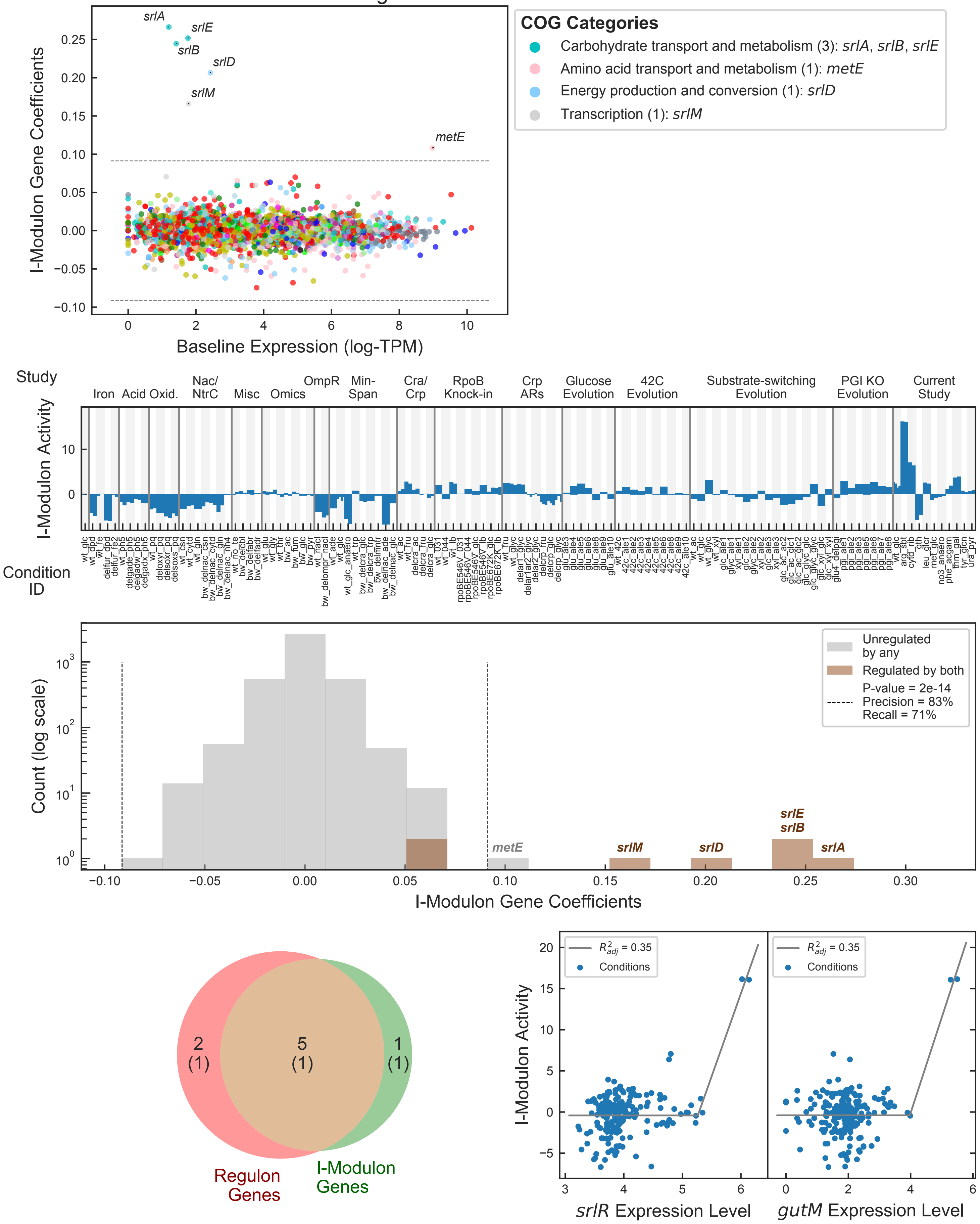

### His − tRNA I-Modulon

Regulated by: His-tRNA

Biological Function: Histidine biosynthesis

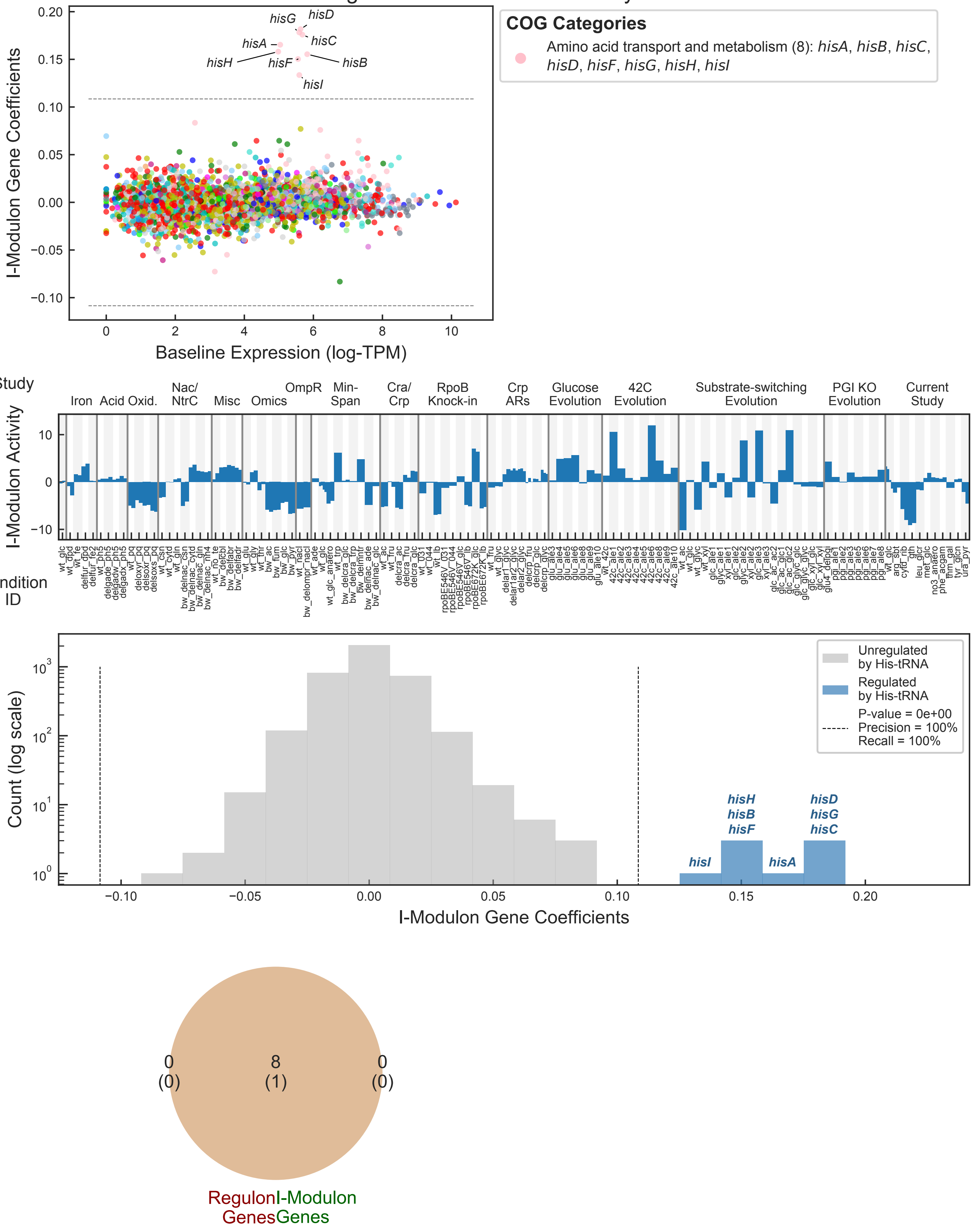

### Leu/Ile I-Modulon

Regulated by: IlvY or leu-tRNA attenuation or ile-tRNA attenuation
Biological Function: Branched-chain amino acid biosynthesis

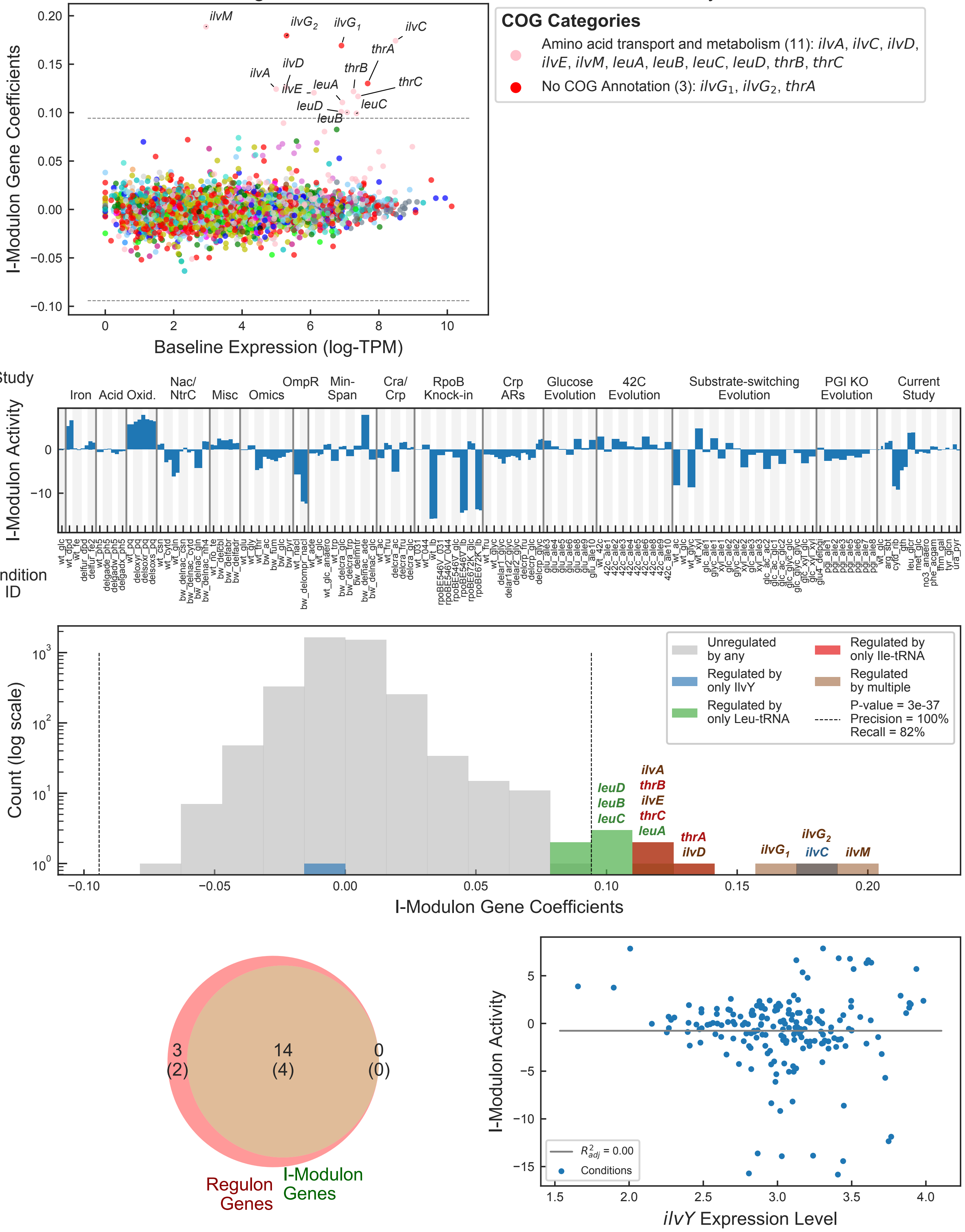

### MetJ I-Modulon

Regulated by: MetJ

Biological Function: Methionine biosynthesis

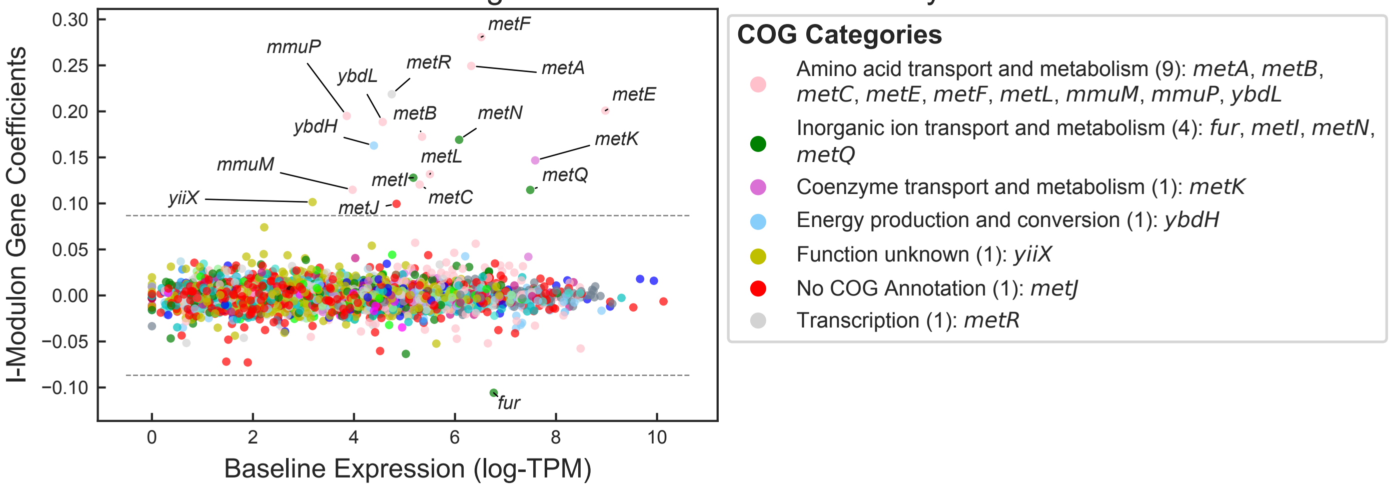


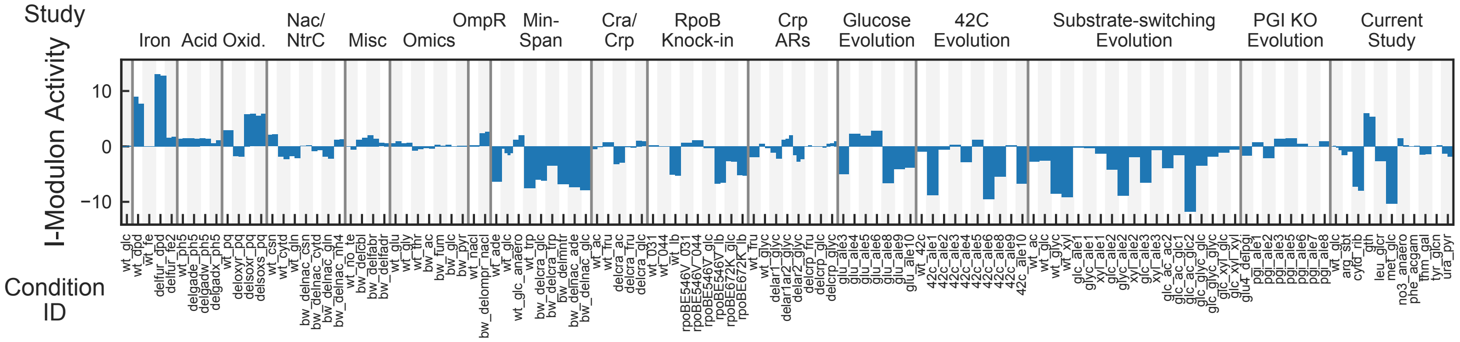


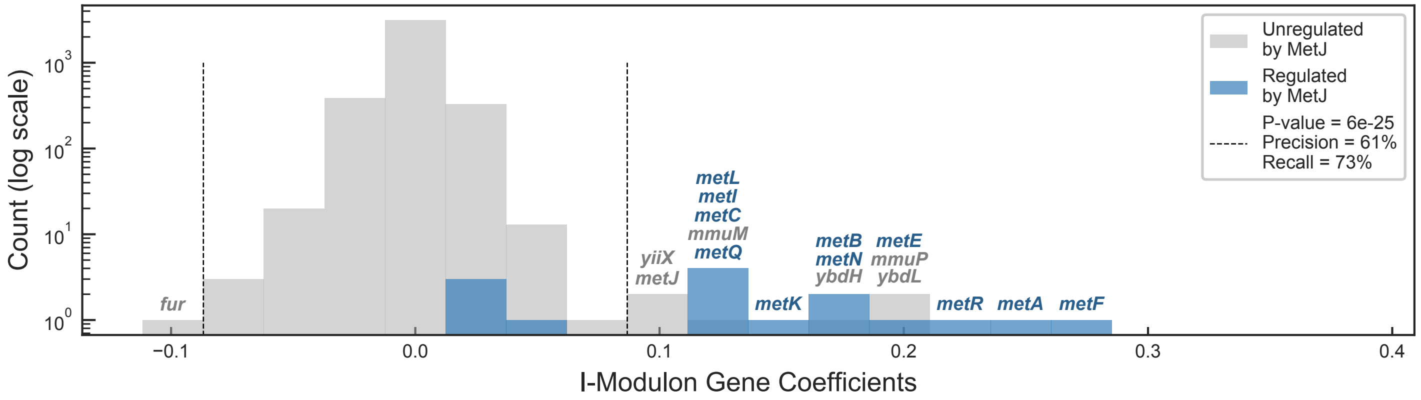


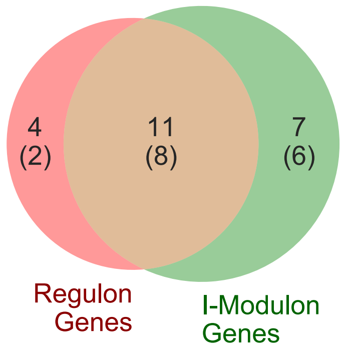


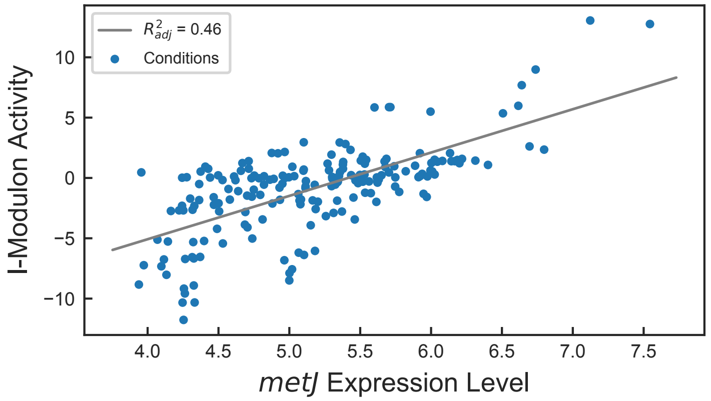


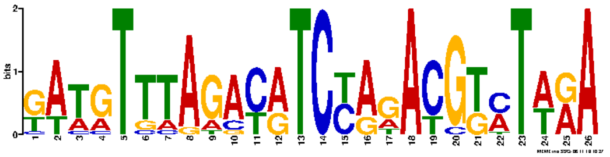

Motif E-value: 5.90e-37

Operons with Upstream Motif: 85%

### NagC/TyrR I-Modulon

Regulated by: NagC or TyrR

Biological Function: N-acetylglucosamine catabolism and tyrosine biosynthesis

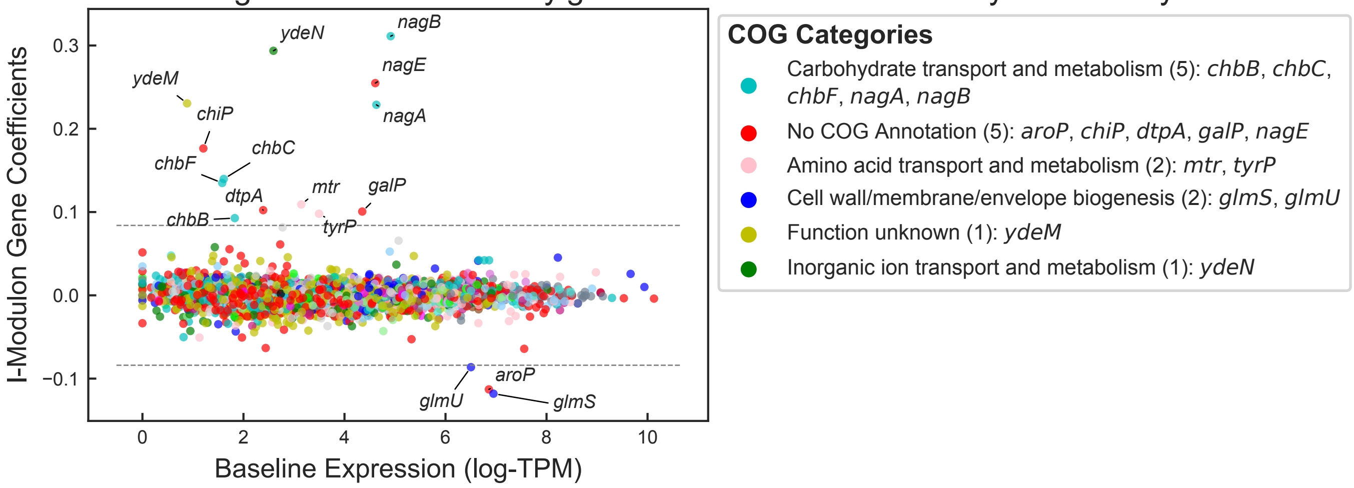


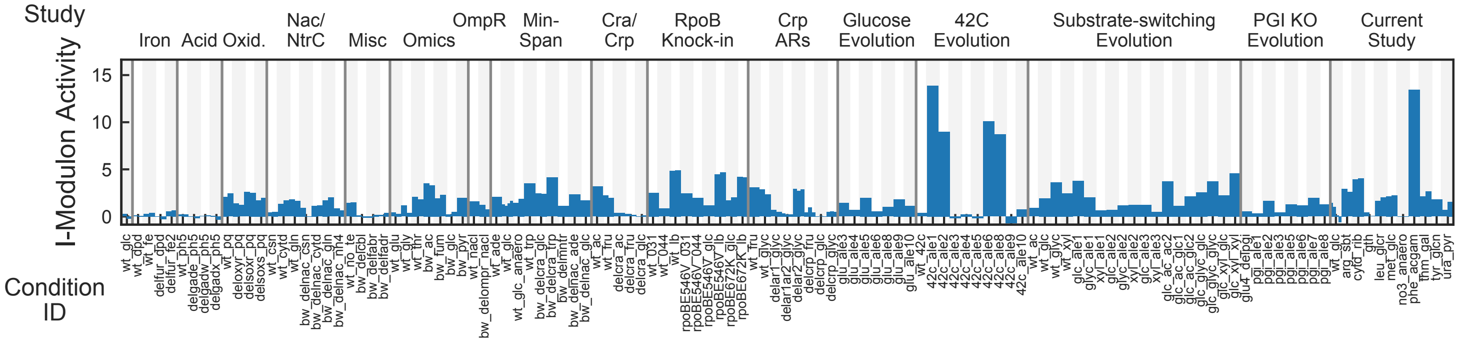


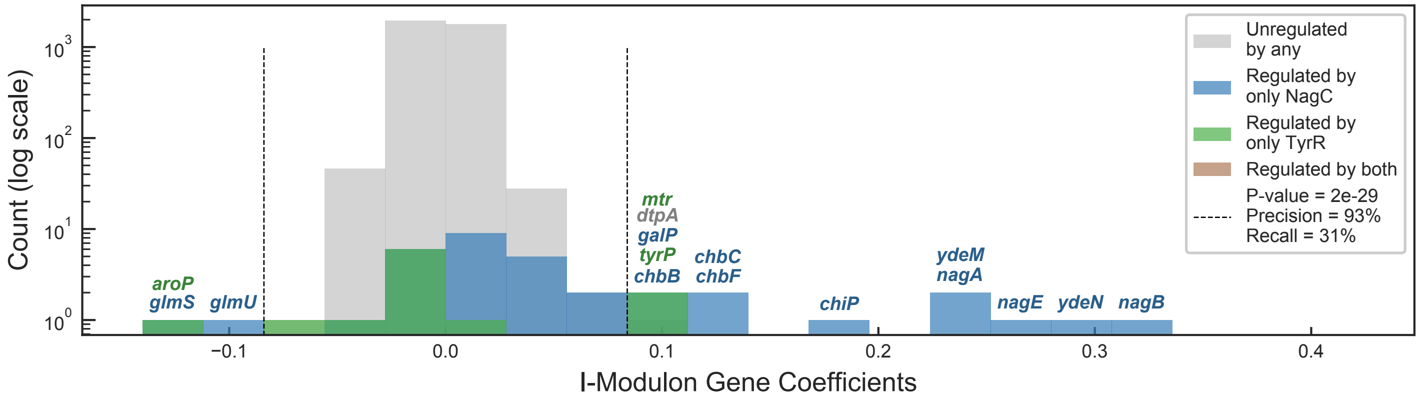


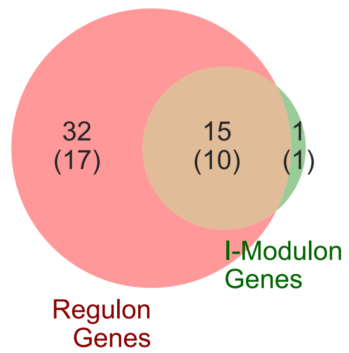


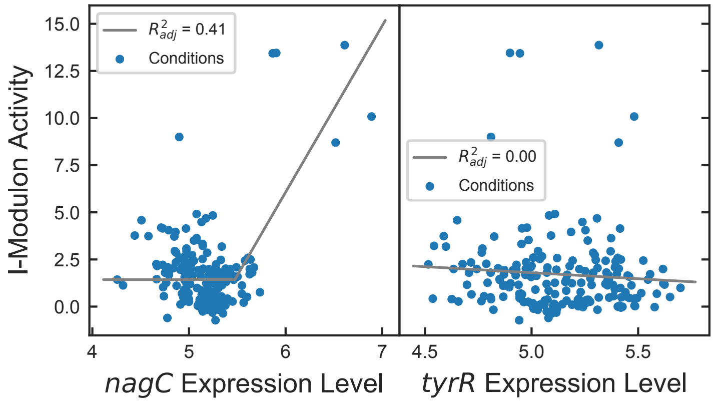


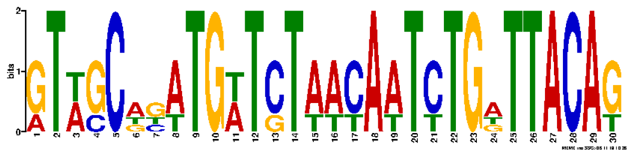

Motif E-value: 1.40e-04

Operons with Upstream Motif: 36%

### NarL I-Modulon

Regulated by: NarL

Biological Function: Nitrate respiration and formate dehydrogenase

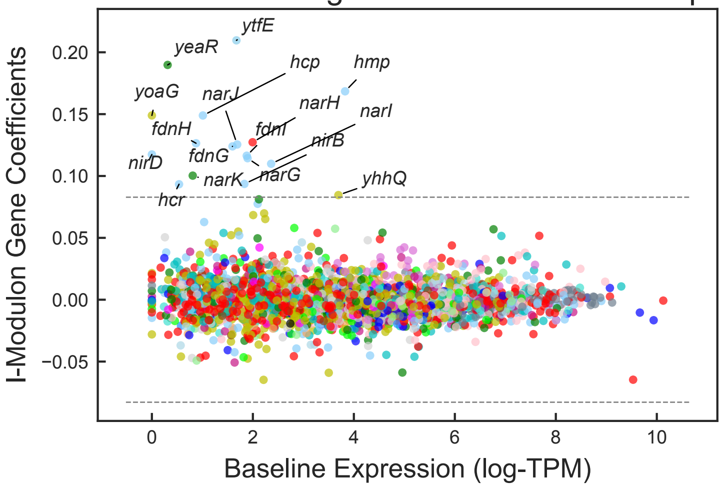


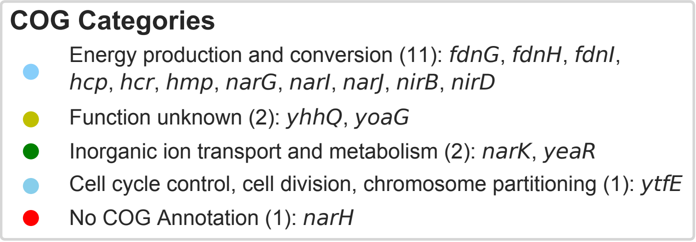


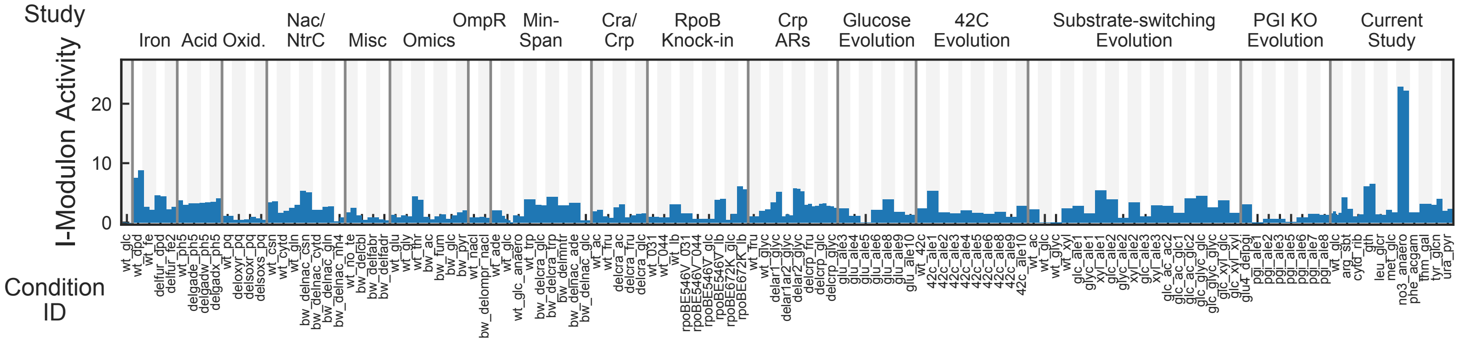


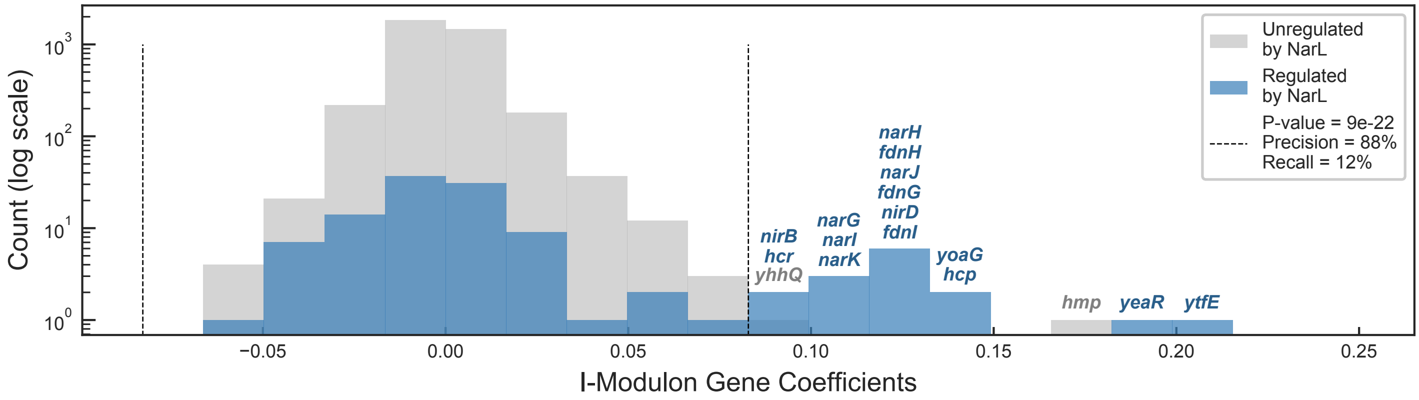


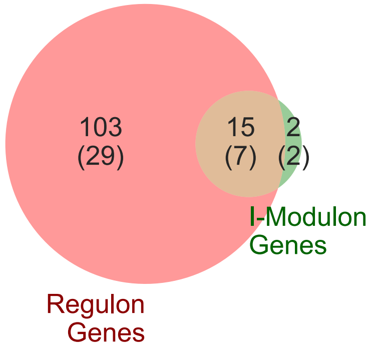


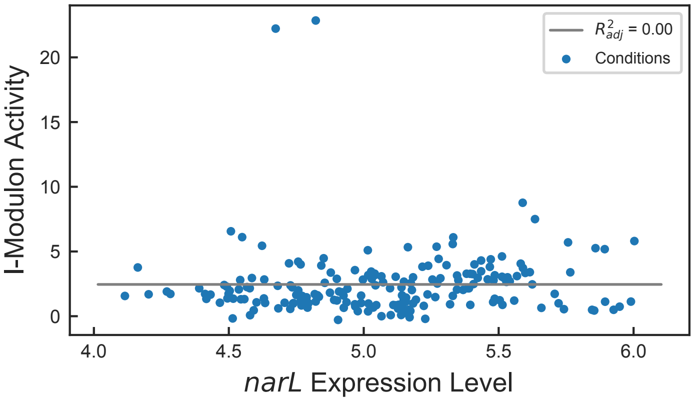


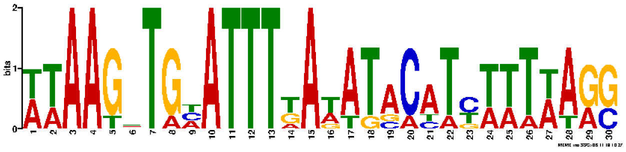

Motif E-value: 1.50e-05

Operons with Upstream Motif: 66%

### NtrC + RpoN I-Modulon

Regulated by: NtrC and RpoN

Biological Function: Nitrogen starvation response

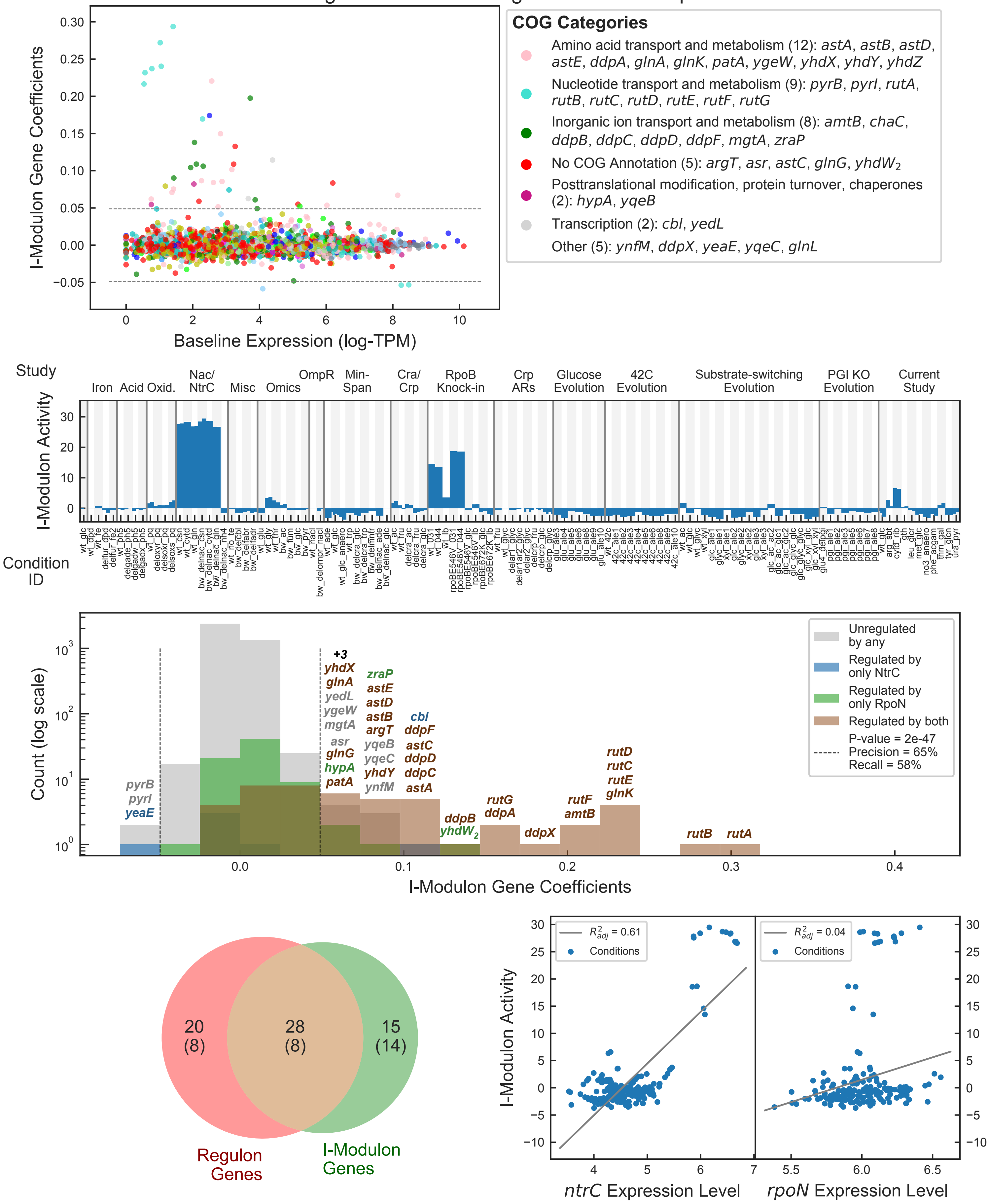

### PurR – 1 I-Modulon

Regulated by: PurR

Biological Function: Purine Biosynthesis

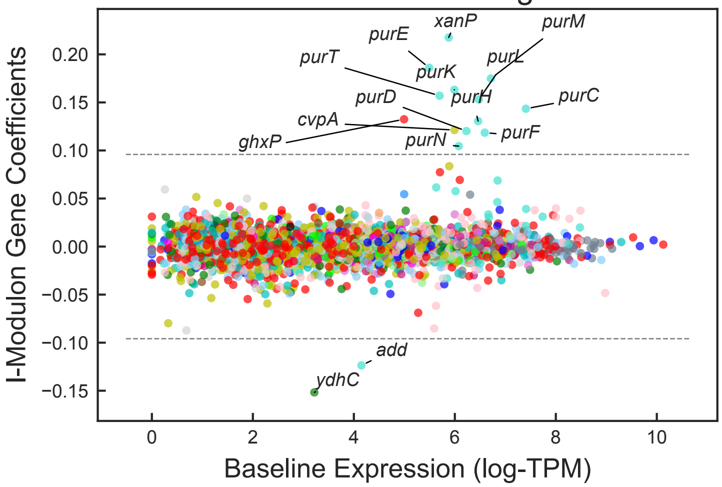


**COG Categories**

- Nucleotide transport and metabolism (12): *add*, *purC*, *purD*, *purE*, *purF*, *purH*, *purK*, *purL*, *purM*, *purN*, *purT*, *xanP*
- Function unknown (1): *cvpA*
- Inorganic ion transport and metabolism (1): *ydhC*
- No COG Annotation (1): *ghxP*

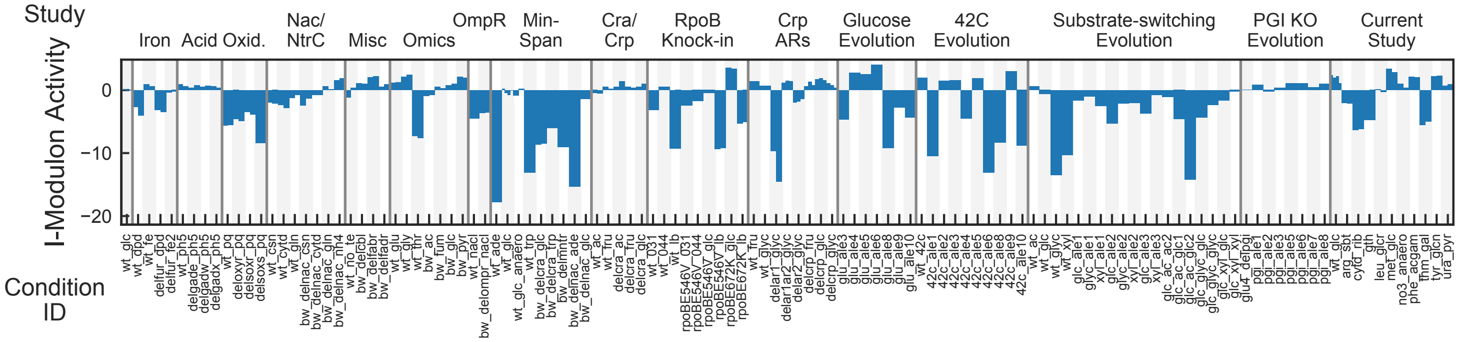


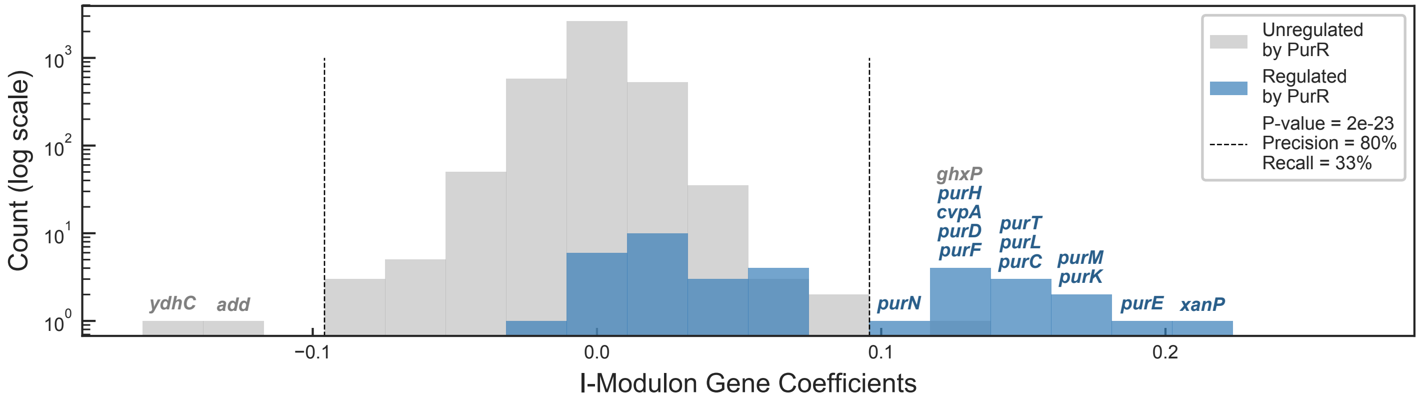


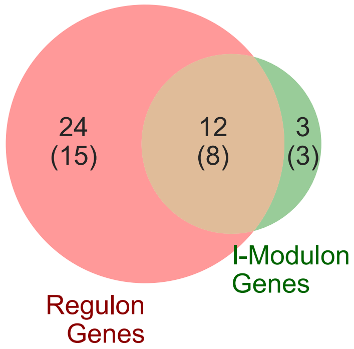


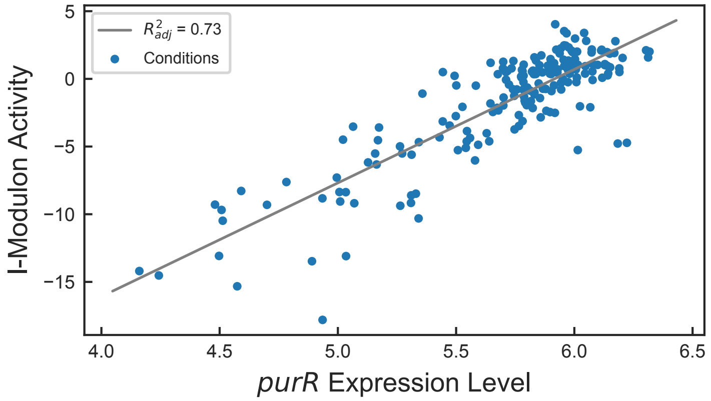


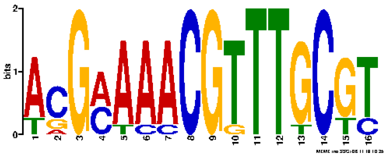

Motif E-value: 4.10e-19

Operons with Upstream Motif: 81%

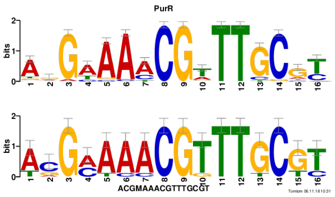


Motif similar to PurR (E-value: 7.38e-12)

### PurR−2 I-Modulon

Regulated by: PurR

Biological Function: Pyrimidine biosynthesis

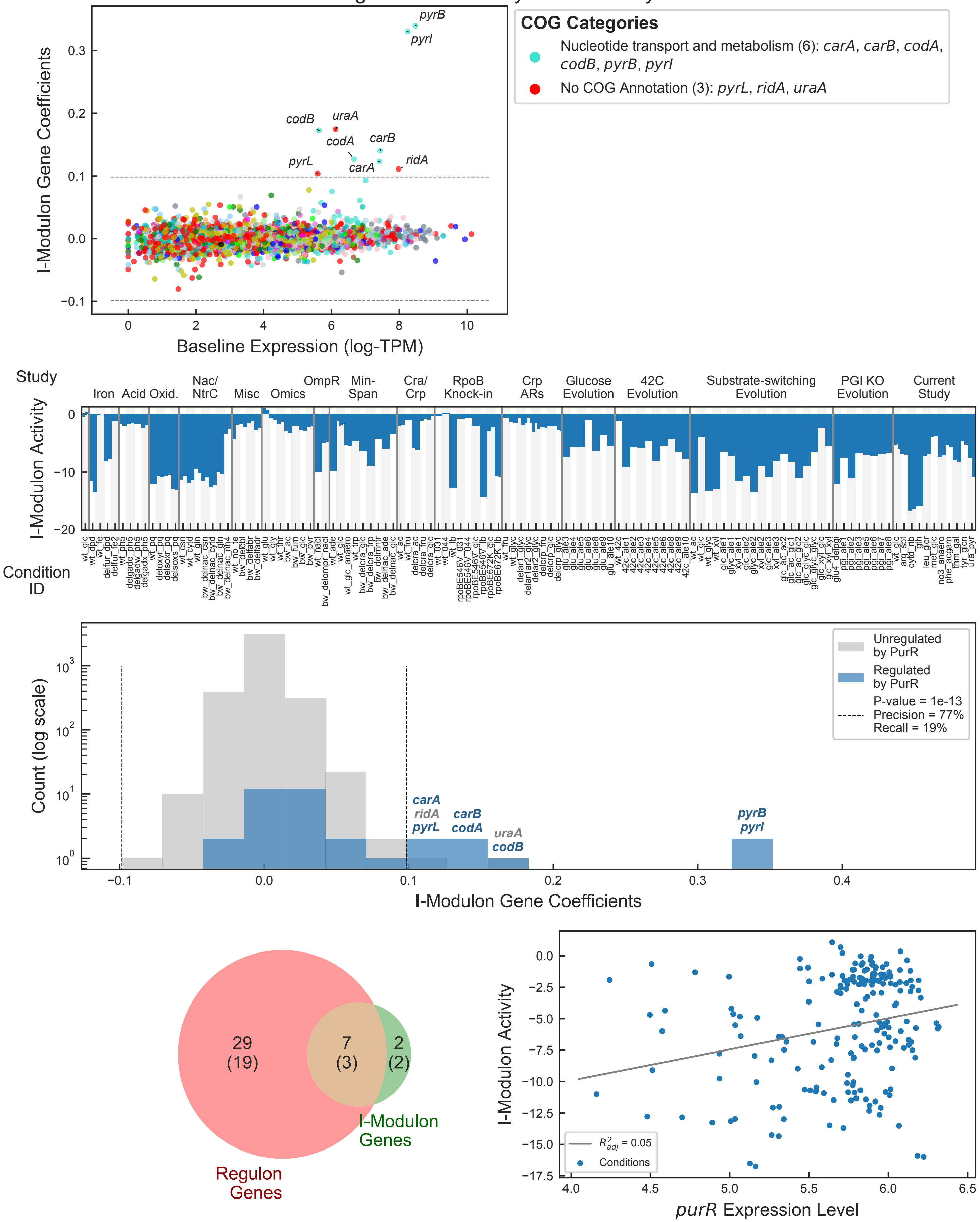

### Pyruvate I-Modulon

Regulated by: PdhR or YheO or YpdB or BtsR

Biological Function: Pyruvate transport and fermentation

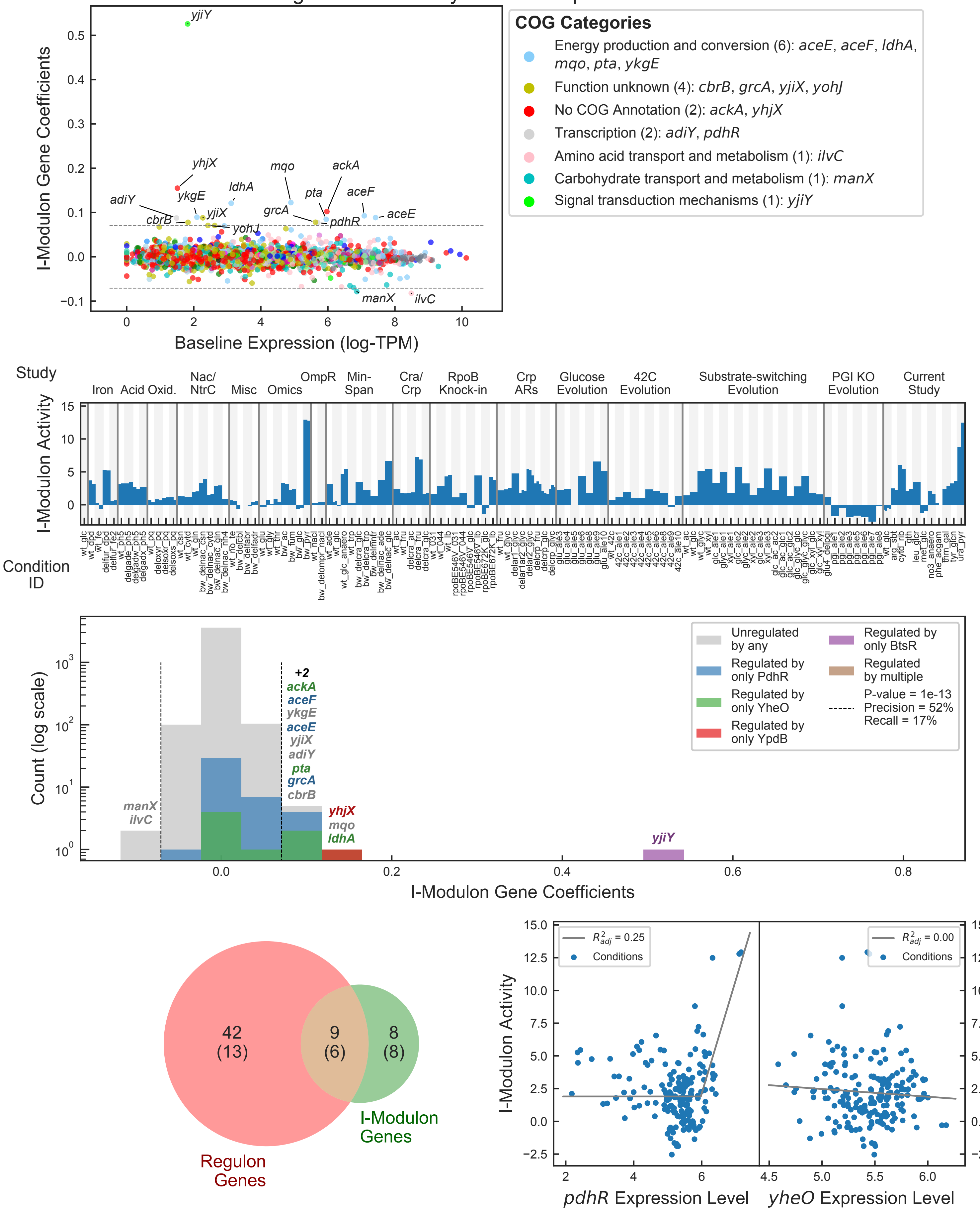

### RpoH I-Modulon

Regulated by: RpoH

Biological Function: Heat shock response

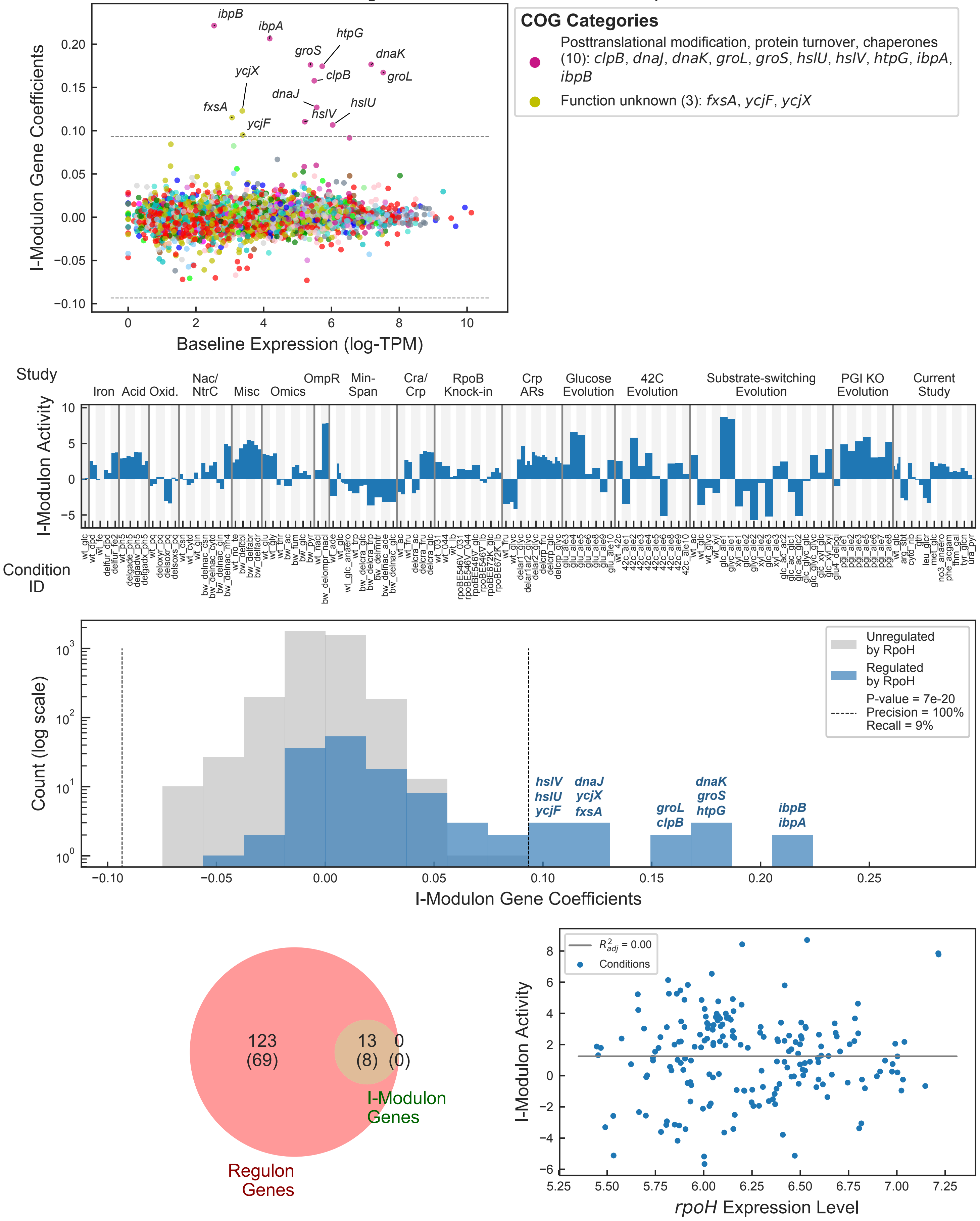

### RpoS I-Modulon

Regulated by: RpoS

Biological Function: General stress response

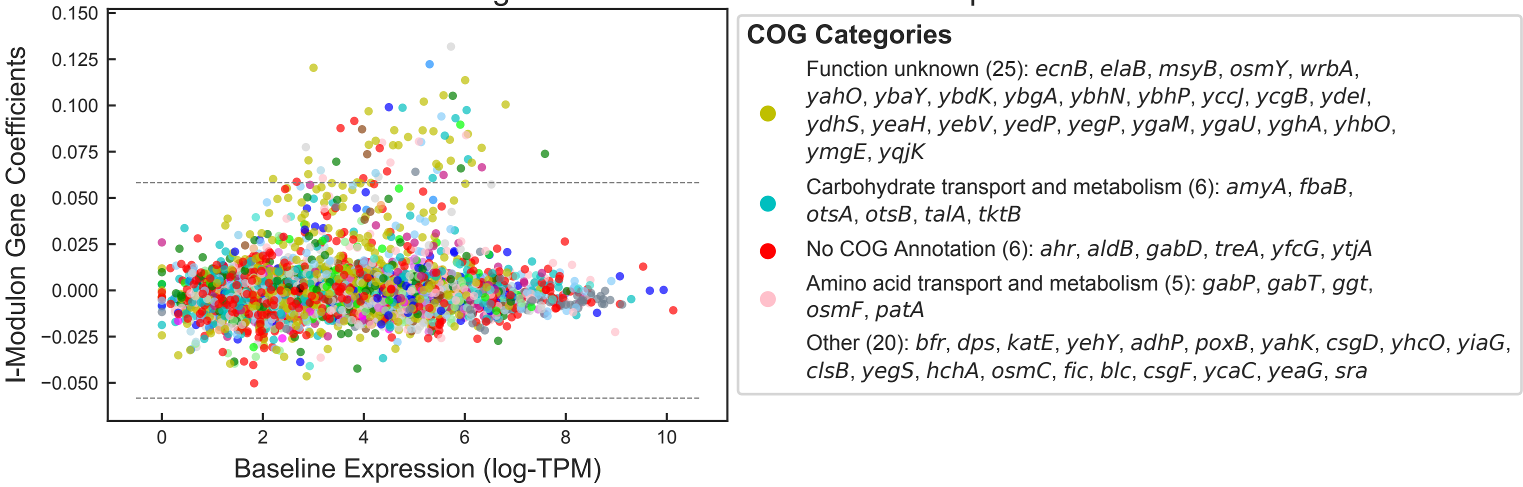


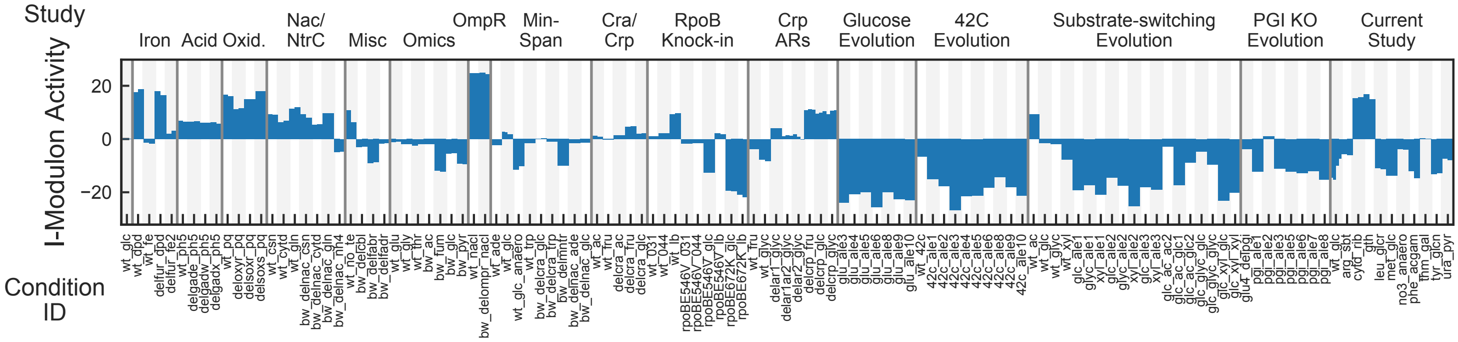


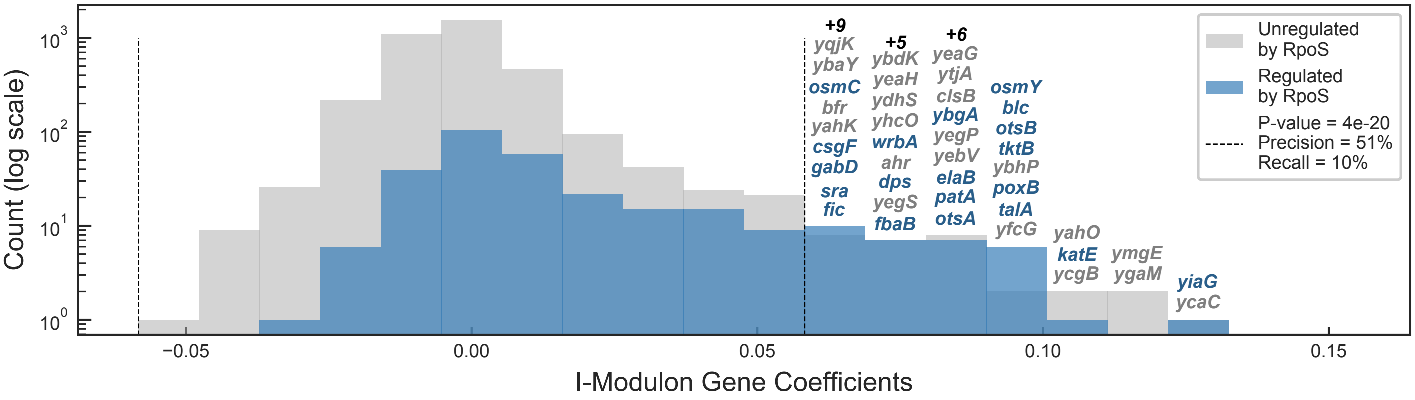


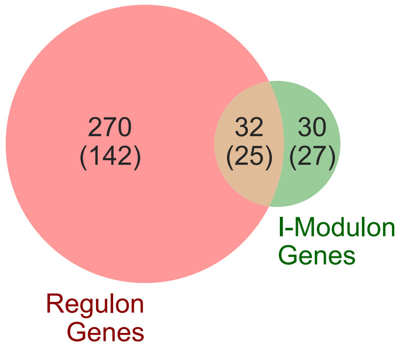


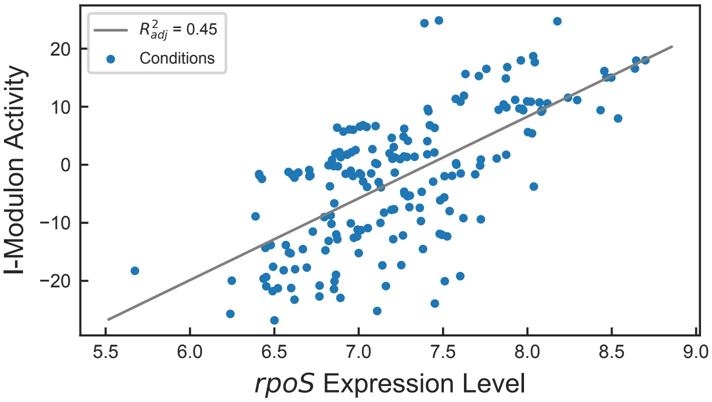


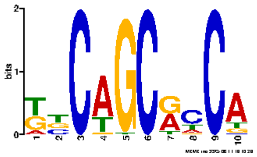

Motif E-value: 3.50e-05

Operons with Upstream Motif: 94%

### Thiamine I-Modulon

Regulated by: Thiamine

Biological Function: Thiamine biosynthesis

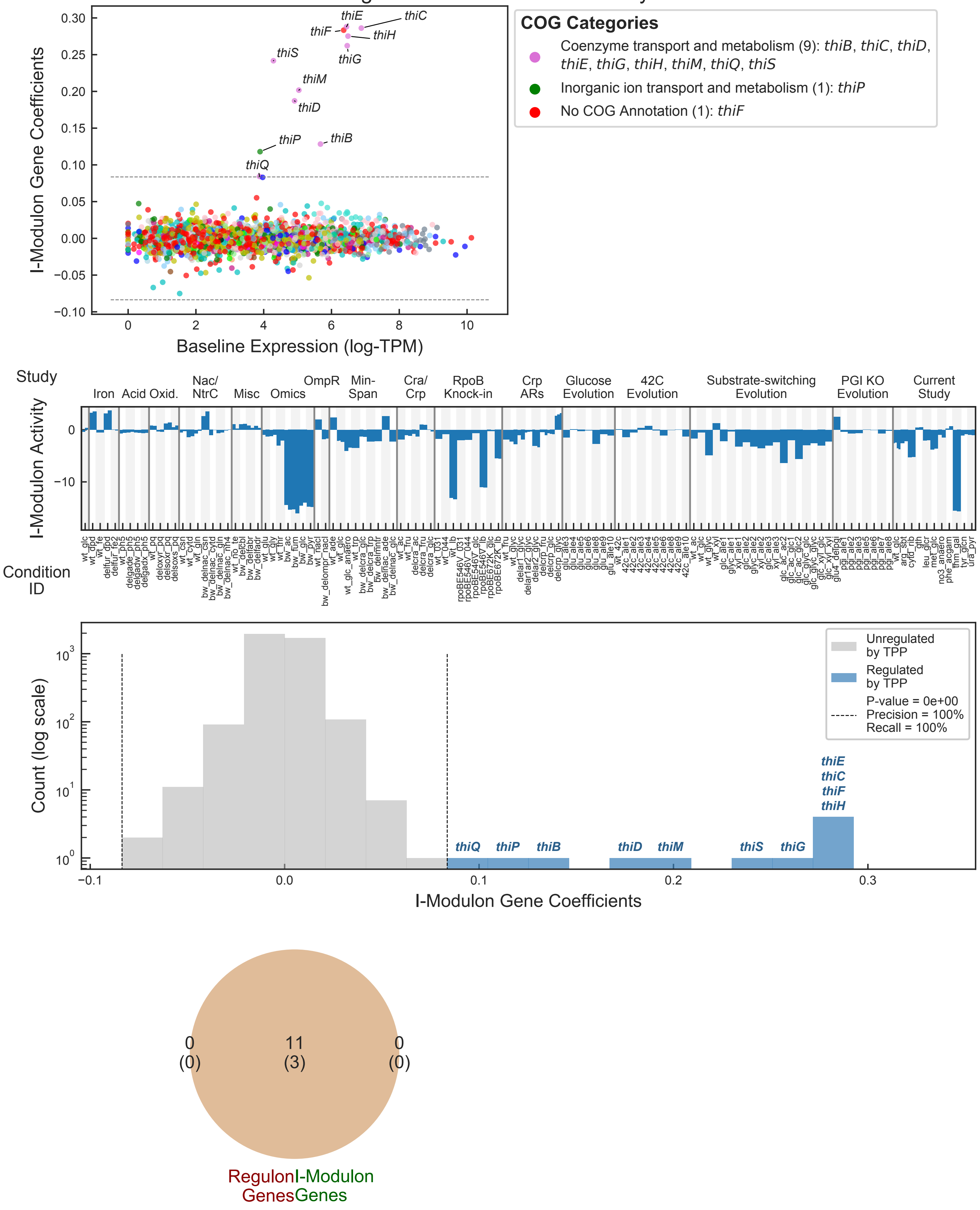

### Tryptophan I-Modulon

Regulated by: TrpR or trp-tRNA attenuation or Tryptophan attenuation
Biological Function: Tryptophan Biosynthesis

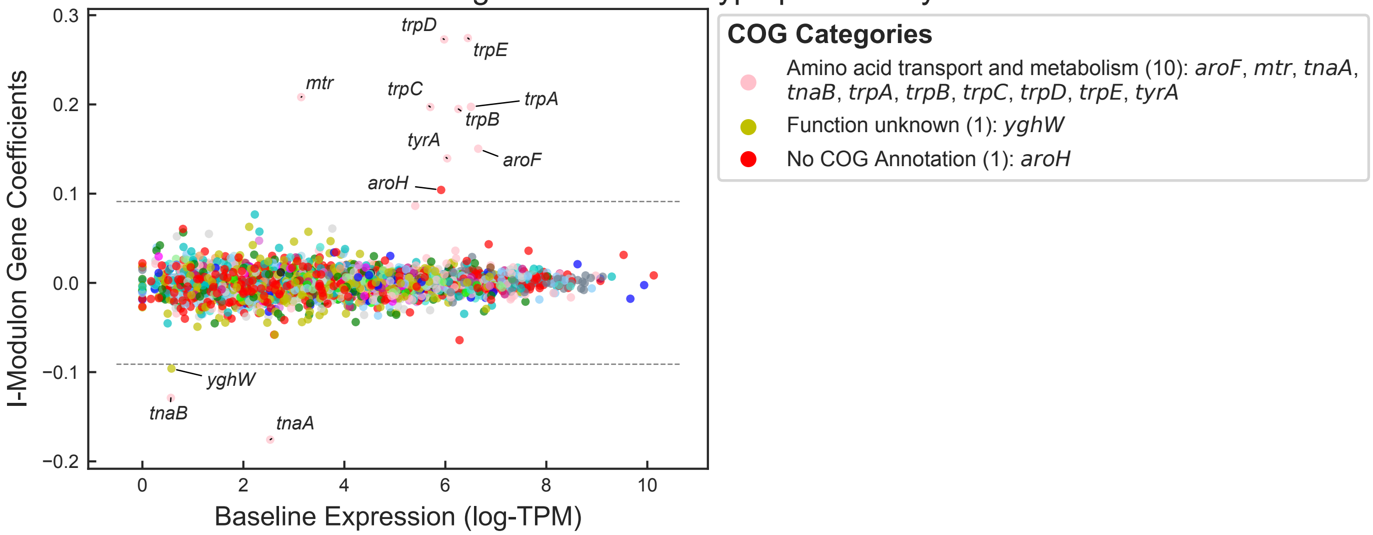


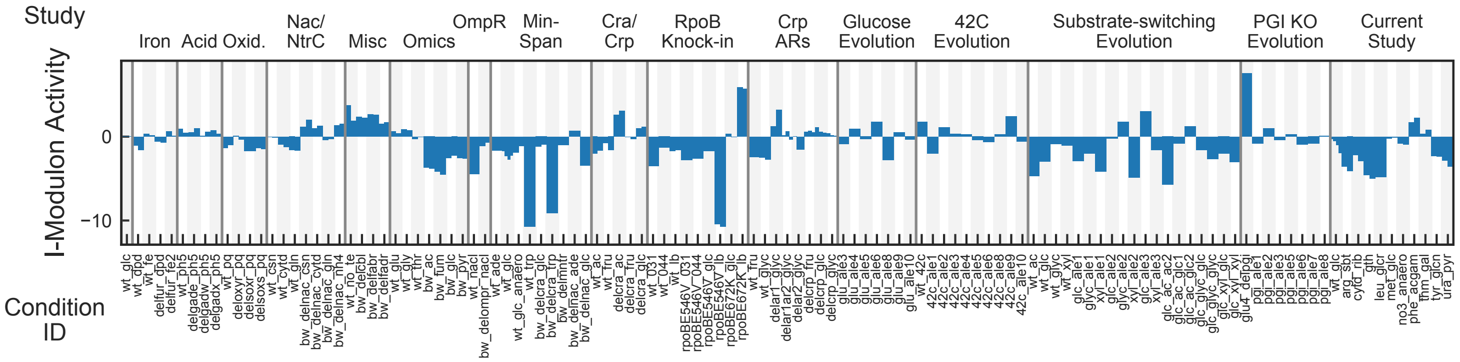


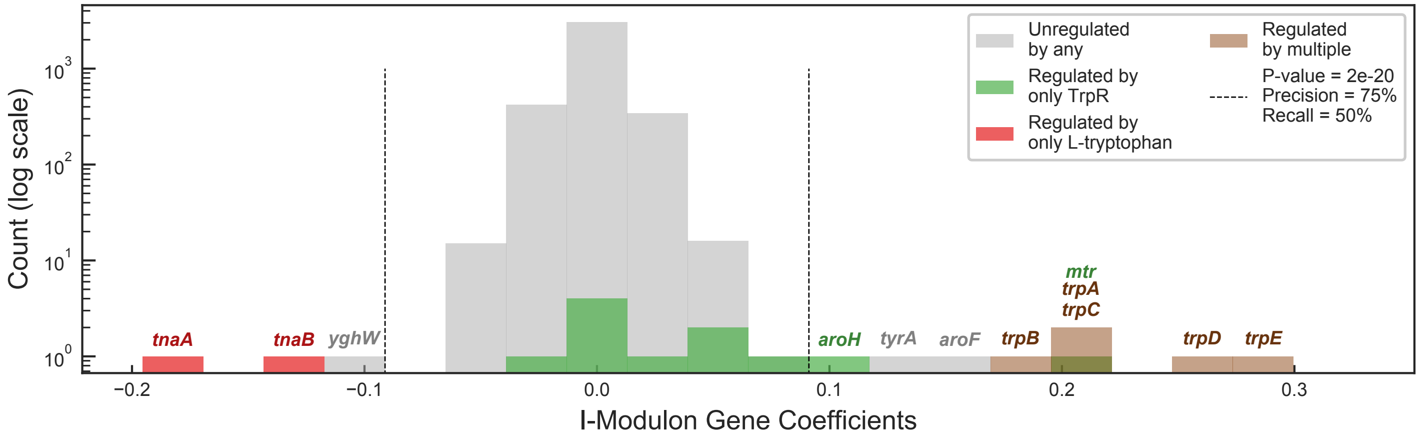


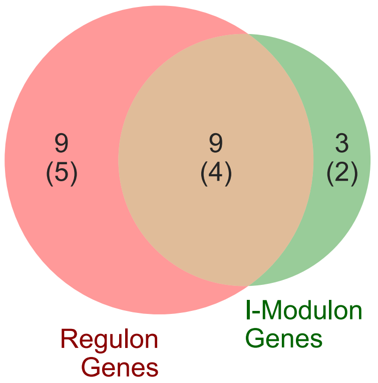


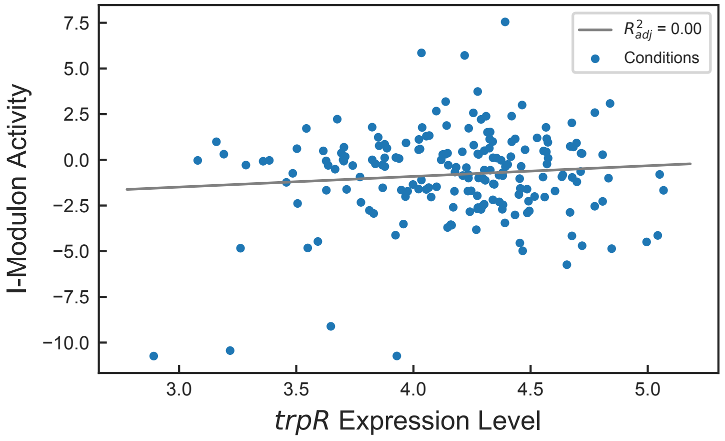

### translation I-Modulon

Biological Function: Enriched in translation machinery

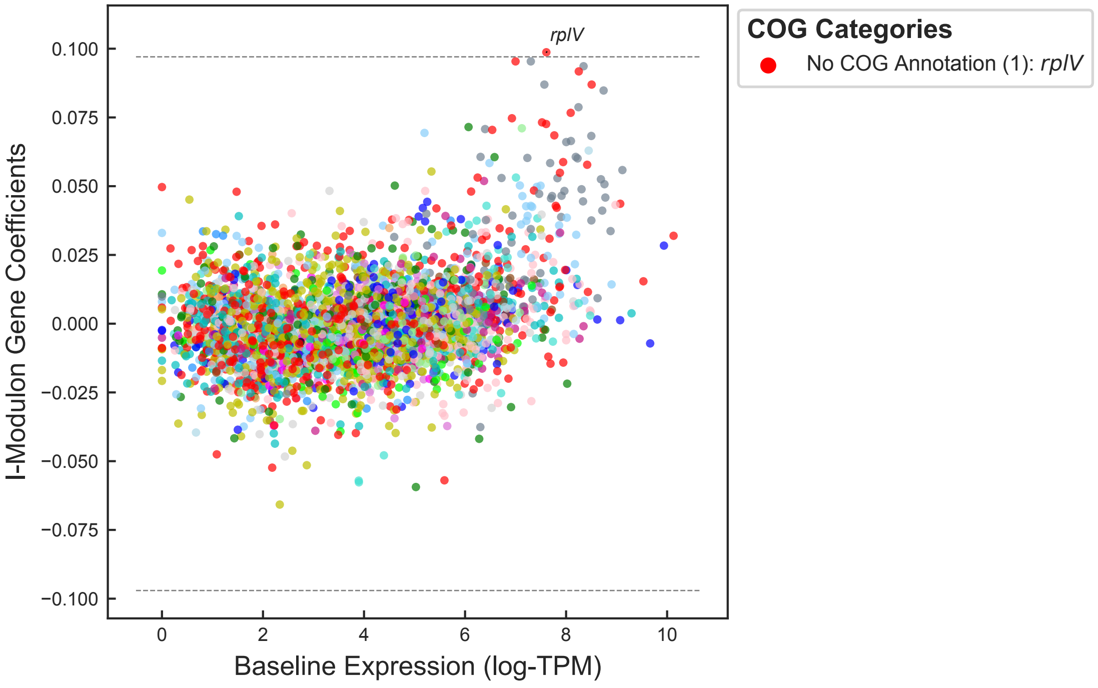


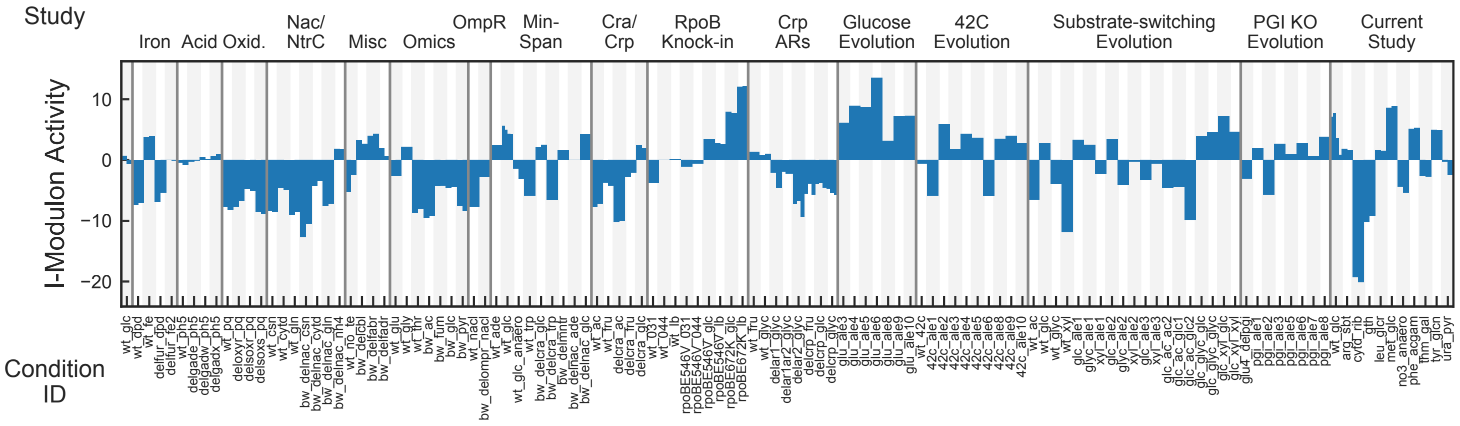


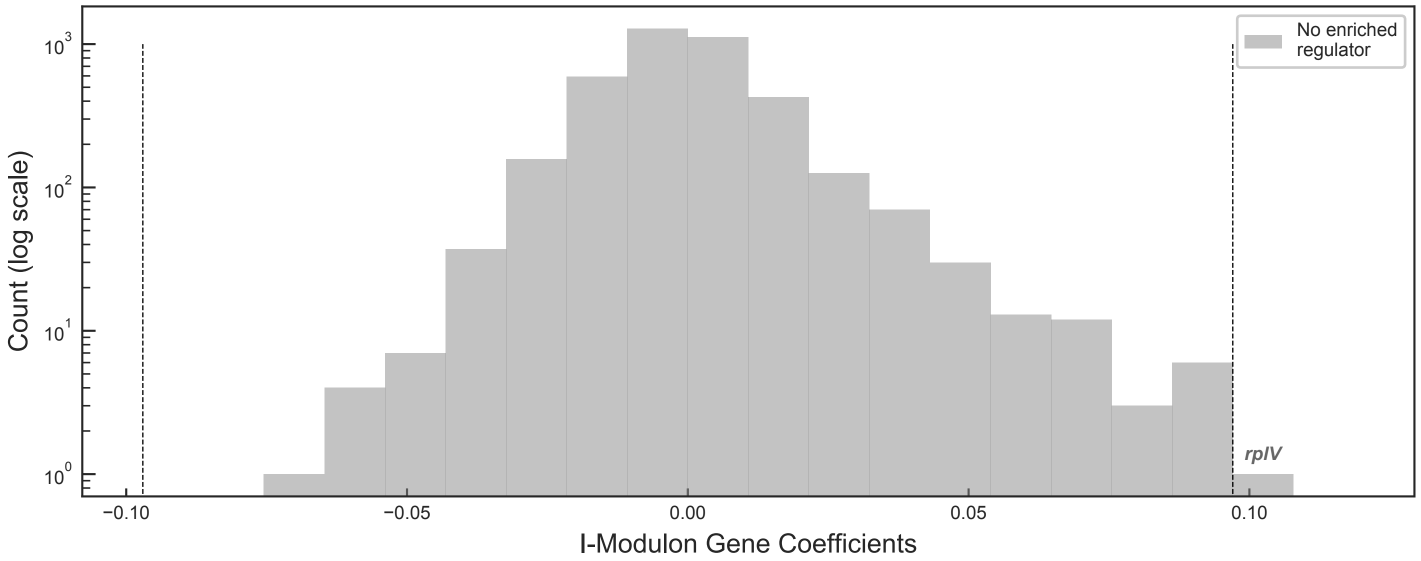
